# Genome-wide association study identifies 32 novel breast cancer susceptibility loci from overall and subtype-specific analyses

**DOI:** 10.1101/778605

**Authors:** Haoyu Zhang, Thomas U. Ahearn, Julie Lecarpentier, Daniel Barnes, Jonathan Beesley, Xia Jiang, Tracy A. O’Mara, Guanghao Qi, Ni Zhao, Manjeet K. Bolla, Alison M. Dunning, Joe Dennis, Qin Wang, Zumuruda Abu Ful, Kristiina Aittomäki, Irene L. Andrulis, Hoda Anton-Culver, Volker Arndt, Kristan J. Aronson, Banu K. Arun, Paul L. Auer, Jacopo Azzollini, Daniel Barrowdale, Heiko Becher, Matthias W. Beckmann, Sabine Behrens, Javier Benitez, Marina Bermisheva, Katarzyna Bialkowska, Ana Blanco, Carl Blomqvist, Natalia V. Bogdanova, Stig E. Bojesen, Bernardo Bonanni, Davide Bondavalli, Ake Borg, Hiltrud Brauch, Hermann Brenner, Ignacio Briceno, Annegien Broeks, Sara Y. Brucker, Thomas Brüning, Barbara Burwinkel, Saundra S. Buys, Helen Byers, Trinidad Caldés, Maria A. Caligo, Mariarosaria Calvello, Daniele Campa, Jose E. Castelao, Jenny Chang-Claude, Stephen J. Chanock, Melissa Christiaens, Hans Christiansen, Wendy K. Chung, Kathleen B.M. Claes, Christine L. Clarke, Sten Cornelissen, Fergus J. Couch, Angela Cox, Simon S. Cross, Kamila Czene, Mary B. Daly, Peter Devilee, Orland Diez, Susan M. Domchek, Thilo Dörk, Miriam Dwek, Diana M. Eccles, Arif B. Ekici, D.Gareth Evans, Peter A. Fasching, Jonine Figueroa, Lenka Foretova, Florentia Fostira, Eitan Friedman, Debra Frost, Manuela Gago-Dominguez, Susan M. Gapstur, Judy Garber, José A. García-Sáenz, Mia M. Gaudet, Simon A. Gayther, Graham G. Giles, Andrew K. Godwin, Mark S. Goldberg, David E. Goldgar, Anna González-Neira, Mark H. Greene, Jacek Gronwald, Pascal Guénel, Lothar Häberle, Eric Hahnen, Christopher A. Haiman, Christopher R. Hake, Per Hall, Ute Hamann, Elaine F. Harkness, Bernadette A.M. Heemskerk-Gerritsen, Peter Hillemanns, Frans B.L. Hogervorst, Bernd Holleczek, Antoinette Hollestelle, Maartje J. Hooning, Robert N. Hoover, John L. Hopper, Anthony Howell, Hanna Huebner, Peter J. Hulick, Evgeny N. Imyanitov, kConFab Investigators, ABCTB Investigators, Claudine Isaacs, Louise Izatt, Agnes Jager, Milena Jakimovska, Anna Jakubowska, Paul James, Ramunas Janavicius, Wolfgang Janni, Esther M. John, Michael E. Jones, Audrey Jung, Rudolf Kaaks, Pooja Middha Kapoor, Beth Y. Karlan, Renske Keeman, Sofia Khan, Elza Khusnutdinova, Cari M. Kitahara, Yon-Dschun Ko, Irene Konstantopoulou, Linetta B. Koppert, Stella Koutros, Vessela N. Kristensen, Anne-Vibeke Laenkholm, Diether Lambrechts, Susanna C. Larsson, Pierre Laurent-Puig, Conxi Lazaro, Emilija Lazarova, Flavio Lejbkowicz, Goska Leslie, Fabienne Lesueur, Annika Lindblom, Jolanta Lissowska, Wing-Yee Lo, Jennifer T. Loud, Jan Lubinski, Alicja Lukomska, Robert J. MacInnis, Arto Mannermaa, Mehdi Manoochehri, Siranoush Manoukian, Sara Margolin, Maria Elena Martinez, Laura Matricardi, Lesley McGuffog, Catriona McLean, Noura Mebirouk, Alfons Meindl, Usha Menon, Austin Miller, Elvira Mingazheva, Marco Montagna, Anna Marie Mulligan, Claire Mulot, Taru A. Muranen, Katherine L. Nathanson, Susan L. Neuhausen, Heli Nevanlinna, Patrick Neven, William G. Newman, Finn C. Nielsen, Liene Nikitina-Zake, Jesse Nodora, Kenneth Offit, Edith Olah, Olufunmilayo I. Olopade, Håkan Olsson, Nick Orr, Laura Papi, Janos Papp, Tjoung-Won Park-Simon, Michael T. Parsons, Bernard Peissel, Ana Peixoto, Beth Peshkin, Paolo Peterlongo, Julian Peto, Kelly-Anne Phillips, Marion Piedmonte, Dijana Plaseska-Karanfilska, Karolina Prajzendanc, Ross Prentice, Darya Prokofyeva, Brigitte Rack, Paolo Radice, Susan J. Ramus, Johanna Rantala, Muhammad U. Rashid, Gad Rennert, Hedy S. Rennert, Harvey A. Risch, Atocha Romero, Matti A. Rookus, Matthias Rübner, Thomas Rüdiger, Emmanouil Saloustros, Sarah Sampson, Dale P. Sandler, Elinor J. Sawyer, Maren T. Scheuner, Rita K. Schmutzler, Andreas Schneeweiss, Minouk J. Schoemaker, Ben Schöttker, Peter Schürmann, Leigha Senter, Priyanka Sharma, Mark E. Sherman, Xiao-Ou Shu, Christian F. Singer, Snezhana Smichkoska, Penny Soucy, Melissa C. Southey, John J. Spinelli, Jennifer Stone, Dominique Stoppa-Lyonnet, EMBRACE Study, GEMO Study Collaborators, Anthony J. Swerdlow, Csilla I. Szabo, Rulla M. Tamimi, William J. Tapper, Jack A. Taylor, Manuel R. Teixeira, MaryBeth Terry, Mads Thomassen, Darcy L. Thull, Marc Tischkowitz, Amanda E. Toland, Rob A.E.M. Tollenaar, Ian Tomlinson, Diana Torres, Melissa A. Troester, Thérèse Truong, Nadine Tung, Michael Untch, Celine M. Vachon, Ans M.W. van den Ouweland, Lizet E. van der Kolk, Elke M. van Veen, Elizabeth J. van Rensburg, Ana Vega, Barbara Wappenschmidt, Clarice R. Weinberg, Jeffrey N. Weitzel, Hans Wildiers, Robert Winqvist, Alicja Wolk, Xiaohong R. Yang, Drakoulis Yannoukakos, Wei Zheng, Kristin K. Zorn, Monica Zuradelli, Roger L. Milne, Peter Kraft, Jacques Simard, Paul D.P. Pharoah, Kyriaki Michailidou, Antonis C. Antoniou, Marjanka K. Schmidt, Georgia Chenevix-Trench, Douglas F. Easton, Nilanjan Chatterjee, Montserrat García-Closas

## Abstract

Breast cancer susceptibility variants frequently show heterogeneity in associations by tumor subtype. To identify novel loci, we performed a genome-wide association study (GWAS) including 133,384 breast cancer cases and 113,789 controls, plus 18,908 *BRCA1* mutation carriers (9,414 with breast cancer) of European ancestry, using both standard and novel methodologies that account for underlying tumor heterogeneity by estrogen receptor (ER), progesterone receptor (PR), and human epidermal growth factor receptor 2 (HER2) status and tumor grade. We identified 32 novel susceptibility loci (*P*<5.0×10^-8^), 15 of which showed evidence for associations with at least one tumor feature (false discovery rate <0.05). Five loci showed associations (*P*<0.05) in opposite directions between luminal- and non-luminal subtypes. *In-silico* analyses showed these five loci contained cell-specific enhancers that differed between normal luminal and basal mammary cells. The genetic correlations between five intrinsic-like subtypes ranged from 0.35 to 0.80. The proportion of genome-wide chip heritability explained by all known susceptibility loci was 37.6% for triple-negative and 54.2% for luminal A-like disease. These findings provide an improved understanding of genetic predisposition to breast cancer subtypes and will inform the development of subtype-specific polygenic risk scores.

---

GWAS have identified over 170 independent breast cancer susceptibility loci, many of which show differential associations by tumor subtypes, particularly ER-positive versus ER-negative or triple negative (TN) disease^1–3^. However, prior GWAS have not simultaneously investigated multiple, correlated tumor markers to identify additional source(s) of etiologic heterogeneity. We performed a breast cancer GWAS using both standard analyses and a novel two-stage polytomous regression method that efficiently characterizes etiologic heterogeneity while accounting for tumor marker correlations and missing data^4^.

The study populations and genotyping are described elsewhere^1, 2, 5, 6^ and in the **Online Methods**. Briefly, we analyzed data from 118,474 cases and 96,201 controls of European ancestry participating in 82 studies from the Breast Cancer Association Consortium (BCAC) and 9,414 affected and 9,494 unaffected *BRCA1* mutation carriers from 60 studies from the Consortium of Investigators of Modifiers of *BRCA1/2* (CIMBA) with genotyping data from one of two Illumina genome-wide custom arrays. In analyses of overall breast cancer, we also included summary level data from 11 other breast cancer GWAS (14,910 cases and 17,588 controls) without subtype information. Our study expands upon previous BCAC GWAS^1^ with additional data on 10,407 cases and 7,815 controls, an approximate increase of 10% and 9%, respecitvely. (**Supplementary Tables 1-4**).

The statistical methods are further described in the **Online Methods** and in **Supplementary Figure 1**. To identify single nucleotide polymorphisms (SNPs) for overall breast cancer (invasive, *in situ* or unknown invasiveness) in BCAC, we used standard logistic regression to estimate odds ratios (OR) and 95% confidence-intervals (CI) adjusting for country and principal components (PCs). iCOGS and OncoArray data were evaluated separately and results combined with those from the 11 other GWAS using fixed-effects meta-analysis.

To identify invasive breast cancer susceptibility SNPs displaying evidence of heterogeneity, we used a novel score-test based on a two-stage polytomous model^4^ that allows flexible, yet parsimonious, modelling of associations in the presence of underlying heterogeneity by ER, PR, HER2 and/or grade (**Online Methods, Supplementary Note**). The model handles missing tumor characteristic data by implementing an efficient Expectation-Maximization algorithm^4, 7^. These analyses were restricted to BCAC controls and invasive cases (**Online Methods**). We fit an additional two-stage model to estimate case-control ORs and 95% CI between the SNPs and intrinsic-like subtypes defined by combinations of ER, PR, HER2 and grade (**Online Methods**): (1) luminal A-like, (2) luminal B/HER2-negative-like, (3) luminal B-like, (4) HER2-enriched-like and (5) TN or basal-like. We analyzed iCOGS and OncoArray data separately, adjusting for PCs and age, and meta-analyzed the results using a fixed-effects model. We evaluated the effect of country using a leave-one-out sensitivity analysis (**Online Methods**).

We used data from the *BRCA1* mutation carriers who are prone to develop TN disease^8^, to estimate per-allele hazard ratios (HRs) within a retrospective cohort analysis framework. We assumed the estimated ORs for BCAC TN cases and the HRs estimated from CIMBA *BRCA1* carriers approximated the same underlying relative risk^8^, and used a fixed-effect meta-analysis to combine these risk estimates (**Online Methods**). We used the two-stage polytomous model to test for heterogeneity in associations for all newly identified SNPs across subtypes, globally and by tumor-specific markers (**Online Methods**).

Overall, we identified 32 novel independent susceptibility loci marked by SNPs with *P*<5.0×10^-8^ (Figure 1, **Supplementary Table 5-7, Supplementary Figure 2-6**): 22 SNPs using standard logistic regression, eight SNPs using the two-stage polytomous model and three SNPs in the CIMBA/BCAC-TN meta-analysis (rs78378222 was also detected by the two-stage polytomous model in BCAC). Fourteen additional significant (P<5.0×10^-8^) SNPs were excluded, 13 because they lacked evidence of association independent of previously reported susceptibility SNPs in conditional analyses (*P* ≥1.0×10^-6^; **Supplementary Table 8-10**), and one (chr22:40042814) for showing a high-degree of sensitivity to leave-one-out country analysis (**Supplementary Figure 7**). **Supplemental figures 8-9** show the associations between all 32 SNPs and the intrinsic-like subtypes.

**Figure 1.**
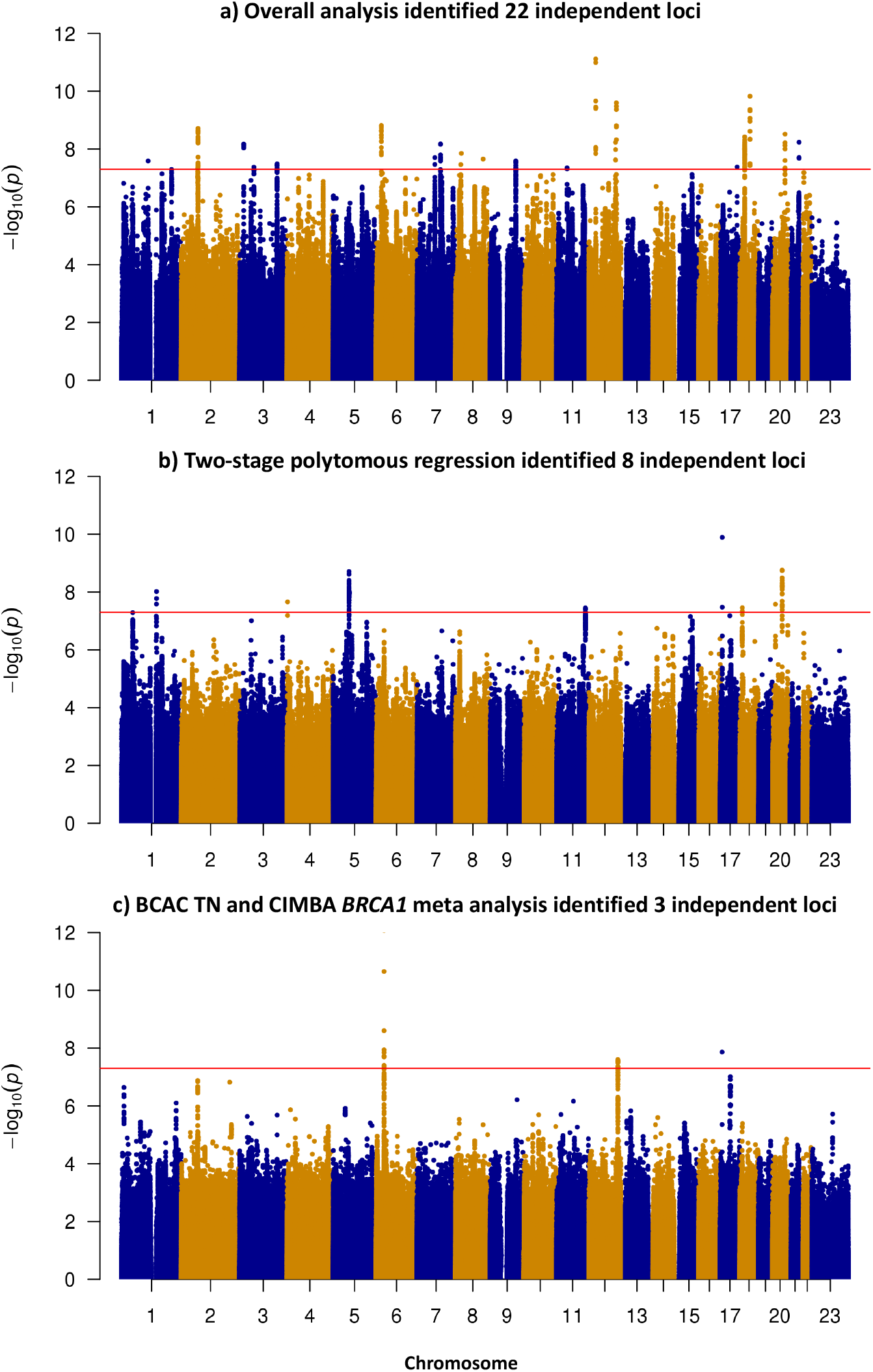
Manhattan plots showing 32 independent susceptibility loci crossing genome-wide significant threshold for associations with breast cancer risk. **a**) Twenty-two loci (**Supplementary Table 6**) were identified from standard logistic regression analysis using 133,384 breast cancer cases and 113,789 controls from BCAC; **b**) eight loci (**Supplementary Table 7**) identified from two-stage polytomous regression using 106,278 invasive cases and 91,477 controls from BCAC; **c**) three loci (**Supplementary Table 8**) were identified from meta-analysis of triple negative in BCAC and 18,908 female *BRCA1(*unaffected *BRCA1* mutation carriers = 9,494). SNP rs78378222 was also detected in two-stage polytomous regression model.

Fifteen of the 32 SNPs showed evidence of heterogeneity (FDR<0.05) according to the global heterogeneity test (Figure 2, **Supplementary Table 11**). Nine of these were identified in analyses accounting for tumor marker heterogeneity. ER (7 SNPs) and grade (7 SNPs) most often contributed to observed heterogeneity (marker-specific *P*<0.05), followed by HER2 (4 SNPs) and PR (2 SNPs). rs17215231, identified in the CIMBA/BCAC-TN meta-analysis, was the only SNP found exclusively associated with TN disease (OR=0.85, 95%CI=0.81-0.89; *P*=8.6×10^-13^). rs2464195, also identified in the CIMBA/BCAC-TN meta-analysis, was associated with both TN (OR=0.93, 95%CI=0.91-0.96; *P*=2.5×10^-8^) and luminal B-like subtypes (OR=0.96, 95%CI=0.92-0.99; *P*=0.02; **Supplementary Table 7**, **Supplementary Figure 9**). This SNP is in LD (r^2^=0.62) with rs7953249, which is differentially associated with risk of subtypes of ovarian cancer^9^. Five of the heterogeneous SNPs showed associations with luminal and non-luminal subtypes in opposite directions (Figure 3). For example, four SNPs were associated in opposite directions with luminal A-like and TN subtypes (respectively, for rs78378222 OR=1.13, 95%CI=1.05-1.20 vs OR=0.67, 95%CI=0.57-0.80; for rs206435 OR=1.03, 95%CI=1.01-1.05 vs OR=0.95, 95%CI=0.92-0.98; for rs141526427 OR=0.96, 95%CI=0.94-0.98 vs OR=1.04, 95%CI=1.01-1.08; and for rs6065254 OR=0.96, 95%CI=0.94-0.97 vs OR=1.04, 95%CI=1.01-1.07). The specific tumor-marker heterogeneity test showed rs78378222 associated with ER (*P_ER_*=7.0×10^-6^) and HER2 (*P_HER2_*=2.07×10^-4^), rs206435 associated with ER (*P_ER_*=2.8×10^-3^) and grade (*P_grade_*=2.8×10^-4^) and rs141526427 (*P_ER_*=1.3×10^-3^) and rs6065254 (*P_ER_*=4.3×10^-3^) associated with ER. rs7924772 showed opposite associations between HER2-negative and HER2-positive subtypes (*e.g.*, OR=1.04, 95%CI=1.03-1.06 for luminal A-like disease and OR=0.95, 95%CI=0.92-0.99 for luminal B-like disease) and, consistent with these findings, was exclusively associated with HER2 (*P_HER2_*=1.4×10^-6^; Figure 3). Notably, rs78378222 located in the 3’ UTR of *TP53* also showed opposite associations with high-grade serous cancers (OR=0.75, *P*=3.7×10^-4^) and low-grade serous cancers (OR=1.58, *P*=1.5×10^-4^; http://ocac.ccge.medschl.cam.ac.uk). Moreover, prior analyses did not find rs78378222 associated with risk of breast cancer, likely due to its opposite effects between subtypes^10^.

**Figure 2.**
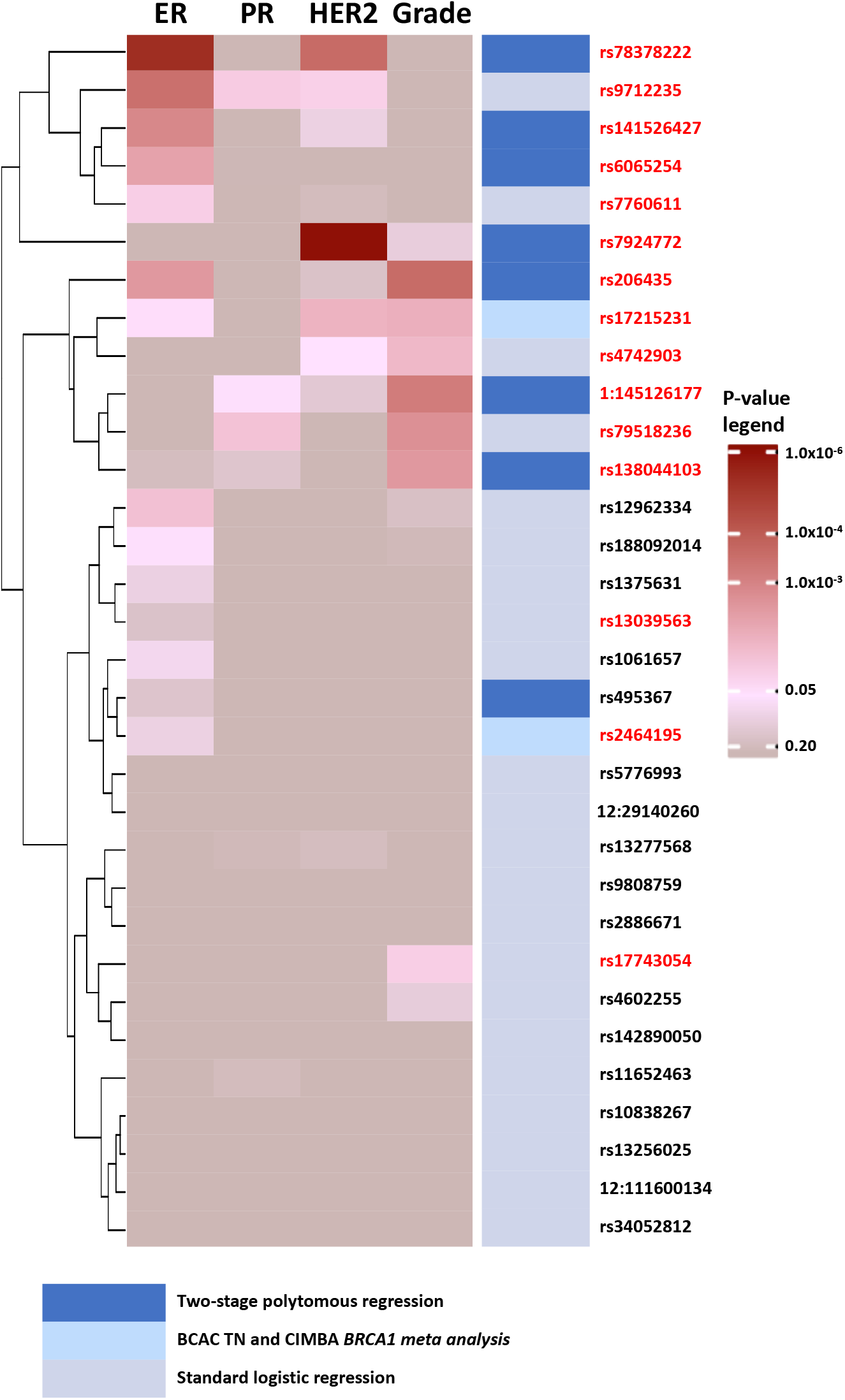
Heatmap and clustering of p-values from marker specific heterogeneity test for 32 breast cancer susceptibility loci. P-values are for associations between the most significant SNPs marking each loci and estrogen receptor (ER), progesterone receptor (PR), human epidermal growth factor receptor 2 (HER2) or grade, adjusting for top ten principal components and age. Fifteen SNPs in red color were significant according to the global heterogeneity tests (FDR <0.05) that evaluates if a SNP’s risk estimates vary by at least one of the four tumor characteristics.

**Figure 3.**
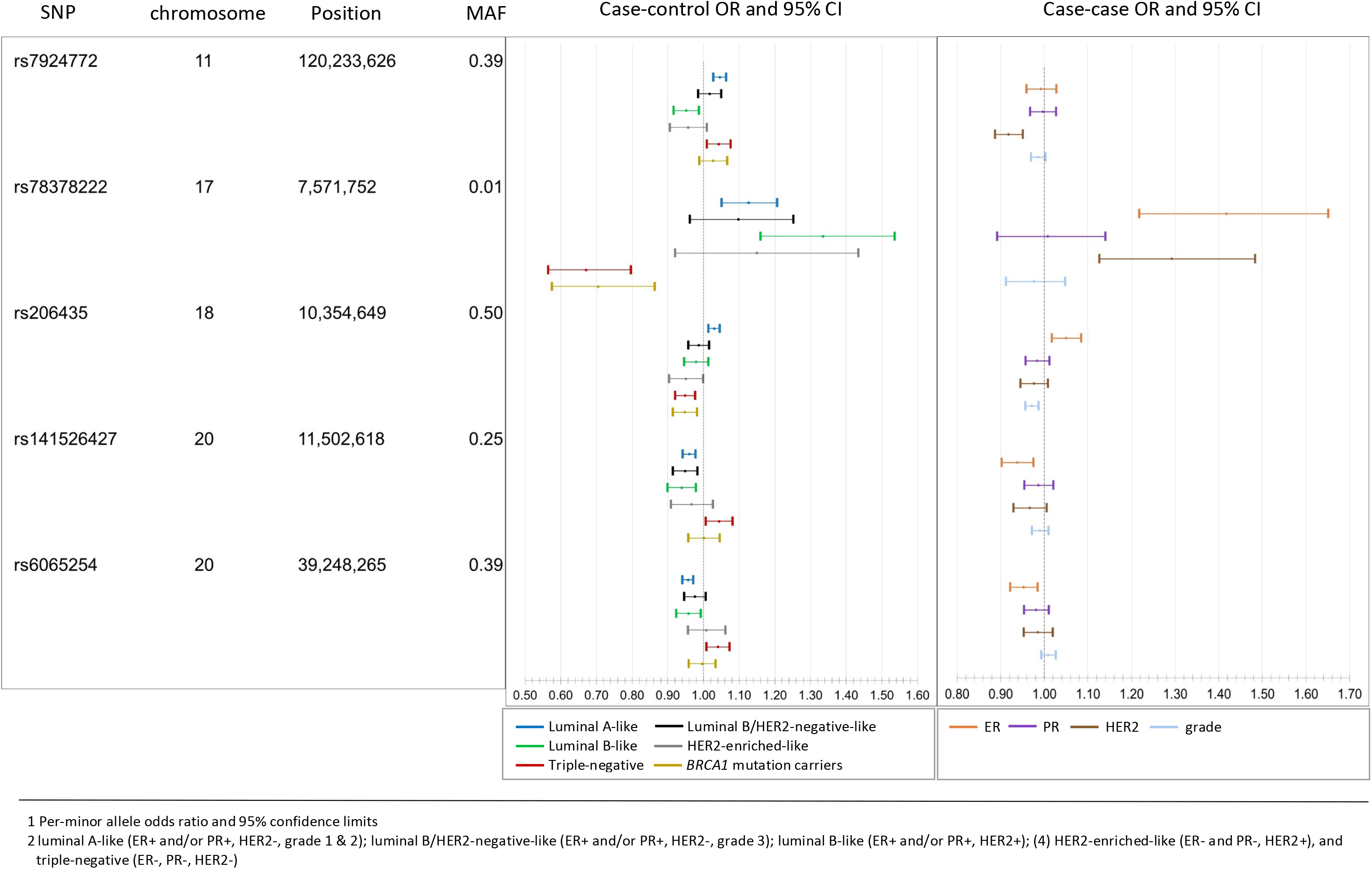
Susceptibility SNPs with associations in opposite direction across subtypes. The case-control odds ratios (OR) and 95% confidence intervals (95% CI)^1^ (left panel) are for associations of each of the five SNPs and risk for breast cancer intrinsic-like subtypes^2^ estimated from the first-stage of the two-stage polytomous regression fixed-effects model. The case-case ORs 95%CI (right panel) are estimated from the second stage parameters of a fixed effect two-stage polytomous models testing for heterogeneity between the five SNPs and estrogen receptor (ER), progesterone receptor (PR), human epidermal growth factor receptor 2 (HER2) and grade, where ER, PR, HER2, and grade are mutually adjusted for each other.

We defined a set of candidate causal variants (CCVs; **Online Methods**) for each novel locus and investigated the CCVs in relation to previously-annotated enhancers in primary breast cells^11^. Based on combinations of H3K4me1 and H3K27ac histone modification ChIP-seq signals, putative enhancers in basal cells (BC), luminal progenitor cells (LP) and mature luminal cells (LM) were characterized as “OFF,” “PRIMED”, and “ACTIVE” (**Online Methods**). We defined “ANYSWITCH” enhancers as those exhibiting different characterizations between cell types. Among the five loci showing evidence of having associations in opposite directions between some subtypes, at least one CCV per locus overlapped an “ANYSWITCH” enhancer (Figure 4). For example, rs78378222 overlapped an ACTIVE enhancer in BC, PRIMED in LP and OFF in LM. In comparison, 63% of the loci with consistent direction of associations across subtypes overlapped with an “ANYSWITCH” enhancer (**Supplementary Table 12-13**). These results support the hypothesis that some variants may modulate enhancer activity in a cell-type specific manner and thus differentially influence the risk of developing different tumor subtypes.

**Figure 4.**
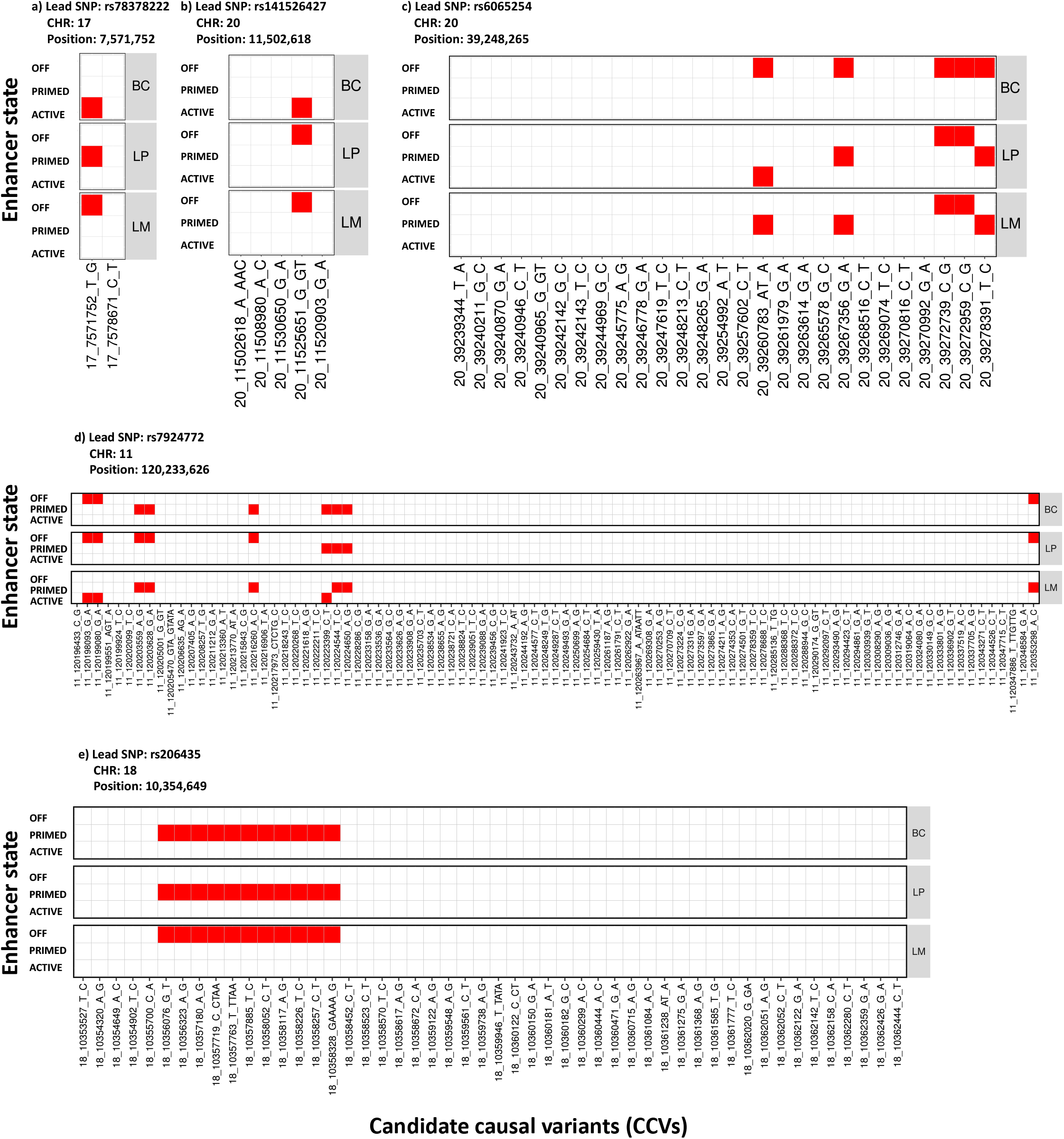
Heatmap of candidate causal variants (CCVs) overlapping results with enhancer states in primary breast subpopulations for five SNPs (**a)** rs78378222 **b)** rs141526427 **c**) rs6065254 **d**) rs7924772 **e**) rs206435 with associations in opposite direction across subtypes. Three different breast subpopulations were considered: basal cells (BC), luminal progenitor (LP) and luminal cells mature (LM). Based on a combination of H3K4me1 and H3K27ac histone modification ChiP-seq signals, putative enhancers in BC, LP, and LM were characterized as “OFF”, “PRIMED” and “ACTIVE” (**Online Methods**). The CCVs overlapping with enhancers were colored as red, otherwise were white.

**Figure 5.**
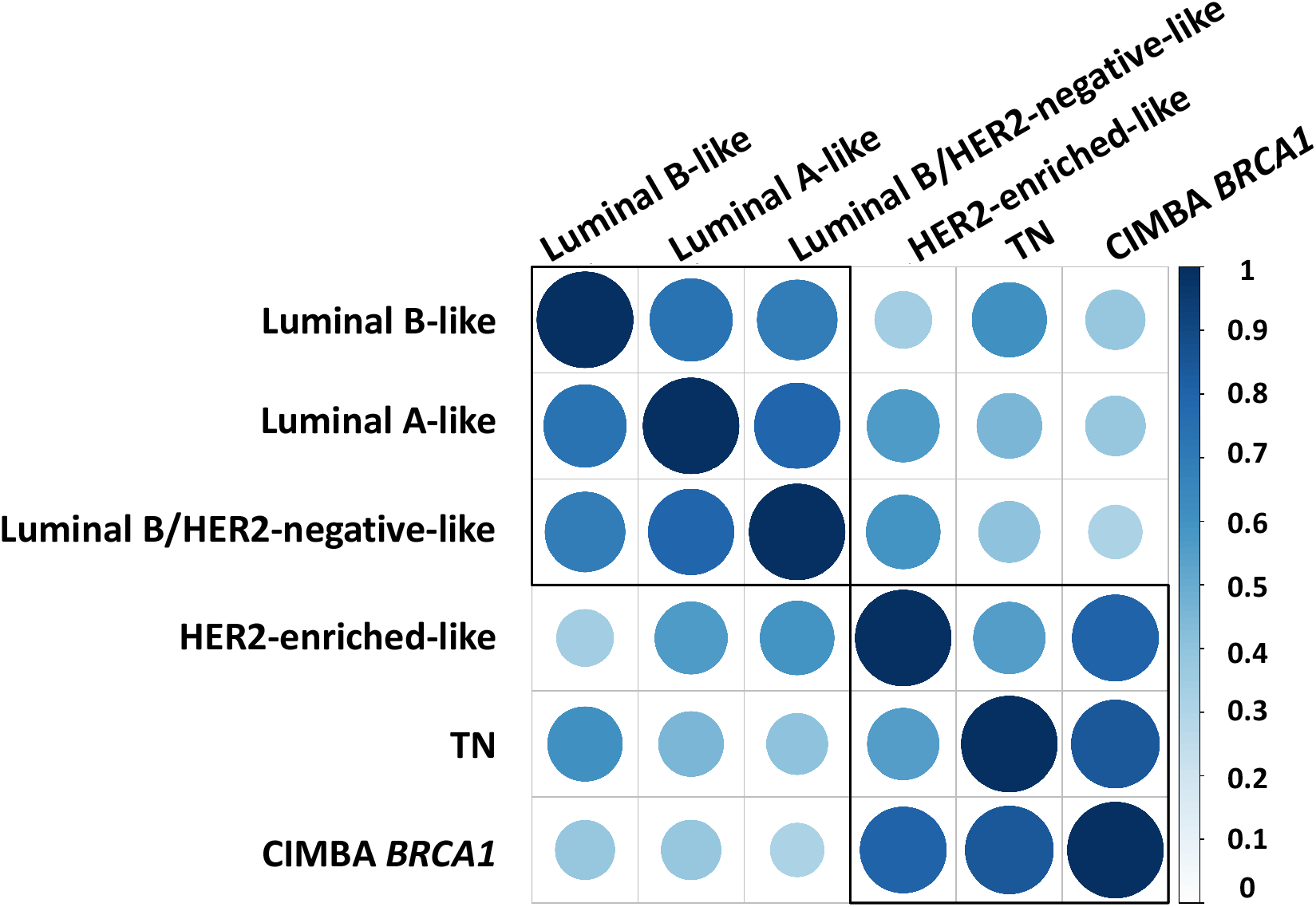
Genetic correlation between the five intrinsic-like breast cancer subtypes and *BRCA1* mutation carriers estimated through LD score regression. See **Supplementary Table 14** for further details. Both the color and size of the circles reflect the strength of the genetic correlations.

We used INQUIST to intersect each of the CCVs with functional annotation data from public databases to identify potential target genes^1^ (**Online Methods**, **Supplementary Table 14**). We predicted 179 unique target genes for 26 of the 32 independent signals. Twenty-three target genes in 14 regions were predicted with high confidence (designated “Level 1”), of which 22 target genes in 13 regions were predicted to be distally regulated. These targets include four genes predicted as INQUISIT targets in previous studies^12, 13^ *POLR3C*, *RNF115*, *SOX4* and *TBX3*, a known somatic breast cancer driver gene^14^, and genes implicated by transcriptome-wide association studies (*LINC00886*^15^ and *YBEY*^16^).

We used LD-regression to investigate the genetic architecture of molecular subtypes by evaluating the genetic correlations^17, 18^ between subtypes and to compare enrichment of genomic features^19^ between luminal A-like and TN subtypes (**Online Methods**). All intrinsic-like subtypes were moderately-to highly-correlated, with luminal B-like and HER2-enriched-like (r=0.35, standard error (SE) =0.10) having the lowest genetic correlation. The correlations between luminal A-like and TN was r=0.46 (SE=0.05), and between TN and *BRCA 1* carriers was r=0.84 (SE=0.15) (Figure 4; **Supplementary Table 15**). To compare genomic enrichment, we first evaluated 53 annotations and found TN tumors were most enriched for “super-enhancers, extend500bp” (3.04-fold, *P*= 3.3×10^-6^), and “digital genomic footprint, extend500bp” (from DNase hypersensitive sites) (2.2-fold, *P*=4.0×10^-4^) (**Supplementary Table 16, Supplementary Figure 10**). However, none of the 53 annotations significantly differed between luminal A-like and TN tumors. We also investigated cell-specific enrichment of four histone markers - H3K4me1, H3K3me3, H3K9ac and H3K27ac (**Online Methods**) - and found enrichment in both luminal-A and TN subtypes for gastrointestinal cell types and suppression of central nervous system cell types (**Supplementary Figure 11**).

The proportion of genome-wide chip heritability explained by the 32 SNPs identified in these analyses, plus 178 previously identified susceptibility SNPs^1, 2, 20^, was 54.2% for luminal A-like disease, 37.6% for TN disease and 26.9% for *BRCA1* carriers (Table1). Moreover, these 210 susceptibility SNPs explained approximately 18.3% of the two-fold familial relative risk for invasive breast cancer, while all reliably imputable SNPs on the OncoArray could explained 37.1% (**Online Methods**). Collectively, these analyses demonstrate the benefit of combining standard GWAS methods with methods accounting for underlying tumor heterogeneity. Moreover, these methods and results may help clarify mechanisms predisposing to specific molecular subtypes, and provide precise risk estimates for molecular subtypes to inform the development of subtype-specific polygenic risk scores^21^.

**Table 1.** Genetic variance of invasive breast cancer explained by identified susceptibility SNPs and all reliably genome-wide imputable variants^1^

| Phenotype | Genetic variance for 210 identified susceptibility SNPs<sup>2</sup> | Genetic variance for 32 newly identified SNPs<sup>2</sup> | Genetic variance for all GWAS variants<sup>3</sup> | Proportion of genetic variance explained by identified susceptibility loci<sup>4</sup> |
| --- | --- | --- | --- | --- |
| **Invasive breast cancer<sup>5</sup>** | 0.253 | 0.016 | 0.515 | 45.51% |
| **Luminal A-like** | 0.336 | 0.022 | 0.620 | 54.22% |
| **Luminal B/HER2-negative-like** | 0.233 | 0.018 | 0.597 | 38.95% |
| **Luminal B-like** | 0.270 | 0.020 | 0.740 | 36.46% |
| **HER2-enriched-like** | 0.200 | 0.011 | 0.689 | 29.05% |
| **Triple negative** | 0.185 | 0.025 | 0.492 | 37.63% |
| **CIMBA *BRCA1* carriers** | 0.083 | 0.016 | 0.309 | 26.86% |

<sup>1</sup> Genetic variance corresponds to heritability on the frailty-scale, which assumes the polygenetic log-additive model as the underlying model.
<sup>2</sup> Susceptibility SNPs included 178 SNPs identified or replicated in Nature 551, 92-94 (2017) and Nat Genet 49, 1767-1778 (2017) and 32 newly identified SNPs in this paper.
<sup>3</sup> Genetic variance of all reliably genome-wide imputable variants was estimated through LD-score regression described in Nat Genet 47, 291-5 (2015). and Nat Genet 47, 1236-41 (2015). Under the frailty-scale, the genetic variance for all GWAS variants is characterized by population variance of the underlying true polygenic risk score as $\sigma_{GWAS}^2 = Var(\sum_{m=1}^M \beta_m G_m)$ , where $G_m$ is the standardized genotype for the $m$ th SNP, $\beta_m$ is the true log odds ratio for the $m$ th SNP and $M$ are the total number of causal SNPs among the GWAS variants. (**Online Methods**).
<sup>4</sup> Proportion of genetic variance explained by 210 identified GWAS significant SNPs over the genetic variance explained by all GWAS variants.
<sup>5</sup> Invasive breast cancer summary level statistics were generated from 106,278 invasive cases and 91,477 controls, which were the same samples used in subtypes analyses (**Supplementary Table 2**).

## Online Methods

### Study populations

The overall breast cancer analyses included women of European ancestry from 82 BCAC studies from over 20 countries, with genotyping data derived from two Illumina genome-wide custom arrays, the iCOGS and OncoArray (**Supplementary Table 1**). Most of the studies were case-control studies in the general population, or hospital setting, or nested within population-based cohorts, but a subset of studies oversampled cases with a family history of the disease. We included controls and cases of invasive breast cancer, carcinoma *in-situ*, and cases of unknown invasiveness. Information on clinicopathologic characteristics were collected by the individual studies and combined in a central database after quality control checks. We used BCAC database version ‘freeze’ 10 for these analyses. Among a subset of participants (n=16,766) that were genotyped on both the iCOGS and OncoArray arrays, we kept only the OncoArray data. One study (LMBC) contributing to the iCOGS dataset was excluded due to inflation of the test statistics that was not corrected by adjustment for the first ten PCs. We also excluded OncoArray data from Norway (the Norwegian Breast Cancer Study) because there were no controls available from Norway with OncoArray data. All participating studies were approved by their appropriate ethics or institutional review board and all participants provided informed consent. The total sample size for this analysis, including iCOGS, OncoArray and other GWAS data, comprised 133,384 cases and 113,789 controls.

In the GWAS analyses accounting for underlying heterogeneity according to ER, PR, HER2 and grade, we included genotyping data from 81 BCAC studies. These analyses were restricted to controls and cases of invasive breast cancer; we excluded cases of carcinoma *in-situ* and cases with missing information on invasiveness, as these cases would potentially bias the implicit “imputation” of tumor marker in the underlying EM algorithm (**Supplemental Table 2)**. We also excluded all studies from a specific country if there were no controls for that country, or if the tumor marker data were missing on two or more of the tumor marker subtypes (see footnote of **Supplemental Table 2** for further explanation of excluded studies). We did not include the summary results from the 14,910 cases and 17,588 controls from the 11 other GWAS in subtype analyses because these studies did not provide data on tumor characteristics. We also excluded invasive cases (n=293) and controls (n=4,285) with missing data on age at diagnosis or age at enrollment, information required by the EM algorithm to impute missing tumor characteristics. In total, the final sample for the two-stage polytomous logistic regression comprised 106,278 invasive cases and 91,477 controls.

Participants included from CIMBA were women of European ancestry, aged 18 years or older with a pathogenic *BRCA1* variant. Most participants were sampled through cancer genetics clinics. In some instances, multiple members of the same family were enrolled. OncoArray genotype data was available from 58 studies from 24 countries. Following quality control and removal of participants that overlapped with the BCAC OncoArray study, data were available on 15,566 *BRCA1* mutation carriers, of whom 7,784 were affected with breast cancer (**Supplementary Table 3**). We also obtained iCOGS genotype data on 3,342 *BRCA1* mutation carriers (1,630 with breast cancer) from 54 studies through CIMBA. All *BRCA1* mutation carriers provided written informed consent and participated under ethically approved protocols.

### Genotyping, quality control, and imputation

Details on genotype calling, quality control and imputation for the OncoArray, iCOGS, and GWAS are described elsewhere^1, 2, 5, 6^. Genotyped or imputed SNPs marking each of the loci were determined using the iCOGS and the OncoArray genotyping arrays and imputation to the 1000 Genomes Project (Phase 3) reference panel. We included SNPs, from each component GWAS with an imputation quality score of >0.3. We restricted analysis to SNPs with a minor allele frequency >0.005 in the overall breast cancer analysis and >0.01 in the subtype analysis.

### Known breast cancer susceptibility variants

Prior studies identified susceptibility SNPs from genome-wide analyses at a significance level *P*< 5.0×10^−8^ for all breast cancer types, ER-negative or ER-positive breast cancer, in *BRCA1* or *BRCA2* mutation carriers, or in meta-analyses of these^1, 2^. We defined known breast cancer susceptibility variants as those variants that were identified or replicated in prior BCAC analyses^1, 2^. We also excluded from consideration variants within 500kb of a previously published locus, since these regions have been subject to separate conditional analyses^13^.

### Standard analysis of BCAC data

Logistic regression analyses were conducted separately for the iCOGS and OncoArray datasets, adjusting for country and the array-specific first 10 PCs for ancestry informative SNPs. The methods for estimating PCs have been described elsewhere^1, 2^. For the remaining GWAS, adjustment for inflation was done by adjusting for up to three PCs and using genomic control adjustment, as previously described^1^. We evaluated the associations between approximately 10.8 million SNPs with imputation quality scores (r^2^) ≥0.3 and MAF >0.005. We excluded SNPs located within ±500 KB of, or in LD (r^2^≥0.1) with known susceptibility SNPs^20^. The association effect size estimates from these, and the previously derived estimates from the 11 other GWAS, were then combined using a fixed effects meta-analysis. Since individual level genotyping data were not available for some previous GWAS, we conservatively approximated the potential overlap between the GWAS and iCOGS and OncoArray datasets, based on the populations contributing to each GWAS (iCOGS/GWAS: 626 controls and 923 cases; OncoArray/GWAS: 20 controls and 990 cases). We then used these adjusted data to estimate the correlation in the effect size estimates, and incorporated these into the meta-analysis using the method of Lin and Sullivan^22^.

### Subtypes analysis of BCAC data

We described the two-stage polytomous logistic regression in more detail elsewhere^4, 23^ (**Supplementary Note**). In brief, this method allows for efficient testing of a SNP-disease association in the presence of tumor subtype heterogeneity defined by multiple tumor characteristics, while accounting for multiple testing and missing data on tumor characteristics. In the first stage, the model uses a polytomous logistic regression to model case-control ORs between the SNPs and all possible subtypes that could be of interest, defined by the combination of the tumor markers. For example, in a model fit to evaluate heterogeneity according to ER, PR and HER2 positive/negative status, and grade of differentiation (low, intermediate and high grade), the first stage incorporates case-control ORs for 24 subtypes defined by the cross-classification of these factors. The second stage restructures the first-stage subtype-specific case-control ORs parameters into second-stage parameters through a decomposition procedure resulting in a second-stage baseline parameter that represents a case-control OR of a baseline cancer subtype, and case-case ORs parameters for each individual tumor characteristic. The second-stage case-case parameters can be used to perform heterogeneity tests with respect to each specific tumor marker while adjusting for the other tumor markers in the model. The two-stage model efficiently handles missing data by implementing an Expectation-Maximization algorithm^4, 7^ that essentially performs iterative “imputation” of the missing tumor characteristics conditional on available tumor characteristics and baseline covariates based on an underlying two-stage polytomous model.

To identify novel susceptibility loci, we used both a fixed-effect two-stage polytomous model and a mixed-effect two-stage polytomous model. The score-test we developed based on the mixed-effect model allows coefficients associated with individual tumor characteristics to enter as either fixed- or random-effect terms. Our previous analyses have shown that incorporation of random effect terms can improve power of the score-test by essentially reducing the effective degrees-of-freedom associated with fixed effects related to exploratory markers (*i.e.,* markers for which there is little prior evidence to suggest that they are a source of heterogeneity)^24^. On the other hand, incorporation of fixed-effect terms can preserve distinct associations of known important tumor characteristics, such as ER. In the mixed-effect two-stage polytomous model, we therefore kept ER as a fixed effect, but modeled PR, HER2 and grade as random effects. We evaluated SNPs with MAF >0.01 (∼10.0 million) and r^2^≥0.3, and excluded SNPs within ±500 kb of, or in LD (r^2^≥0.1) with known susceptibility SNPs, including those identified in the standard analysis for overall breast cancer. We reported SNPs that passed the p-value threshold of P < 5.0×10^-8^ in either the fixed- or mixed-effects models.

We assessed the influence of country on signals identified by the two-stage models by performing a ‘leave one out’ sensitivity analyses in which we reevaluated novel signals after excluding data from each individual country. Data from the OncoArray and iCOGS arrays were analyzed separately and then meta-analyzed using fixed-effects meta-analysis.

### Statistical analysis of CIMBA data

We tested for associations between SNPs and breast cancer risk for *BRCA1* mutation carriers using a score test statistic based on the retrospective likelihood of observing the SNP genotypes conditional on breast cancer phenotypes (breast cancer status and censoring time)^25^. Analyses were performed separately for iCOGS and OncoArray data. To allow for non-independence among related individuals, a kinship-adjusted test was used that accounted for familial correlations^26^. We stratified analyses by country of residence and, for countries where the strata were sufficiently large (United States and Canada), by Ashkenazi Jewish ancestry. The results from the iCOGS and OncoArray data were then pooled using fixed-effects meta-analysis.

### Meta-analysis of BCAC and CIMBA

We performed a fixed-effects meta-analysis of the results from BCAC TN cases and CIMBA *BRCA1* mutation carriers, using an inverse-variance fixed-effects approach implemented in METAL^27^. The estimates of association used were the logarithm of the per-allele hazard ratio estimate for association with breast cancer risk for *BRCA1* mutation carriers from CIMBA and the logarithm of the per-allele odds ratio estimate for association with risk of TN breast cancer based on BCAC data.

### Conditional analyses

We performed two sets of conditional analyses. First, we investigated for evidence of multiple independent signals in identified loci by performing forward selection logistic regression, in which we adjusted the lead SNP and analyzed association for all remaining SNPs within ±500 kb of the lead SNPs, irrespective of LD. Second, we confirmed the independence of 20 SNPs that were located within ±2 MB of a known susceptibility region by conditioning the identified signals on the nearby known signal. Since these 20 SNPs are already genome-wide significant in the original GWAS scan and the conditional analyses restricted to local regions, we therefore used a significance threshold of *P*<1×10^-6^ to control for type-one error^28^.

### Heterogeneity analysis of new association signals

We evaluated all novel signals for evidence of heterogeneity using the two-stage polytomous model. We first performed a global test for heterogeneity under the mixed-effect model test to identify SNPs showing evidence of heterogeneity with respect to any of the underlying tumor markers, ER, PR, HER2 and/or grade. We accounted for multiple testing of the global heterogeneity test using a FDR <0.05 under the Benjamini-Hochberg procedure^29^. Among the SNPs with observed heterogeneity, we then further used a fixed-effect two-stage model to evaluate influence of specific tumor characteristic(s) driving observed heterogeneity, adjusted for the other markers in the model. We also fit a separate fixed-effect two-stage models to estimate case-control ORs and 95% confidence intervals (CI) for five surrogate intrinsic-like subtypes defined by combinations of ER, PR, HER2 and grade^30^: (1) luminal A-like (ER+ and/or PR+, HER2-, grade 1 & 2); (2) luminal B/HER2-negative-like (ER+ and/or PR+, HER2-, grade 3); (3) luminal B-like (ER+ and/or PR+, HER2+); (4) HER2-enriched-like (ER- and PR-, HER2+), and (5) TN (ER-, PR-, HER2-).

### Candidate causal variants

We defined credible sets of candidate causal variants (CCVs) as variants located within ±500kb of the lead SNPs in each novel region and with *P* values within 100-fold of magnitude of the lead SNPs. This is approximately equivalent to selecting variants whose posterior probability of causality is within two orders of magnitude of the most significant SNP^31, 32^. This approach was applied for detecting a set of potentially causal variants for all 32 identified SNPs. For the novel SNPs located within ±2Mb of the known signals, we used the conditional *P* values to adjust for the known signals’ associations.

### eQTL Analysis

Data from breast cancer tumors and adjacent normal breast tissue were accessed from The Cancer Genome Atlas (TCGA)^33^. Germline SNP genotypes (Affymetrix 6.0 arrays) were processed and imputed to the 1000 Genomes reference panel (October 2014) and European ancestry ascertained as previously described^1^. Tumor tissue copy number was estimated from the Affymetrix 6.0 and called using the GISTIC2 algorithm^34^. Complete genotype, RNA-seq and copy number data were available for 679 genetically European patients (78 with adjacent normal tissue). Further, RNA-seq for normal breast tissue and imputed germline genotype data were available from 80 females from the GTEx Consortium^35^. Genes with a median expression level of 0 RPKM across samples were removed, and RPKM values of each gene were log2 transformed. Expression values of samples were quantile normalized. Genetic variants were evaluated for association with the expression of genes located within ±2Mb of the lead variant at each risk region using linear regression models, adjusting for ESR1 expression. Tumor tissue was also adjusted for copy number variation, as previously described^36^. eQTL analyses were performed using the MatrixEQTL program in R^37^.

### INQUISIT target gene analysis

#### Logic underlying INQUISIT predictions

Details of the INQUISIT pipeline have been previously described^1^. Briefly, genes were evaluated as potential targets of candidate causal variants through effects on: (1) distal gene regulation, (2) proximal regulation, or (3) a gene’s coding sequence. We intersected CCV positions with multiple sources of genomic information, chromatin interaction analysis by paired-end tag sequencing (ChIA-PET)^38^ in MCF7 cells, and genome-wide chromosome conformation capture (Hi-C) in HMECs^39^. We used breast cell line computational enhancer–promoter correlations (PreSTIGE^40^, IM-PET^41^, FANTOM5^42^) breast cell super-enhancer^43^, breast tissue-specific expression variants (eQTL) from multiple independent studies (TCGA (normal breast and breast tumor) and GTEx breast, **See eQTL Methods**), transcription factor and histone modification chromatin immunoprecipitation followed by sequencing (ChIP-seq) from the ENCODE and Roadmap Epigenomics Projects together with the genomic features found to be significantly enriched for all known breast cancer CCVs^13^, gene expression RNA-seq from several breast cancer lines and normal samples (ENCODE) and topologically associated domain (TAD) boundaries from T47D cells (ENCODE^44^). To assess the impact of intragenic variants, we evaluated their potential to alter primary protein coding sequence and splicing using Ensembl Variant Effect Predictor^45^ using MaxEntScan and dbscSNV modules for splicing alterations based on “ada” and “rf” scores. Nonsense and missense changes were assessed with the REVEL ensemble algorithm, with CCVs displaying REVEL scores > 0.5 deemed deleterious.

#### Scoring hierarchy

Each target gene prediction category (distal, promoter or coding) was scored according to different criteria. Genes predicted to be distally-regulated targets of CCVs were awarded two points based on physical links (for example ChIA-PET), and one point for computational prediction methods, or eQTL associations. All CCVs were considered as potentially involved in distal regulation and all CCVs (including coding SNPs) were scored in this category. Intersection of a putative distal enhancer with genomic features found to be significantly enriched^20^ were further upweighted with an additional point. In the case of multiple, independent interactions, an additional point was awarded. CCVs in gene proximal regulatory regions were intersected with histone ChIP-Seq peaks characteristic of promoters and assigned to the overlapping transcription start sites (defined as −1.0 kb - +0.1 kb). Further points were awarded to such genes if there was evidence for an eQTL association, while a lack of expression resulted in down-weighting as potential targets. Potential coding changes including missense, nonsense and predicted splicing alterations resulted in addition of one point to the encoded gene for each type of change, while lack of expression reduced the score. We added an additional point for predicted target genes that were also breast cancer drivers (278 genes^1, 20^). For each category, scores potentially ranged from 0-8 (distal); 0-4 (promoter) or 0-3 (coding). We converted these scores into ‘confidence levels’: Level 1 (highest confidence) when distal score >4, promoter score ≥3 or coding score >1; Level 2 when distal score ≤4 and ≥1, promoter score=1 or=2, coding score=1; and Level 3 when distal score <1 and >0, promoter score <1 and >0, and coding <1 and >0. For genes with multiple scores (for example, predicted as targets from multiple independent risk signals or predicted to be impacted in several categories), we recorded the highest score.

### Enhancer states analysis in breast sub-populations

We obtained enhancer maps for three enriched primary breast sub-populations (basal, luminal progenitor, and mature luminal) from Pellacani et al.^11^. Enhancer annotations were defined as ACTIVE, PRIMED, or OFF based on a combination of H3K27ac and H3K4me1 histone modification ChIP-seq signals using FPKM thresholds as previously described^11^. Briefly, genomic regions containing high H3K4me1 signal observed in any cell type were used to define the superset of breast regulatory elements. Sub-population cell type-specific H3K27ac signal (which is characteristic of active elements) within these elements was used as a measure of overall regulatory activity, where “ACTIVE” sites were characterized by H3K4me1-high, H3K27ac-high; “PRIMED” by H3K4me1-high, H3K27ac-low; and “OFF” by H3K4me1-low, H3K27ac-low. This enabled annotation of each enhancer element as either “OFF”, “PRIMED” or “ACTIVE” in all cell types. We then defined enhancers which exhibit differing states between at least one cell type as “ANYSWITCH” enhancers.

### Genetic correlation analyses

We used LD score regression^17–19^ to estimate the genetic correlation between five intrinsic-like breast cancer subtypes. The analysis used the summary statistics based on the meta-analysis of the OncoArray, and iCOGS, and CIMBA meta-analysis. The genetic correlation^17^ analysis was restricted to the ∼1 million SNPs included in HapMap 3 with MAF > 1% and imputation quality score R2>0.3 in the OncoArray data. Since two-stage polytomous models integrated an imputation algorithm for missing tumor characteristic data, we modified the LD score regression to generate the effective sample size for each SNP (**Supplementary Note**).

### Global genomic enrichment analyses

We performed stratified LD score regression analyses^17–19^ as previously described^1^ for two major intrinsic-like subtypes, luminal A-like and TN, using the summary statistics from the meta-analyses of OncoArray, iCOGs, and CIMBA. The analysis included all SNPs in the 1000 Genome Project Phase 1v3 release with MAF>1% and imputation quality score R2>0.3 in the OncoArray data. We restricted analysis to all SNPs present on the HapMap version 3 dataset. We first fit a model that included 24 non-cell-type-specific, publicly available annotations as well as 24 additional annotations that included a 500-bp window around each of the 24 main annotations. We also included 100-bp windows around ChIP–seq peaks and one annotation containing all SNPs, leading to a total of 53 overlapping annotations. In addition to the baseline model using 24 main annotations, we also performed cell-type-specific analyses using annotations of the four histone marks (H3K4me1, H3K4me3, H3K9ac and H3K27ac). Each cell-type-specific annotation corresponds to a histone mark in a single cell type (for example, H3K27ac in adipose nuclei tissues)^19^. There was a total of 220 such annotations. We further subdivided these 220 cell-type-specific annotations into 10 categories by aggregating the cell-type-specific annotations within each group (for example, SNPs related with any of the four histone modifications in any hematopoietic and immune cells were considered as one category). To estimate the enrichment of each marker, we ran 220 LD score regressions after adding each different histone mark to the baseline model. We used a Wald test to evaluate the differences in the functional enrichment between the luminal A-like and TN subtypes, using the regression coefficients and standard error based on the models above. After Bonferroni correction none of the differences were significant. Notably, the Wald test assumes that the enrichment estimates of luminal A-like and TN subtypes were independent, but this assumption was violated by the sharing of controls between the subtypes. Under this scenario, our Wald test statistics were less conservative than had we adjusted for the correlation between estimates. However, given the lack of significant differences observed between luminal A-like and TN subtypes we had no concern about a type one error.

### Genetic variance explained by identified susceptibility SNPs and all genome-wide imputable variants

Genetic variance corresponds to heritability on the frailty-scale, which assumes a polygenetic log-additive model as the underlying model. Under the log-additive model, the frailty-scale heritability explained by the identified variants can be estimated by:

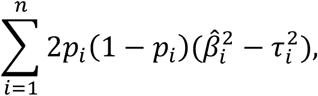

where *n* is the total number of identified SNPs, *p_i_* is the MAF for *i*th variant, 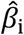 is the log odds ratio estimate for the *i*th variant, and *τ_i_* is the standard error of 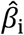. To obtain the frailty scale heritability for invasive breast cancer explained by all of the GWAS variants, we used LD score regression to estimate heritability 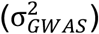 using the full set of summary statistics from either standard logistic regression for overall invasive breast cancer, the two-stage polytomous regression for the intrinsic-like subtypes, or the CIMBA *BRCA1* analysis for *BRCA1* carriers. 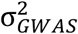 is characterized by population variance of the underlying true polygenetic risk scores as 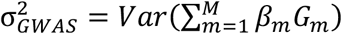, where *G_m_* is the standardized genotype for the *m*th SNP, *β_m_* is the true log odds ratio for the *m*th SNP and *M* are the total number of causal SNPs among the GWAS variants. Thus, the proportion of heritability explained by identified variants relative to all imputable SNPs is:

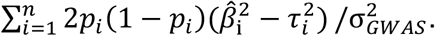

To estimate the proportion of the familial risk of invasive breast cancer that is explained by susceptibility variants, we defined the familial relative risk, λ, as the familial relative risk assuming a polygenic log-additive model that explains all the familial aggregation of the disease^46^. Under the frailty scale, we define the broad sense heritability^47^ as σ^2^. The relationship between λ and σ^2^ was shown to be σ^2^ = 2 ∗ log (L) ^46^. We assumed λ = 2 as the overall familial relative risk of invasive breast cancer, thus σ^2^ = 2log (2) and the proportion of the familial relative risk explained by identified susceptibility SNPs is 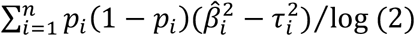, and the proportion of the familial relative risk explained by GWAS variants is 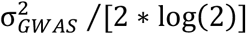. Analyses of heritability and the proportion of explained familial risk were restricted to 106,278 invasive cases and 91,477 controls (**Supplementary Table 2**).

## BCAC Funding and Acknowledgments

### Funding

BCAC is funded by Cancer Research UK [C1287/A16563, C1287/A10118], the European Union’s Horizon 2020 Research and Innovation Programme (grant numbers 634935 and 633784 for BRIDGES and B-CAST respectively), and by the European Communitýs Seventh Framework Programme under grant agreement number 223175 (grant number HEALTH-F2-2009-223175) (COGS). The EU Horizon 2020 Research and Innovation Programme funding source had no role in study design, data collection, data analysis, data interpretation or writing of the report.

Genotyping of the OncoArray was funded by the NIH Grant U19 CA148065, and Cancer UK Grant C1287/A16563 and the PERSPECTIVE project supported by the Government of Canada through Genome Canada and the Canadian Institutes of Health Research (grant GPH-129344) and, the Ministère de l’Économie, Science et Innovation du Québec through Genome Québec and the PSRSIIRI-701 grant, and the Quebec Breast Cancer Foundation. Funding for the iCOGS infrastructure came from: the European Community’s Seventh Framework Programme under grant agreement n° 223175 (HEALTH-F2-2009-223175) (COGS), Cancer Research UK (C1287/A10118, C1287/A10710, C12292/A11174, C1281/A12014, C5047/A8384, C5047/A15007, C5047/A10692, C8197/A16565), the National Institutes of Health (CA128978) and Post-Cancer GWAS initiative (1U19 CA148537, 1U19 CA148065 and 1U19 CA148112 - the GAME-ON initiative), the Department of Defence (W81XWH-10-1-0341), the Canadian Institutes of Health Research (CIHR) for the CIHR Team in Familial Risks of Breast Cancer, and Komen Foundation for the Cure, the Breast Cancer Research Foundation, and the Ovarian Cancer Research Fund. The DRIVE Consortium was funded by U19 CA148065.

The Australian Breast Cancer Family Study (ABCFS) was supported by grant UM1 CA164920 from the National Cancer Institute (USA). The content of this manuscript does not necessarily reflect the views or policies of the National Cancer Institute or any of the collaborating centers in the Breast Cancer Family Registry (BCFR), nor does mention of trade names, commercial products, or organizations imply endorsement by the USA Government or the BCFR. The ABCFS was also supported by the National Health and Medical Research Council of Australia, the New South Wales Cancer Council, the Victorian Health Promotion Foundation (Australia) and the Victorian Breast Cancer Research Consortium. J.L.H. is a National Health and Medical Research Council (NHMRC) Senior Principal Research Fellow. M.C.S. is a NHMRC Senior Research Fellow. The ABCS study was supported by the Dutch Cancer Society [grants NKI 2007-3839; 2009 4363]. The Australian Breast Cancer Tissue Bank (ABCTB) was supported by the National Health and Medical Research Council of Australia, The Cancer Institute NSW and the National Breast Cancer Foundation. The AHS study is supported by the intramural research program of the National Institutes of Health, the National Cancer Institute (grant number Z01-CP010119), and the National Institute of Environmental Health Sciences (grant number Z01-ES049030). The work of the BBCC was partly funded by ELAN-Fond of the University Hospital of Erlangen. The BBCS is funded by Cancer Research UK and Breast Cancer Now and acknowledges NHS funding to the NIHR Biomedical Research Centre, and the National Cancer Research Network (NCRN). The BCEES was funded by the National Health and Medical Research Council, Australia and the Cancer Council Western Australia and acknowledges funding from the National Breast Cancer Foundation (JS). For the BCFR-NY, BCFR-PA, BCFR-UT this work was supported by grant UM1 CA164920 from the National Cancer Institute. The content of this manuscript does not necessarily reflect the views or policies of the National Cancer Institute or any of the collaborating centers in the Breast Cancer Family Registry (BCFR), nor does mention of trade names, commercial products, or organizations imply endorsement by the US Government or the BCFR. For BIGGS, ES is supported by NIHR Comprehensive Biomedical Research Centre, Guy’s & St. Thomas’ NHS Foundation Trust in partnership with King’s College London, United Kingdom. IT is supported by the Oxford Biomedical Research Centre. The BREast Oncology GAlician Network (BREOGAN) is funded by Acción Estratégica de Salud del Instituto de Salud Carlos III FIS PI12/02125/Cofinanciado FEDER; Acción Estratégica de Salud del Instituto de Salud Carlos III FIS Intrasalud (PI13/01136); Programa Grupos Emergentes, Cancer Genetics Unit, Instituto de Investigacion Biomedica Galicia Sur. Xerencia de Xestion Integrada de Vigo-SERGAS, Instituto de Salud Carlos III, Spain; Grant 10CSA012E, Consellería de Industria Programa Sectorial de Investigación Aplicada, PEME I + D e I + D Suma del Plan Gallego de Investigación, Desarrollo e Innovación Tecnológica de la Consellería de Industria de la Xunta de Galicia, Spain; Grant EC11-192. Fomento de la Investigación Clínica Independiente, Ministerio de Sanidad, Servicios Sociales e Igualdad, Spain; and Grant FEDER-Innterconecta. Ministerio de Economia y Competitividad, Xunta de Galicia, Spain. The BSUCH study was supported by the Dietmar-Hopp Foundation, the Helmholtz Society and the German Cancer Research Center (DKFZ). CBCS is funded by the Canadian Cancer Society (grant # 313404) and the Canadian Institutes of Health Research. CCGP is supported by funding from the University of Crete. The CECILE study was supported by Fondation de France, Institut National du Cancer (INCa), Ligue Nationale contre le Cancer, Agence Nationale de Sécurité Sanitaire, de l’Alimentation, de l’Environnement et du Travail (ANSES), Agence Nationale de la Recherche (ANR). The CGPS was supported by the Chief Physician Johan Boserup and Lise Boserup Fund, the Danish Medical Research Council, and Herlev and Gentofte Hospital. The CNIO-BCS was supported by the Instituto de Salud Carlos III, the Red Temática de Investigación Cooperativa en Cáncer and grants from the Asociación Española Contra el Cáncer and the Fondo de Investigación Sanitario (PI11/00923 and PI12/00070). COLBCCC is supported by the German Cancer Research Center (DKFZ), Heidelberg, Germany. Diana Torres was in part supported by a postdoctoral fellowship from the Alexander von Humboldt Foundation. The American Cancer Society funds the creation, maintenance, and updating of the CPS-II cohort. The CTS was initially supported by the California Breast Cancer Act of 1993 and the California Breast Cancer Research Fund (contract 97-10500) and is currently funded through the National Institutes of Health (R01 CA77398, UM1 CA164917, and U01 CA199277). Collection of cancer incidence data was supported by the California Department of Public Health as part of the statewide cancer reporting program mandated by California Health and Safety Code Section 103885. HAC receives support from the Lon V Smith Foundation (LVS39420). The University of Westminster curates the DietCompLyf database funded by Against Breast Cancer Registered Charity No. 1121258 and the NCRN. The coordination of EPIC is financially supported by the European Commission (DG-SANCO) and the International Agency for Research on Cancer. The national cohorts are supported by: Ligue Contre le Cancer, Institut Gustave Roussy, Mutuelle Générale de l’Education Nationale, Institut National de la Santé et de la Recherche Médicale (INSERM) (France); German Cancer Aid, German Cancer Research Center (DKFZ), Federal Ministry of Education and Research (BMBF) (Germany); the Hellenic Health Foundation, the Stavros Niarchos Foundation (Greece); Associazione Italiana per la Ricerca sul Cancro-AIRC-Italy and National Research Council (Italy); Dutch Ministry of Public Health, Welfare and Sports (VWS), Netherlands Cancer Registry (NKR), LK Research Funds, Dutch Prevention Funds, Dutch ZON (Zorg Onderzoek Nederland), World Cancer Research Fund (WCRF), Statistics Netherlands (The Netherlands); Health Research Fund (FIS), PI13/00061 to Granada, PI13/01162 to EPIC-Murcia, Regional Governments of Andalucía, Asturias, Basque Country, Murcia and Navarra, ISCIII RETIC (RD06/0020) (Spain); Cancer Research UK (14136 to EPIC-Norfolk; C570/A16491 and C8221/A19170 to EPIC-Oxford), Medical Research Council (1000143 to EPIC-Norfolk, MR/M012190/1 to EPIC-Oxford) (United Kingdom). The ESTHER study was supported by a grant from the Baden Württemberg Ministry of Science, Research and Arts. Additional cases were recruited in the context of the VERDI study, which was supported by a grant from the German Cancer Aid (Deutsche Krebshilfe). FHRISK is funded from NIHR grant PGfAR 0707-10031. The GC-HBOC (German Consortium of Hereditary Breast and Ovarian Cancer) is supported by the German Cancer Aid (grant no 110837 and 113049, coordinator: Rita K. Schmutzler, Cologne). This work was also funded by the European Regional Development Fund and Free State of Saxony, Germany (LIFE - Leipzig Research Centre for Civilization Diseases, project numbers 713-241202, 713-241202, 14505/2470, 14575/2470). The GENICA was funded by the Federal Ministry of Education and Research (BMBF) Germany grants 01KW9975/5, 01KW9976/8, 01KW9977/0 and 01KW0114, the Robert Bosch Foundation, Stuttgart, Deutsches Krebsforschungszentrum (DKFZ), Heidelberg, the Institute for Prevention and Occupational Medicine of the German Social Accident Insurance, Institute of the Ruhr University Bochum (IPA), Bochum, as well as the Department of Internal Medicine, Evangelische Kliniken Bonn gGmbH, Johanniter Krankenhaus, Bonn, Germany. Generation Scotland (GENSCOT) received core support from the Chief Scientist Office of the Scottish Government Health Directorates [CZD/16/6] and the Scottish Funding Council [HR03006]. Genotyping of the GS:SFHS samples was carried out by the Genetics Core Laboratory at the Edinburgh Clinical Research Facility, University of Edinburgh, Scotland and was funded by the Medical Research Council UK and the Wellcome Trust (Wellcome Trust Strategic Award “STratifying Resilience and Depression Longitudinally” (STRADL) Reference 104036/Z/14/Z). Funding for identification of cases and contribution to BCAC funded in part by the Wellcome Trust Seed Award “Temporal trends in incidence and mortality of molecular subtypes of breast cancer to inform public health, policy and prevention” Reference 207800/Z/17/Z. The GEPARSIXTO study was conducted by the German Breast Group GmbH. The GESBC was supported by the Deutsche Krebshilfe e. V. [70492] and the German Cancer Research Center (DKFZ). The HABCS study was supported by the Claudia von Schilling Foundation for Breast Cancer Research, by the Lower Saxonian Cancer Society, and by the Rudolf Bartling Foundation. The HEBCS was financially supported by the Helsinki University Hospital Research Fund, the Finnish Cancer Society, and the Sigrid Juselius Foundation..The HMBCS was supported by a grant from the Friends of Hannover Medical School and by the Rudolf Bartling Foundation. The HUBCS was supported by a grant from the German Federal Ministry of Research and Education (RUS08/017), B.M. was supported by grant 17-44-020498, 17-29-06014 of the Russian Foundation for Basic Research, D.P. was supported by grant 18-29-09129 of the Russian Foundation for Basic Research, E.K was supported by the program for support the bioresource collections №007-030164/2, and the study was performed as part of the assignment of the Ministry of Science and Higher Education of the Russian Federation (№АААА-А16-116020350032-1). Financial support for KARBAC was provided through the regional agreement on medical training and clinical research (ALF) between Stockholm County Council and Karolinska Institutet, the Swedish Cancer Society, The Gustav V Jubilee foundation and Bert von Kantzows foundation. The KARMA study was supported by Märit and Hans Rausings Initiative Against Breast Cancer. The KBCP was financially supported by the special Government Funding (EVO) of Kuopio University Hospital grants, Cancer Fund of North Savo, the Finnish Cancer Organizations, and by the strategic funding of the University of Eastern Finland. kConFab is supported by a grant from the National Breast Cancer Foundation, and previously by the National Health and Medical Research Council (NHMRC), the Queensland Cancer Fund, the Cancer Councils of New South Wales, Victoria, Tasmania and South Australia, and the Cancer Foundation of Western Australia. Financial support for the AOCS was provided by the United States Army Medical Research and Materiel Command [DAMD17-01-1-0729], Cancer Council Victoria, Queensland Cancer Fund, Cancer Council New South Wales, Cancer Council South Australia, The Cancer Foundation of Western Australia, Cancer Council Tasmania and the National Health and Medical Research Council of Australia (NHMRC; 400413, 400281, 199600). G.C.T. and P.W. are supported by the NHMRC. RB was a Cancer Institute NSW Clinical Research Fellow. LAABC is supported by grants (1RB-0287, 3PB-0102, 5PB-0018, 10PB-0098) from the California Breast Cancer Research Program. Incident breast cancer cases were collected by the USC Cancer Surveillance Program (CSP) which is supported under subcontract by the California Department of Health. The CSP is also part of the National Cancer Institute’s Division of Cancer Prevention and Control Surveillance, Epidemiology, and End Results Program, under contract number N01CN25403. LMBC is supported by the ‘Stichting tegen Kanker’. DL is supported by the FWO. The MABCS study is funded by the Research Centre for Genetic Engineering and Biotechnology “Georgi D. Efremov”, MASA. The MARIE study was supported by the Deutsche Krebshilfe e.V. [70-2892-BR I, 106332, 108253, 108419, 110826, 110828], the Hamburg Cancer Society, the German Cancer Research Center (DKFZ) and the Federal Ministry of Education and Research (BMBF) Germany [01KH0402]. MBCSG is supported by grants from the Italian Association for Cancer Research (AIRC) and by funds from the Italian citizens who allocated the 5/1000 share of their tax payment in support of the Fondazione IRCCS Istituto Nazionale Tumori, according to Italian laws (INT-Institutional strategic projects “5×1000”). The MCBCS was supported by the NIH grants CA192393, CA116167, CA176785 an NIH Specialized Program of Research Excellence (SPORE) in Breast Cancer [CA116201], and the Breast Cancer Research Foundation and a generous gift from the David F. and Margaret T. Grohne Family Foundation. The Melbourne Collaborative Cohort Study (MCCS) cohort recruitment was funded by VicHealth and Cancer Council Victoria. The MCCS was further augmented by Australian National Health and Medical Research Council grants 209057, 396414 and 1074383 and by infrastructure provided by Cancer Council Victoria. Cases and their vital status were ascertained through the Victorian Cancer Registry and the Australian Institute of Health and Welfare, including the National Death Index and the Australian Cancer Database. The MEC was supported by NIH grants CA63464, CA54281, CA098758, CA132839 and CA164973. The MISS study is supported by funding from ERC-2011-294576 Advanced grant, Swedish Cancer Society, Swedish Research Council, Local hospital funds, Berta Kamprad Foundation, Gunnar Nilsson. The MMHS study was supported by NIH grants CA97396, CA128931, CA116201, CA140286 and CA177150. MSKCC is supported by grants from the Breast Cancer Research Foundation and Robert and Kate Niehaus Clinical Cancer Genetics Initiative. The work of MTLGEBCS was supported by the Quebec Breast Cancer Foundation, the Canadian Institutes of Health Research for the “CIHR Team in Familial Risks of Breast Cancer” program – grant # CRN-87521 and the Ministry of Economic Development, Innovation and Export Trade – grant # PSR-SIIRI-701. The NBCS has received funding from the K.G. Jebsen Centre for Breast Cancer Research; the Research Council of Norway grant 193387/V50 (to A-L Børresen-Dale and V.N. Kristensen) and grant 193387/H10 (to A-L Børresen-Dale and V.N. Kristensen), South Eastern Norway Health Authority (grant 39346 to A-L Børresen-Dale) and the Norwegian Cancer Society (to A-L Børresen-Dale and V.N. Kristensen). The NBHS was supported by NIH grant R01CA100374. Biological sample preparation was conducted the Survey and Biospecimen Shared Resource, which is supported by P30 CA68485. The Northern California Breast Cancer Family Registry (NC-BCFR) and Ontario Familial Breast Cancer Registry (OFBCR) were supported by grant UM1 CA164920 from the National Cancer Institute (USA). The content of this manuscript does not necessarily reflect the views or policies of the National Cancer Institute or any of the collaborating centers in the Breast Cancer Family Registry (BCFR), nor does mention of trade names, commercial products, or organizations imply endorsement by the USA Government or the BCFR. The Carolina Breast Cancer Study was funded by Komen Foundation, the National Cancer Institute (P50 CA058223, U54 CA156733, U01 CA179715), and the North Carolina University Cancer Research Fund. The NHS was supported by NIH grants P01 CA87969, UM1 CA186107, and U19 CA148065. The NHS2 was supported by NIH grants UM1 CA176726 and U19 CA148065. The OBCS was supported by research grants from the Finnish Cancer Foundation, the Academy of Finland (grant number 250083, 122715 and Center of Excellence grant number 251314), the Finnish Cancer Foundation, the Sigrid Juselius Foundation, the University of Oulu, the University of Oulu Support Foundation and the special Governmental EVO funds for Oulu University Hospital-based research activities. The ORIGO study was supported by the Dutch Cancer Society (RUL 1997-1505) and the Biobanking and Biomolecular Resources Research Infrastructure (BBMRI-NL CP16). The PBCS was funded by Intramural Research Funds of the National Cancer Institute, Department of Health and Human Services, USA. Genotyping for PLCO was supported by the Intramural Research Program of the National Institutes of Health, NCI, Division of Cancer Epidemiology and Genetics. The PLCO is supported by the Intramural Research Program of the Division of Cancer Epidemiology and Genetics and supported by contracts from the Division of Cancer Prevention, National Cancer Institute, National Institutes of Health. The POSH study is funded by Cancer Research UK (grants C1275/A11699, C1275/C22524, C1275/A19187, C1275/A15956 and Breast Cancer Campaign 2010PR62, 2013PR044. PROCAS is funded from NIHR grant PGfAR 0707-10031. The RBCS was funded by the Dutch Cancer Society (DDHK 2004-3124, DDHK 2009-4318). The SASBAC study was supported by funding from the Agency for Science, Technology and Research of Singapore (A*STAR), the US National Institute of Health (NIH) and the Susan G. Komen Breast Cancer Foundation. The SBCS was supported by Sheffield Experimental Cancer Medicine Centre and Breast Cancer Now Tissue Bank. SEARCH is funded by Cancer Research UK [C490/A10124, C490/A16561] and supported by the UK National Institute for Health Research Biomedical Research Centre at the University of Cambridge. The University of Cambridge has received salary support for PDPP from the NHS in the East of England through the Clinical Academic Reserve. The Sister Study (SISTER) is supported by the Intramural Research Program of the NIH, National Institute of Environmental Health Sciences (Z01-ES044005 and Z01-ES049033). The Two Sister Study (2SISTER) was supported by the Intramural Research Program of the NIH, National Institute of Environmental Health Sciences (Z01-ES044005 and Z01-ES102245), and, also by a grant from Susan G. Komen for the Cure, grant FAS0703856. SKKDKFZS is supported by the DKFZ. The SMC is funded by the Swedish Cancer Foundation and the Swedish Research Council/Infrastructure gran. The SZBCS and IHCC were supported by Grant PBZ_KBN_122/P05/2004 and the program of the Minister of Science and Higher Education under the name “Regional Initiative of Excellence” in 2019-2022 project number 002/RID/2018/19 amount of financing 12 000 000 PLN. The UKBGS is funded by Breast Cancer Now and the Institute of Cancer Research (ICR), London. ICR acknowledges NHS funding to the NIHR Biomedical Research Centre. The UKOPS study was funded by The Eve Appeal (The Oak Foundation) and supported by the National Institute for Health Research University College London Hospitals Biomedical Research Centre. The USRT Study was funded by Intramural Research Funds of the National Cancer Institute, Department of Health and Human Services, USA. The WHI program is funded by the National Heart, Lung, and Blood Institute, the US National Institutes of Health and the US Department of Health and Human Services (HHSN268201100046C, HHSN268201100001C, HHSN268201100002C, HHSN268201100003C, HHSN268201100004C and HHSN271201100004C). This work was also funded by NCI U19 CA148065-01. The BCINIS was supported by the BCRF (breast cancer research foundation), NY. Nilanjan Chatterjee received supported from the National Human Genome Research Institute (1 R01 HG010480-01).

### Acknowledgements

We thank all the individuals who took part in these studies and all the researchers, clinicians, technicians and administrative staff who have enabled this work to be carried out. The COGS study would not have been possible without the contributions of the following: Rosalind A. Eeles, Ali Amin Al Olama, Zsofia Kote-Jarai, Lesley McGuffog, Andrew Lee, and Ed Dicks, Craig Luccarini and the staff of the Centre for Genetic Epidemiology Laboratory, Anna Gonzalez-Neira and the staff of the CNIO genotyping unit, Daniel C. Tessier, Francois Bacot, Daniel Vincent, Sylvie LaBoissière and Frederic Robidoux and the staff of the McGill University and Génome Québec Innovation Centre, Borge G. Nordestgaard, and the staff of the Copenhagen DNA laboratory, and Julie M. Cunningham, Sharon A. Windebank, Christopher A. Hilker, Jeffrey Meyer and the staff of Mayo Clinic Genotyping Core Facility. ABCFS thank Maggie Angelakos, Judi Maskiell, Gillian Dite. ABCS thanks the Blood bank Sanquin, The Netherlands. ABCTB Investigators: Christine Clarke, Rosemary Balleine, Robert Baxter, Stephen Braye, Jane Carpenter, Jane Dahlstrom, John Forbes, Soon Lee, Debbie Marsh, Adrienne Morey, Nirmala Pathmanathan, Rodney Scott, Allan Spigelman, Nicholas Wilcken, Desmond Yip. Samples are made available to researchers on a non-exclusive basis. BBCS thanks Eileen Williams, Elaine Ryder-Mills, Kara Sargus. BCEES thanks Allyson Thomson, Christobel Saunders, Terry Slevin, BreastScreen Western Australia, Elizabeth Wylie, Rachel Lloyd. The BCINIS study would not have been possible without the contributions of Dr. K. Landsman, Dr. N. Gronich, Dr. A. Flugelman, Dr. W. Saliba, Dr. E. Liani, Dr. I. Cohen, Dr. S. Kalet, Dr. V. Friedman, Dr. O. Barnet of the NICCC in Haifa, and all the contributing family medicine, surgery, pathology and oncology teams in all medical institutes in Northern Israel. BIGGS thanks Niall McInerney, Gabrielle Colleran, Andrew Rowan, Angela Jones. The BREOGAN study would not have been possible without the contributions of the following: Manuela Gago-Dominguez, Jose Esteban Castelao, Angel Carracedo, Victor Muñoz Garzón, Alejandro Novo Domínguez, Maria Elena Martinez, Sara Miranda Ponte, Carmen Redondo Marey, Maite Peña Fernández, Manuel Enguix Castelo, Maria Torres, Manuel Calaza (BREOGAN), José Antúnez, Máximo Fraga and the staff of the Department of Pathology and Biobank of the University Hospital Complex of Santiago-CHUS, Instituto de Investigación Sanitaria de Santiago, IDIS, Xerencia de Xestion Integrada de Santiago-SERGAS; Joaquín González-Carreró and the staff of the Department of Pathology and Biobank of University Hospital Complex of Vigo, Instituto de Investigacion Biomedica Galicia Sur, SERGAS, Vigo, Spain. BSUCH thanks Peter Bugert, Medical Faculty Mannheim. CBCS thanks study participants, co-investigators, collaborators and staff of the Canadian Breast Cancer Study, and project coordinators Agnes Lai and Celine Morissette. CCGP thanks Styliani Apostolaki, Anna Margiolaki, Georgios Nintos, Maria Perraki, Georgia Saloustrou, Georgia Sevastaki, Konstantinos Pompodakis. CGPS thanks staff and participants of the Copenhagen General Population Study. For the excellent technical assistance: Dorthe Uldall Andersen, Maria Birna Arnadottir, Anne Bank, Dorthe Kjeldgård Hansen. The Danish Cancer Biobank is acknowledged for providing infrastructure for the collection of blood samples for the cases. CNIO-BCS thanks Guillermo Pita, Charo Alonso, Nuria Álvarez, Pilar Zamora, Primitiva Menendez, the Human Genotyping-CEGEN Unit (CNIO). Investigators from the CPS-II cohort thank the participants and Study Management Group for their invaluable contributions to this research. They also acknowledge the contribution to this study from central cancer registries supported through the Centers for Disease Control and Prevention National Program of Cancer Registries, as well as cancer registries supported by the National Cancer Institute Surveillance Epidemiology and End Results program. The CTS Steering Committee includes Leslie Bernstein, Susan Neuhausen, James Lacey, Sophia Wang, Huiyan Ma, and Jessica Clague DeHart at the Beckman Research Institute of City of Hope, Dennis Deapen, Rich Pinder, and Eunjung Lee at the University of Southern California, Pam Horn-Ross, Peggy Reynolds, Christina Clarke Dur and David Nelson at the Cancer Prevention Institute of California, Hoda Anton-Culver, Argyrios Ziogas, and Hannah Park at the University of California Irvine, and Fred Schumacher at Case Western University. DIETCOMPLYF thanks the patients, nurses and clinical staff involved in the study. The DietCompLyf study was funded by the charity Against Breast Cancer (Registered Charity Number 1121258) and the NCRN. We thank the participants and the investigators of EPIC (European Prospective Investigation into Cancer and Nutrition). ESTHER thanks Hartwig Ziegler, Sonja Wolf, Volker Hermann, Christa Stegmaier, Katja Butterbach. FHRISK thanks NIHR for funding. GC-HBOC thanks Stefanie Engert, Heide Hellebrand, Sandra Kröber and LIFE - Leipzig Research Centre for Civilization Diseases (Markus Loeffler, Joachim Thiery, Matthias Nüchter, Ronny Baber). The GENICA Network: Dr. Margarete Fischer-Bosch-Institute of Clinical Pharmacology, Stuttgart, and University of Tübingen, Germany [HB, WYL], German Cancer Consortium (DKTK) and German Cancer Research Center (DKFZ) [HB], Deutsche Forschungsgemeinschaft (DFG, German Research Foundation) under Germany’s Excellence Strategy - EXC 2180 - 390900677 [HB], Department of Internal Medicine, Evangelische Kliniken Bonn gGmbH, Johanniter Krankenhaus, Bonn, Germany [YDK, Christian Baisch], Institute of Pathology, University of Bonn, Germany [Hans-Peter Fischer], Molecular Genetics of Breast Cancer, Deutsches Krebsforschungszentrum (DKFZ), Heidelberg, Germany [Ute Hamann], Institute for Prevention and Occupational Medicine of the German Social Accident Insurance, Institute of the Ruhr University Bochum (IPA), Bochum, Germany [TB, Beate Pesch, Sylvia Rabstein, Anne Lotz]; and Institute of Occupational Medicine and Maritime Medicine, University Medical Center Hamburg-Eppendorf, Germany [Volker Harth]. HABCS thanks Michael Bremer. HEBCS thanks Johanna Kiiski, Rainer Fagerholm, Kirsimari Aaltonen, Karl von Smitten, Irja Erkkilä. HMBCS thanks Peter Hillemanns, Hans Christiansen and Johann H. Karstens. HUBCS thanks Shamil Gantsev. KARMA and SASBAC thank the Swedish Medical Research Counsel. KBCP thanks Eija Myöhänen, Helena Kemiläinen. kConFab/AOCS wish to thank Heather Thorne, Eveline Niedermayr, all the kConFab research nurses and staff, the heads and staff of the Family Cancer Clinics, and the Clinical Follow Up Study (which has received funding from the NHMRC, the National Breast Cancer Foundation, Cancer Australia, and the National Institute of Health (USA)) for their contributions to this resource, and the many families who contribute to kConFab. LMBC thanks Gilian Peuteman, Thomas Van Brussel, EvyVanderheyden and Kathleen Corthouts. MABCS thanks Emilija Lazarova (Clinic of Radiotherapy and Oncology), Dzengis Jasar, Mitko Karadjozov (Adzibadem-Sistina” Hospital), Andrej Arsovski and Liljana Stojanovska (Re-Medika” Hospital) for their contributions and commitment to this study. MARIE thanks Petra Seibold, Dieter Flesch-Janys, Judith Heinz, Nadia Obi, Alina Vrieling, Sabine Behrens, Ursula Eilber, Muhabbet Celik, Til Olchers and Stefan Nickels. MBCSG (Milan Breast Cancer Study Group): Irene Feroce, Aliana Guerrieri Gonzaga, Monica Marabelli and and the personnel of the Cogentech Cancer Genetic Test Laboratory. The MCCS was made possible by the contribution of many people, including the original investigators, the teams that recruited the participants and continue working on follow-up, and the many thousands of Melbourne residents who continue to participate in the study. We thank the coordinators, the research staff and especially the MMHS participants for their continued collaboration on research studies in breast cancer. MSKCC thanks Marina Corines, Lauren Jacobs. MTLGEBCS would like to thank Martine Tranchant (CHU de Québec – Université Laval Research Center), Marie-France Valois, Annie Turgeon and Lea Heguy (McGill University Health Center, Royal Victoria Hospital; McGill University) for DNA extraction, sample management and skilful technical assistance. J.S. is Chair holder of the Canada Research Chair in Oncogenetics. The following are NBCS Collaborators: Kristine K. Sahlberg (PhD), Lars Ottestad (MD), Rolf Kåresen (Prof. Em.) Dr. Ellen Schlichting (MD), Marit Muri Holmen (MD), Toril Sauer (MD), Vilde Haakensen (MD), Olav Engebråten (MD), Bjørn Naume (MD), Alexander Fosså (MD), Cecile E. Kiserud (MD), Kristin V. Reinertsen (MD), Åslaug Helland (MD), Margit Riis (MD), Jürgen Geisler (MD), OSBREAC and Grethe I. Grenaker Alnæs (MSc). NBHS and For NHS and NHS2 the study protocol was approved by the institutional review boards of the Brigham and Women’s Hospital and Harvard T.H. Chan School of Public Health, and those of participating registries as required. We would like to thank the participants and staff of the NHS and NHS2 for their valuable contributions as well as the following state cancer registries for their help: AL, AZ, AR, CA, CO, CT, DE, FL, GA, ID, IL, IN, IA, KY, LA, ME, MD, MA, MI, NE, NH, NJ, NY, NC, ND, OH, OK, OR, PA, RI, SC, TN, TX, VA, WA, WY. The authors assume full responsibility for analyses and interpretation of these data. OBCS thanks Arja Jukkola-Vuorinen, Mervi Grip, Saila Kauppila, Meeri Otsukka, Leena Keskitalo and Kari Mononen for their contributions to this study. OFBCR thanks Teresa Selander, Nayana Weerasooriya. ORIGO thanks E. Krol-Warmerdam, and J. Blom for patient accrual, administering questionnaires, and managing clinical information. The LUMC survival data were retrieved from the Leiden hospital-based cancer registry system (ONCDOC) with the help of Dr. J. Molenaar. PBCS thanks Louise Brinton, Mark Sherman, Neonila Szeszenia-Dabrowska, Beata Peplonska, Witold Zatonski, Pei Chao, Michael Stagner. The ethical approval for the POSH study is MREC /00/6/69, UKCRN ID: 1137. We thank staff in the Experimental Cancer Medicine Centre (ECMC) supported Faculty of Medicine Tissue Bank and the Faculty of Medicine DNA Banking resource. PREFACE thanks Sonja Oeser and Silke Landrith. PROCAS thanks NIHR for funding. RBCS thanks Jannet Blom, Saskia Pelders, Annette Heemskerk and the Erasmus MC Family Cancer Clinic. SBCS thanks Sue Higham, Helen Cramp, Dan Connley, Ian Brock, Sabapathy Balasubramanian and Malcolm W.R. Reed. We thank the SEARCH and EPIC teams. SGBCC thanks the participants and research coordinator Ms Tan Siew Li. SKKDKFZS thanks all study participants, clinicians, family doctors, researchers and technicians for their contributions and commitment to this study. We thank the SUCCESS Study teams in Munich, Duessldorf, Erlangen and Ulm. SZBCS thanks Ewa Putresza. UCIBCS thanks Irene Masunaka. UKBGS thanks Breast Cancer Now and the Institute of Cancer Research for support and funding of the Breakthrough Generations Study, and the study participants, study staff, and the doctors, nurses and other health care providers and health information sources who have contributed to the study. We acknowledge NHS funding to the Royal Marsden/ICR NIHR Biomedical Research Centre. DGE, and AH, are supported by the all Manchester NIHR Biomedical Research Centre (IS-BRC-1215-20007). The authors thank the WHI investigators and staff for their dedication and the study participants for making the program possible. Support for title page creation and format was provided by AuthorArranger, a tool developed at the National Cancer Institute.

## Members of consortia listed as authors

### kConFab/AOCS Investigators

Stephen Fox, Ian Campbell (Peter MacCallum Cancer Centre, Melbourne, Australia); Georgia Chenevix-Trench, Amanda Spurdle, Penny Webb (QIMR Berghofer Medical Research Institute, Brisbane, Australia); Anna de Fazio (Westmead Millenium Institute, Sydney, Australia); Margaret Tassell (BCNA delegate, Community Representative); Judy Kirk (Westmead Hospital, Sydney, Australia); Geoff Lindeman (Walter and Eliza Hall Institute, Melbourne, Australia); Melanie Price (University of Sydney, Sydney, Australia); Melissa Southey (University of Melbourne, Melbourne, Australia); Roger Milne (Cancer Council Victoria, Melbourne, Australia); Sid Deb (Melbourne Health, Melbourne, Australia); David Bowtell (Garvan Institute of Medical Research, Sydney, Australia).

### ABCTB Investigators

Christine Clarke (Westmead Institute for Medical Research, University of Sydney, NSW, Australia); Rosemary Balleine (Pathology West ICPMR, Westmead, NSW, Australia); Robert Baxter (Kolling Institute of Medical Research, University of Sydney, Royal North Shore Hospital, NSW, Australia); Stephen Braye (Pathology North, John Hunter Hospital, Newcastle, NSW, 2305, Australia); Jane Carpenter (Westmead Institute for Medical Research, University of Sydney); Jane Dahlstrom (Department of Anatomical Pathology, ACT Pathology, Canberra Hospital, ACT, Australia; ANU Medical School, Australian National University, ACT, Australia); John Forbes (Department of Surgical Oncology, Calvary Mater Newcastle Hospital, Australian New Zealand Breast Cancer Trials Group, and School of Medicine and Public Health, University of Newcastle, NSW, Australia); C Soon Lee (School of Science and Health, The University of Western Sydney, Sydney, Australia); Deborah Marsh (Hormones and Cancer Group, Kolling Institute of Medical Research, Royal North Shore Hospital, University of Sydney, NSW, Australia); Adrienne Morey (SydPath St Vincent’s Hospital, Sydney, NSW, Australia); Nirmala Pathmanathan (Department of Tissue Pathology and Diagnostic Oncology, Pathology West; Westmead Breast Cancer Institute, Westmead Hospital, NSW, Australia); Rodney Scott (Centre for Information Based Medicine, Hunter Medical Research Institute, NSW, 2305, Australia; Priority Research Centre for Cancer, School of Biomedical Sciences and Pharmacy, Faculty of Health, University of Newcastle, NSW, Australia); Peter Simpson (The University of Queensland: UQ Centre for Clinical Research and School of Medicine, QLD, Australia); Allan Spigelman (Hereditary Cancer Clinic, St Vincent’s Hospital, The Kinghorn Cancer Centre, Sydney, New South Wales, 2010, Australia); Nicholas Wilcken (Crown Princess Mary Cancer Centre, Westmead Hospital, Westmead, Australia; Sydney Medical School - Westmead, University of Sydney, NSW, Australia); Desmond Yip (Department of Medical Oncology, The Canberra Hospital, ACT, Australia; ANU Medical School, Australian National University, ACT, Australia); Nikolajs Zeps (St John of God Perth Northern Hospitals, Perth, WA, Australia).

## CIMBA Funding and Acknowledgements

### Funding

CIMBA: The CIMBA data management and data analysis were supported by Cancer Research – UK grants C12292/A20861, C12292/A11174. ACA is a Cancer Research -UK Senior Cancer Research Fellow. GCT and ABS are NHMRC Research Fellows. iCOGS: the European Community’s Seventh Framework Programme under grant agreement n° 223175 (HEALTH-F2-2009-223175) (COGS), Cancer Research UK (C1287/A10118, C1287/A 10710, C12292/A11174, C1281/A12014, C5047/A8384, C5047/A15007, C5047/A10692, C8197/A16565), the National Institutes of Health (CA128978) and Post-Cancer GWAS initiative (1U19 CA148537, 1U19 CA148065 and 1U19 CA148112 - the GAME-ON initiative), the Department of Defence (W81XWH-10-1-0341), the Canadian Institutes of Health Research (CIHR) for the CIHR Team in Familial Risks of Breast Cancer (CRN-87521), and the Ministry of Economic Development, Innovation and Export Trade (PSR-SIIRI-701), Komen Foundation for the Cure, the Breast Cancer Research Foundation, and the Ovarian Cancer Research Fund. The PERSPECTIVE project was supported by the Government of Canada through Genome Canada and the Canadian Institutes of Health Research, the Ministry of Economy, Science and Innovation through Genome Québec, and The Quebec Breast Cancer Foundation.

BCFR: UM1 CA164920 from the National Cancer Institute. The content of this manuscript does not necessarily reflect the views or policies of the National Cancer Institute or any of the collaborating centers in the Breast Cancer Family Registry (BCFR), nor does mention of trade names, commercial products, or organizations imply endorsement by the US Government or the BCFR. BFBOCC: Lithuania (BFBOCC-LT): Research Council of Lithuania grant SEN-18/2015. BIDMC: Breast Cancer Research Foundation. BMBSA: Cancer Association of South Africa (PI Elizabeth J. van Rensburg). CNIO: Spanish Ministry of Health PI16/00440 supported by FEDER funds, the Spanish Ministry of Economy and Competitiveness (MINECO) SAF2014-57680-R and the Spanish Research Network on Rare diseases (CIBERER). COH-CCGCRN: Research reported in this publication was supported by the National Cancer Institute of the National Institutes of Health under grant number R25CA112486, and RC4CA153828 (PI: J. Weitzel) from the National Cancer Institute and the Office of the Director, National Institutes of Health. The content is solely the responsibility of the authors and does not necessarily represent the official views of the National Institutes of Health. CONSIT TEAM: Associazione Italiana Ricerca sul Cancro (AIRC; IG2015 no.16732) to P. Peterlongo. DEMOKRITOS: European Union (European Social Fund – ESF) and Greek national funds through the Operational Program “Education and Lifelong Learning” of the National Strategic Reference Framework (NSRF) - Research Funding Program of the General Secretariat for Research & Technology: SYN11_10_19 NBCA. Investing in knowledge society through the European Social Fund. DFKZ: German Cancer Research Center. EMBRACE: Cancer Research UK Grants C1287/A10118 and C1287/A11990. D. Gareth Evans and Fiona Lalloo are supported by an NIHR grant to the Biomedical Research Centre, Manchester. The Investigators at The Institute of Cancer Research and The Royal Marsden NHS Foundation Trust are supported by an NIHR grant to the Biomedical Research Centre at The Institute of Cancer Research and The Royal Marsden NHS Foundation Trust. Ros Eeles and Elizabeth Bancroft are supported by Cancer Research UK Grant C5047/A8385. Ros Eeles is also supported by NIHR support to the Biomedical Research Centre at The Institute of Cancer Research and The Royal Marsden NHS Foundation Trust. FCCC: The University of Kansas Cancer Center (P30 CA168524) and the Kansas Bioscience Authority Eminent Scholar Program. A.K.G. was funded by R0 1CA140323, R01 CA214545, and by the Chancellors Distinguished Chair in Biomedical Sciences Professorship. A.Vega is supported by the Spanish Health Research Foundation, Instituto de Salud Carlos III (ISCIII), partially supported by FEDER funds through Research Activity Intensification Program (contract grant numbers: INT15/00070, INT16/00154, INT17/00133), and through Centro de Investigación Biomédica en Red de Enferemdades Raras CIBERER (ACCI 2016: ER17P1AC7112/2018); Autonomous Government of Galicia (Consolidation and structuring program: IN607B), and by the Fundación Mutua Madrileña. GC-HBOC: German Cancer Aid (grant no 110837, Rita K. Schmutzler) and the European Regional Development Fund and Free State of Saxony, Germany (LIFE - Leipzig Research Centre for Civilization Diseases, project numbers 713-241202, 713-241202, 14505/2470, 14575/2470). GEMO: Ligue Nationale Contre le Cancer; the Association “Le cancer du sein, parlons-en!” Award, the Canadian Institutes of Health Research for the “CIHR Team in Familial Risks of Breast Cancer” program, the Fondation ARC pour la recherche sur le cancer (grant PJA 20151203365) and the French National Institute of Cancer (INCa grants AOR 01 082, 2001-2003, 2013-1-BCB-01-ICH-1 and SHS-E-SP 18-015). GEORGETOWN: the Non-Therapeutic Subject Registry Shared Resource at Georgetown University (NIH/NCI grant P30-CA051008), the Fisher Center for Hereditary Cancer and Clinical Genomics Research, and Swing Fore the Cure. G-FAST: Bruce Poppe is a senior clinical investigator of FWO. Mattias Van Heetvelde obtained funding from IWT. HCSC: Spanish Ministry of Health PI15/00059, PI16/01292, and CB-161200301 CIBERONC from ISCIII (Spain), partially supported by European Regional Development FEDER funds. HEBCS: Helsinki University Hospital Research Fund, the Finnish Cancer Society and the Sigrid Juselius Foundation. HEBON: the Dutch Cancer Society grants NKI1998-1854, NKI2004-3088, NKI2007-3756, the Netherlands Organization of Scientific Research grant NWO 91109024, the Pink Ribbon grants 110005 and 2014-187.WO76, the BBMRI grant NWO 184.021.007/CP46 and the Transcan grant JTC 2012 Cancer 12-054. HEBON thanks the registration teams of Dutch Cancer Registry (IKNL; S. Siesling, J. Verloop) and the Dutch Pathology database (PALGA; L. Overbeek) for part of the data collection. HUNBOCS: Hungarian Research Grants KTIA-OTKA CK-80745 and NKFI_OTKA K-112228. ICO: The authors would like to particularly acknowledge the support of the Asociación Española Contra el Cáncer (AECC), the Instituto de Salud Carlos III (organismo adscrito al Ministerio de Economía y Competitividad) and “Fondo Europeo de Desarrollo Regional (FEDER), una manera de hacer Europa” (PI10/01422, PI13/00285, PIE13/00022, PI15/00854, PI16/00563 and CIBERONC) and the Institut Català de la Salut and Autonomous Government of Catalonia (2009SGR290, 2014SGR338 and PERIS Project MedPerCan). IHCC: PBZ_KBN_122/P05/2004. INHERIT: Canadian Institutes of Health Research for the “CIHR Team in Familial Risks of Breast Cancer” program – grant # CRN-87521 and the Ministry of Economic Development, Innovation and Export Trade – grant # PSR-SIIRI-701. IOVHBOCS: Ministero della Salute and “5×1000” Istituto Oncologico Veneto grant. IPOBCS: Liga Portuguesa Contra o Cancro. kConFab: The National Breast Cancer Foundation, and previously by the National Health and Medical Research Council (NHMRC), the Queensland Cancer Fund, the Cancer Councils of New South Wales, Victoria, Tasmania and South Australia, and the Cancer Foundation of Western Australia. MAYO: NIH grants CA116167, CA192393 and CA176785, an NCI Specialized Program of Research Excellence (SPORE) in Breast Cancer (CA116201),and a grant from the Breast Cancer Research Foundation. MCGILL: Jewish General Hospital Weekend to End Breast Cancer, Quebec Ministry of Economic Development, Innovation and Export Trade. Marc Tischkowitz is supported by the funded by the European Union Seventh Framework Program (2007Y2013)/European Research Council (Grant No. 310018). MODSQUAD: MH CZ - DRO (MMCI, 00209805), **MEYS - NPS I - LO1413** to LF, and by Charles University in Prague project UNCE204024 (MZ). MSKCC: the Breast Cancer Research Foundation, the Robert and Kate Niehaus Clinical Cancer Genetics Initiative, the Andrew Sabin Research Fund and a Cancer Center Support Grant/Core Grant (P30 CA008748). NAROD: 1R01 CA149429-01. NCI: the Intramural Research Program of the US National Cancer Institute, NIH, and by support services contracts NO2-CP-11019-50, N02-CP-21013-63 and N02-CP-65504 with Westat, Inc, Rockville, MD. NICCC: Clalit Health Services in Israel, the Israel Cancer Association and the Breast Cancer Research Foundation (BCRF), NY. NNPIO: the Russian Foundation for Basic Research (grants 17-00-00171, 18-515-45012 and 19-515-25001). NRG Oncology: U10 CA180868, NRG SDMC grant U10 CA180822, NRG Administrative Office and the NRG Tissue Bank (CA 27469), the NRG Statistical and Data Center (CA 37517) and the Intramural Research Program, NCI, KAP is an Australian National Breast Cancer Foundation Fellow. OSUCCG: Ohio State University Comprehensive Cancer Center. PBCS: Italian Association of Cancer Research (AIRC) [IG 2013 N.14477] and Tuscany Institute for Tumors (ITT) grant 2014-2015-2016. SMC: the Israeli Cancer Association. SWE-BRCA: the Swedish Cancer Society. UCHICAGO: NCI Specialized Program of Research Excellence (SPORE) in Breast Cancer (CA125183), R01 CA142996, 1U01CA161032 and by the Ralph and Marion Falk Medical Research Trust, the Entertainment Industry Fund National Women’s Cancer Research Alliance and the Breast Cancer research Foundation. OIO is an ACS Clinical Research Professor. UCSF: UCSF Cancer Risk Program and Helen Diller Family Comprehensive Cancer Center. UKFOCR: Cancer Researc h UK. UPENN: Breast Cancer Research Foundation (to SMD, KLN); Susan G. Komen Foundation for the cure (SMD), Basser Research Center for BRCA (SMD, KLN). UPITT/MWH: Hackers for Hope Pittsburgh. VFCTG: Victorian Cancer Agency, Cancer Australia, National Breast Cancer Foundation. WCP: Dr Karlan is funded by the American Cancer Society Early Detection Professorship (SIOP-06-258-01-COUN) and the National Center for Advancing Translational Sciences (NCATS), Grant UL1TR000124. Tracy A. O’Mara was supported by NHMRC Early Career Research Fellow.

### Acknowledgements

All the families and clinicians who contribute to the studies; Catherine M. Phelan for her contribution to CIMBA until she passed away on 22 September 2017; Sue Healey, in particular taking on the task of mutation classification with the late Olga Sinilnikova; Maggie Angelakos, Judi Maskiell, Gillian Dite, Helen Tsimiklis; members and participants in the New York site of the Breast Cancer Family Registry; members and participants in the Ontario Familial Breast Cancer Registry; Vilius Rudaitis and Laimonas Griškevičius; Drs Janis Eglitis, Anna Krilova and Aivars Stengrevics; Yuan Chun Ding and Linda Steele for their work in participant enrollment and biospecimen and data management; Bent Ejlertsen and Anne-Marie Gerdes for the recruitment and genetic counseling of participants; Alicia Barroso, Rosario Alonso and Guillermo Pita; all the individuals and the researchers who took part in CONSIT TEAM (Consorzio Italiano Tumori Ereditari Alla Mammella), in particular: Dario Zimbalatti, Daniela Zaffaroni, Laura Ottini, Giuseppe Giannini, Liliana Varesco, Viviana Gismondi, Maria Grazia Tibiletti, Daniela Furlan, Antonella Savarese, Aline Martayan, Stefania Tommasi, Brunella Pilato and the personnel of the Cogentech Cancer Genetic Test Laboratory, Milan, Italy. Ms. JoEllen Weaver and Dr. Betsy Bove; FPGMX: members of the Cancer Genetics group (IDIS): Ana Blanco, Miguel Aguado, Uxía Esperón and Belinda Rodríguez.; IFE - Leipzig Research Centre for Civilization Diseases (Markus Loeffler, Joachim Thiery, Matthias Nüchter, Ronny Baber); We thank all participants, clinicians, family doctors, researchers, and technicians for their contributions and commitment to the DKFZ study and the collaborating groups in Lahore, Pakistan (Noor Muhammad, Sidra Gull, Seerat Bajwa, Faiz Ali Khan, Humaira Naeemi, Saima Faisal, Asif Loya, Mohammed Aasim Yusuf) and Bogota, Colombia (Diana Torres, Ignacio Briceno, Fabian Gil). Genetic Modifiers of Cancer Risk in BRCA1/2 Mutation Carriers (GEMO) study is a study from the National Cancer Genetics Network UNICANCER Genetic Group, France. We wish to pay a tribute to Olga M. Sinilnikova, who with Dominique Stoppa-Lyonnet initiated and coordinated GEMO until she sadly passed away on the 30th June 2014. The team in Lyon (Olga Sinilnikova, Mélanie Léoné, Laure Barjhoux, Carole Verny-Pierre, Sylvie Mazoyer, Francesca Damiola, Valérie Sornin) managed the GEMO samples until the biological resource centre was transferred to Paris in December 2015 (Noura Mebirouk, Fabienne Lesueur, Dominique Stoppa-Lyonnet). We want to thank all the GEMO collaborating groups for their contribution to this study: Coordinating Centre, Service de Génétique, Institut Curie, Paris, France: Muriel Belotti, Ophélie Bertrand, Anne-Marie Birot, Bruno Buecher, Sandrine Caputo, Anaïs Dupré, Emmanuelle Fourme, Marion Gauthier-Villars, Lisa Golmard, Claude Houdayer, Marine Le Mentec, Virginie Moncoutier, Antoine de Pauw, Claire Saule, Dominique Stoppa-Lyonnet, and Inserm U900, Institut Curie, Paris, France: Fabienne Lesueur, Noura Mebirouk.Contributing Centres: Unité Mixte de Génétique Constitutionnelle des Cancers Fréquents, Hospices Civils de Lyon - Centre Léon Bérard, Lyon, France: Nadia Boutry-Kryza, Alain Calender, Sophie Giraud, Mélanie Léone. Institut Gustave Roussy, Villejuif, France: Brigitte Bressac-de-Paillerets, Olivier Caron, Marine Guillaud-Bataille. Centre Jean Perrin, Clermont–Ferrand, France: Yves-Jean Bignon, Nancy Uhrhammer. Centre Léon Bérard, Lyon, France: Valérie Bonadona, Christine Lasset. Centre François Baclesse, Caen, France: Pascaline Berthet, Laurent Castera, Dominique Vaur. Institut Paoli Calmettes, Marseille, France: Violaine Bourdon, Catherine Noguès, Tetsuro Noguchi, Cornel Popovici, Audrey Remenieras, Hagay Sobol. CHU Arnaud-de-Villeneuve, Montpellier, France: Isabelle Coupier, Pascal Pujol. Centre Oscar Lambret, Lille, France: Claude Adenis, Aurélie Dumont, Françoise Révillion. Centre Paul Strauss, Strasbourg, France: Danièle Muller. Institut Bergonié, Bordeaux, France: Emmanuelle Barouk-Simonet, Françoise Bonnet, Virginie Bubien, Michel Longy, Nicolas Sevenet, Institut Claudius Regaud, Toulouse, France: Laurence Gladieff, Rosine Guimbaud, Viviane Feillel, Christine Toulas. CHU Grenoble, France: Hélène Dreyfus, Christine Dominique Leroux, Magalie Peysselon, Rebischung. CHU Dijon, France: Amandine Baurand, Geoffrey Bertolone, Fanny Coron, Laurence Faivre, Caroline Jacquot, Sarab Lizard. CHU St-Etienne, France: Caroline Kientz, Marine Lebrun, Fabienne Prieur. Hôtel Dieu Centre Hospitalier, Chambéry, France: Sandra Fert Ferrer. Centre Antoine Lacassagne, Nice, France: Véronique Mari. CHU Limoges, France: Laurence Vénat-Bouvet. CHU Nantes, France: Stéphane Bézieau, Capucine Delnatte. CHU Bretonneau, Tours and Centre Hospitalier de Bourges France: Isabelle Mortemousque. Groupe Hospitalier Pitié-Salpétrière, Paris, France: Chrystelle Colas, Florence Coulet, Florent Soubrier, Mathilde Warcoin. CHU Vandoeuvre-les-Nancy, France: Myriam Bronner, Johanna Sokolowska. CHU Besançon, France: Marie-Agnès Collonge-Rame, Alexandre Damette. CHU Poitiers, Centre Hospitalier d’Angoulême and Centre Hospitalier de Niort, France: Paul Gesta. Centre Hospitalier de La Rochelle: Hakima Lallaoui. CHU Nîmes Carémeau, France: Jean Chiesa. CHI Poissy, France: Denise Molina-Gomes. CHU Angers, France: Olivier Ingster; Ilse Coene en Brecht Crombez; Ilse Coene and Brecht Crombez; Alicia Tosar and Paula Diaque; Irja Erkkilä and Virpi Palola; The Hereditary Breast and Ovarian Cancer Research Group Netherlands (HEBON) consists of the following Collaborating Centers: Coordinating center: Netherlands Cancer Institute, Amsterdam, NL: M.A. Rookus, F.B.L. Hogervorst, F.E. van Leeuwen, S. Verhoef, M.K. Schmidt, N.S. Russell, D.J. Jenner; Erasmus Medical Center, Rotterdam, NL: J.M. Collée, A.M.W. van den Ouweland, M.J. Hooning, C. Seynaeve, C.H.M. van Deurzen, I.M. Obdeijn; Leiden University Medical Center, NL: C.J. van Asperen, J.T. Wijnen, R.A.E.M. Tollenaar, P. Devilee, T.C.T.E.F. van Cronenburg; Radboud University Nijmegen Medical Center, NL: C.M. Kets, A.R. Mensenkamp; University Medical Center Utrecht, NL: M.G.E.M. Ausems, R.B. van der Luijt, C.C. van der Pol; Amsterdam Medical Center, NL: C.M. Aalfs, T.A.M. van Os; VU University Medical Center, Amsterdam, NL: J.J.P. Gille, Q. Waisfisz, H.E.J. Meijers-Heijboer; University Hospital Maastricht, NL: E.B. Gómez-Garcia, M.J. Blok; University Medical Center Groningen, NL: J.C. Oosterwijk, A.H. van der Hout, M.J. Mourits, G.H. de Bock; The Netherlands Foundation for the detection of hereditary tumours, Leiden, NL: H.F. Vasen; The Netherlands Comprehensive Cancer Organization (IKNL): S. Siesling, J.Verloop; The Dutch Pathology Registry (PALGA): L.I.H. Overbeek; Hong Kong Sanatorium and Hospital; the Hungarian Breast and Ovarian Cancer Study Group members (Janos Papp, Aniko Bozsik, Timea Pocza, Zoltan Matrai, Miklos Kasler, Judit Franko, Maria Balogh, Gabriella Domokos, Judit Ferenczi, Department of Molecular Genetics, National Institute of Oncology, Budapest, Hungary) and the clinicians and patients for their contributions to this study; the Oncogenetics Group (VHIO) and the High Risk and Cancer Prevention Unit of the University Hospital Vall d’Hebron, Miguel Servet Progam (CP10/00617), and the Cellex Foundation for providing research facilities and equipment; the ICO Hereditary Cancer Program team led by Dr. Gabriel Capella; the ICO Hereditary Cancer Program team led by Dr. Gabriel Capella; Dr Martine Dumont for sample management and skillful assistance; Catarina Santos and Pedro Pinto; members of the Center of Molecular Diagnosis, Oncogenetics Department and Molecular Oncology Research Center of Barretos Cancer Hospital; Heather Thorne, Eveline Niedermayr, all the kConFab research nurses and staff, the heads and staff of the Family Cancer Clinics, and the Clinical Follow Up Study (which has received funding from the NHMRC, the National Breast Cancer Foundation, Cancer Australia, and the National Institute of Health (USA)) for their contributions to this resource, and the many families who contribute to kConFab; the KOBRA Study Group; (National Human Genome Research Institute, National Institutes of Health, Bethesda, MD, USA); Eva Machackova (Department of Cancer Epidemiology and Genetics, Masaryk Memorial Cancer Institute and MF MU, Brno, Czech Republic); and Michal Zikan, Petr Pohlreich and Zdenek Kleibl (Oncogynecologic Center and Department of Biochemistry and Experimental Oncology, First Faculty of Medicine, Charles University, Prague, Czech Republic); Anne Lincoln, Lauren Jacobs; the participants in Hereditary Breast/Ovarian Cancer Study and Breast Imaging Study for their selfless contributions to our research; the NICCC National Familial Cancer Consultation Service team led by Sara Dishon, the lab team led by Dr. Flavio Lejbkowicz, and the research field operations team led by Dr. Mila Pinchev; the investigators of the Australia New Zealand NRG Oncology group; members and participants in the Ontario Cancer Genetics Network; Kevin Sweet, Caroline Craven, Julia Cooper, Amber Aielts, and Michelle O’Conor; Yip Cheng Har, Nur Aishah Mohd Taib, Phuah Sze Yee, Norhashimah Hassan and all the research nurses, research assistants and doctors involved in the MyBrCa Study for assistance in patient recruitment, data collection and sample preparation, Philip Iau, Sng Jen-Hwei and Sharifah Nor Akmal for contributing samples from the Singapore Breast Cancer Study and the HUKM-HKL Study respectively; the Meirav Comprehensive breast cancer center team at the Sheba Medical Center; Christina Selkirk; Helena Jernström, Karin Henriksson, Katja Harbst, Maria Soller, Ulf Kristoffersson; from Gothenburg Sahlgrenska University Hospital: Anna Öfverholm, Margareta Nordling, Per Karlsson, Zakaria Einbeigi; from Stockholm and Karolinska University Hospital: Anna von Wachenfeldt, Annelie Liljegren, Brita Arver, Gisela Barbany Bustinza; from Umeå University Hospital: Beatrice Melin, Christina Edwinsdotter Ardnor, Monica Emanuelsson; from Uppsala University: Hans Ehrencrona, Maritta Hellström Pigg, Richard Rosenquist; from Linköping University Hospital: Marie Stenmark-Askmalm, Sigrun Liedgren; Cecilia Zvocec, Qun Niu; Joyce Seldon and Lorna Kwan; Dr. Robert Nussbaum, Beth Crawford, Kate Loranger, Julie Mak, Nicola Stewart, Robin Lee, Amie Blanco and Peggy Conrad and Salina Chan; Carole Pye, Patricia Harrington and Eva Wozniak; Geoffrey Lindeman, Marion Harris, Martin Delatycki, Sarah Sawyer, Rebecca Driessen, and Ella Thompson for performing all DNA amplification.

## Members of consortia listed as authors

### EMBRACE

Helen Gregory (North of Scotland Regional Genetics Service, NHS Grampian & University of Aberdeen, Foresterhill, Aberdeen, UK); Zosia Miedzybrodzka (North of Scotland Regional Genetics Service, NHS Grampian & University of Aberdeen, Foresterhill, Aberdeen, UK); Patrick J. Morrison (Northern Ireland Regional Genetics Centre, Belfast Health and Social Care Trust, and Department of Medical Genetics, Queens University Belfast, Belfast, UK); Kai-ren Ong (West Midlands Regional Genetics Service, Birmingham Women’s Hospital Healthcare NHS Trust, Edgbaston, Birmingham, UK); Alan Donaldson (Clinical Genetics Department, St Michael’s Hospital, Bristol, UK); Marc Tischkowitz (Department of Medical Genetics, University of Cambridge, UK); Mark T. Rogers (All Wales Medical Genetics Services, University Hospital of Wales, Cardiff, UK); M. John Kennedy (Academic Unit of Clinical and Molecular Oncology, Trinity College Dublin and St James’s Hospital, Dublin, Eire); Mary E. Porteous (South East of Scotland Regional Genetics Service, Western General Hospital, Edinburgh, UK); Carole Brewer (Department of Clinical Genetics, Royal Devon & Exeter Hospital, Exeter, UK); Rosemarie Davidson (Clinical Genetics, Southern General Hospital, Glasgow, UK); Louise Izatt (Clinical Genetics, Guy’s and St. Thomas’ NHS Foundation Trust, London, UK); Angela Brady (North West Thames Regional Genetics Service, Kennedy-Galton Centre, Harrow, UK); Julian Barwell (Leicestershire Clinical Genetics Service, University Hospitals of Leicester NHS Trust, UK); Julian Adlard (Yorkshire Regional Genetics Service, Leeds, UK); Claire Foo (Department of Clinical Genetics, Alder Hey Hospital, Eaton Road, Liverpool, UK); D. Gareth Evans (Genetic Medicine, Manchester Academic Health Sciences Centre, Central Manchester University Hospitals NHS Foundation Trust, Manchester, UK); Fiona Lalloo (Genetic Medicine, Manchester Academic Health Sciences Centre, Central Manchester University Hospitals NHS Foundation Trust, Manchester, UK); Lucy E. Side (North East Thames Regional Genetics Service, Great Ormond Street Hospital for Children NHS Trust, London, UK); Jacqueline Eason (Nottingham Clinical Genetics Service, Nottingham University Hospitals NHS Trust, UK); Alex Henderson (Institute of Genetic Medicine, Centre for Life, Newcastle Upon Tyne Hospitals NHS Trust, Newcastle upon Tyne, UK); Lisa Walker (Oxford Regional Genetics Service, Churchill Hospital, Oxford, UK); Rosalind A. Eeles (Oncogenetics Team, The Institute of Cancer Research and Royal Marsden NHS Foundation Trust, UK); Jackie Cook (Sheffield Clinical Genetics Service, Sheffield Children’s Hospital, Sheffield, UK); Katie Snape (South West Thames Regional Genetics Service, St.Georges Hospital, Cranmer Terrace, Tooting, London, UK); Diana Eccles (University of Southampton Faculty of Medicine, Southampton University Hospitals NHS Trust, Southampton, UK); Alex Murray (All Wales Medical Genetics Services, Singleton Hospital, Swansea, UK); Emma McCann (All Wales Medical Genetics Service, Glan Clwyd Hospital, Rhyl, UK).

### GEMO Study Collaborators

Dominique Stoppa-Lyonnet, Muriel Belotti, Anne-Marie Birot, Bruno Buecher, Emmanuelle Fourme, Marion Gauthier-Villars, Lisa Golmard, Claude Houdayer, Virginie Moncoutier, Antoine de Pauw, Claire Saule (Service de Génétique, Institut Curie, Paris, France); Fabienne Lesueur, Noura Mebirouk (Inserm U900, Institut Curie, Paris, France); Olga Sinilnikova†, Sylvie Mazoyer, Francesca Damiola, Laure Barjhoux, Carole Verny-Pierre, Mélanie Léone, Nadia Boutry-Kryza, Alain Calender, Sophie Giraud (Unité Mixte de Génétique Constitutionnelle des Cancers Fréquents, Hospices Civils de Lyon - Centre Léon Bérard, Lyon, France); Olivier Caron, Marine GuillaudBataille (Institut Gustave Roussy, Villejuif, France: Brigitte Bressac-de-Paillerets); YvesJean Bignon, Nancy Uhrhammer (Centre Jean Perrin, Clermont–Ferrand, France); Christine Lasset, Valérie Bonadona (Centre Léon Bérard, Lyon, France); Pascaline Berthet, Dominique Vaur, Laurent Castera (Centre François Baclesse, Caen, France); Hagay Sobol, Violaine Bourdon, Tetsuro Noguchi, Audrey Remenieras, François Eisinger, Catherine Noguès (Institut Paoli Calmettes, Marseille, France); Isabelle Coupier, Pascal Pujol (CHU Arnaud-de-Villeneuve, Montpellier, France); Jean-Philippe Peyrat, Joëlle Fournier, Françoise Révillion, Claude Adenis (Centre Oscar Lambret, Lille, France); Danièle Muller, Jean-Pierre Fricker (Centre Paul Strauss, Strasbourg, France); Emmanuelle Barouk-Simonet, Françoise Bonnet, Virginie Bubien, Nicolas Sevenet, Michel Longy (Institut Bergonié, Bordeaux, France); Christine Toulas, Rosine Guimbaud, Laurence Gladieff, Viviane Feillel (Institut Claudius Regaud, Toulouse, France); Dominique Leroux, Hélène Dreyfus, Christine Rebischung, Magalie Peysselon (CHU Grenoble, France); Fanny Coron, Laurence Faivre, Amandine Baurand, Caroline Jacquot, Geoffrey, Bertolone, Sarab Lizard (CHU Dijon, France); Fabienne Prieur, Marine Lebrun, Caroline Kientz (CHU St-Etienne, France); Sandra Fert Ferrer (Hôtel Dieu Centre Hospitalier, Chambéry, France); Véronique Mari (Centre Antoine Lacassagne, Nice, France); Laurence Vénat-Bouvet (CHU Limoges, France); Capucine Delnatte, Stéphane Bézieau (CHU Nantes, France); Isabelle Mortemousque (CHU Bretonneau, Tours and Centre Hospitalier de Bourges France); Florence Coulet, Chrystelle Colas, Florent Soubrier, Mathilde Warcoin (Groupe Hospitalier PitiéSalpétrière, Paris, France); Johanna Sokolowska, Myriam Bronner (CHU Vandoeuvreles-Nancy, France); Marie-Agnès Collonge-Rame, Alexandre Damette (CHU Besançon, France); Paul Gesta (CHU Poitiers, Centre Hospitalier d’Angoulême and Centre Hospitalier de Niort, France); Hakima Lallaoui (Centre Hospitalier de La Rochelle); Jean Chiesa (CHU Nîmes Carémeau, France); Denise Molina-Gomes (CHI Poissy, France); Olivier Ingster (CHU Angers, France).

