## supplementary Figure and Note for "Genome-wide association study identifies 32 novel breast cancer susceptibility loci from overall and subtype-specific analyses"

**Supplementary figure 1**. Overview of the analytic strategy and results from the investigation of breast cancer susceptibility single-nucleotide polymorphisms (SNPs) in women of European descent. Analyses included investigating for susceptibility SNPs for overall breast cancer (invasive, in-situ or unknown invasiveness) and for susceptibility SNPs accounting for tumor heterogeneity according to the estrogen receptor (ER), progesterone receptor (PR), human epidermal growth factor receptor 2 (HER2), and grade, and specifically investigating for SNPs that predispose for risk of the triple-negative (TN) subtype.

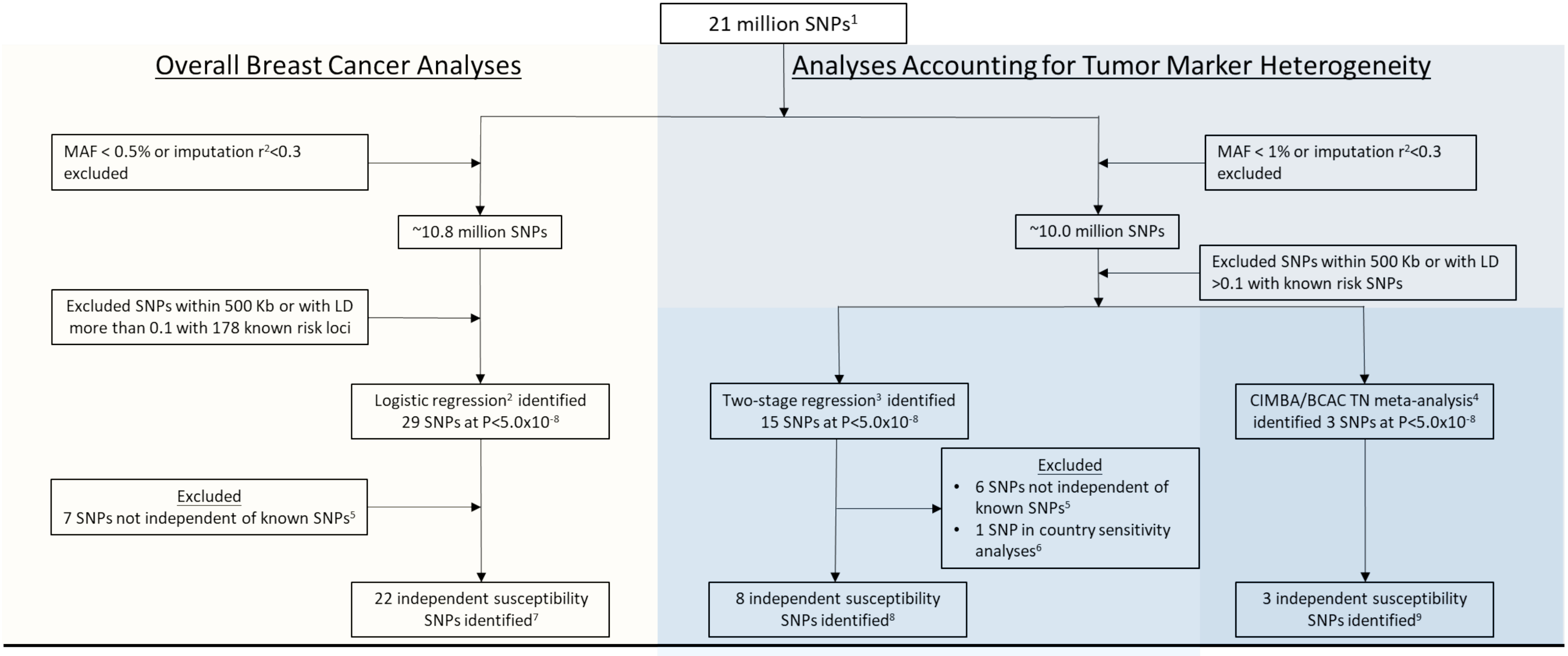


**(1)** Genotyping data from two Illumina genome-wide custom arrays, the iCOGS and Oncoarray, and imputed to the 1000 Genomes Project (Phase 3). **(2)** Overall breast cancer (invasive, in-situ, or unknown invasiveness) analyses included 82 studies from the Breast Cancer Association Consortium (BCAC; 118,474 cases and 96,201 controls) and summary level data from 11 other breast cancer GWAS (14,910 cases and 17,588 controls; **Supplementary Table 1**). **(3)** Analyses accounting for tumor marker heterogeneity according to ER, PR, HER2 and grade included 81 studies from BCAC (106,278 invasive cases and 91,477 controls). **(4)** Analyses investigating triple-negative susceptibility SNPs included 91,477 controls and 8,602 TN (effective sample, see **Supplementary Note**) cases from BCAC and 9,414 affected and 9,494 unaffected *BRCA1/2* carriers from 60 studies from the Consortium of Investigators of Modifiers of BRCA1/2 (CIMBA; **Supplementary Table 3**). **(5)** SNPs excluded following conditional analyses showing the identified SNPs to not be independent ($P>1x10^{-6}$) of 178 known susceptibility SNPs (see **Online Methods**). **(6)** See **Supplementary Figure 7** for results of country-specific sensitivity analyses. **(7)** See **Supplementary Table 5** for the 22 independent susceptibility SNPs identified in overall breast cancer analyses. **(8)** See **Supplementary Table 6** for the 8 independent susceptibility SNPs identified using two-stage polytomous regression, accounting for tumor markers heterogeneity according to ER, PR, HER2, and grade. **(9)** See **Supplementary Table 7** for the 3 independent susceptibility SNPs identified in the CIMBA / BCAC-TN meta-analysis. Note that rs78378222 was detected in both the analyses using the two-stage polytomous regression and in CIMBA / BCAC-TN.

**Supplementary figure 2**. SNP associations with overall breast cancer risk identified using standard logistic regression **a)** Manhattan plot showing -$\log_{10}P$ values for SNP associations with breast cancer risk. **b)** Manhattan plot after excluding previous known regions (Online Methods) **c)** Quantile-Quantile (Q-Q) plot of observed P-values versus expected P-values for all SNPs. **d)** QQ plot[1] after excluding previous known regions.

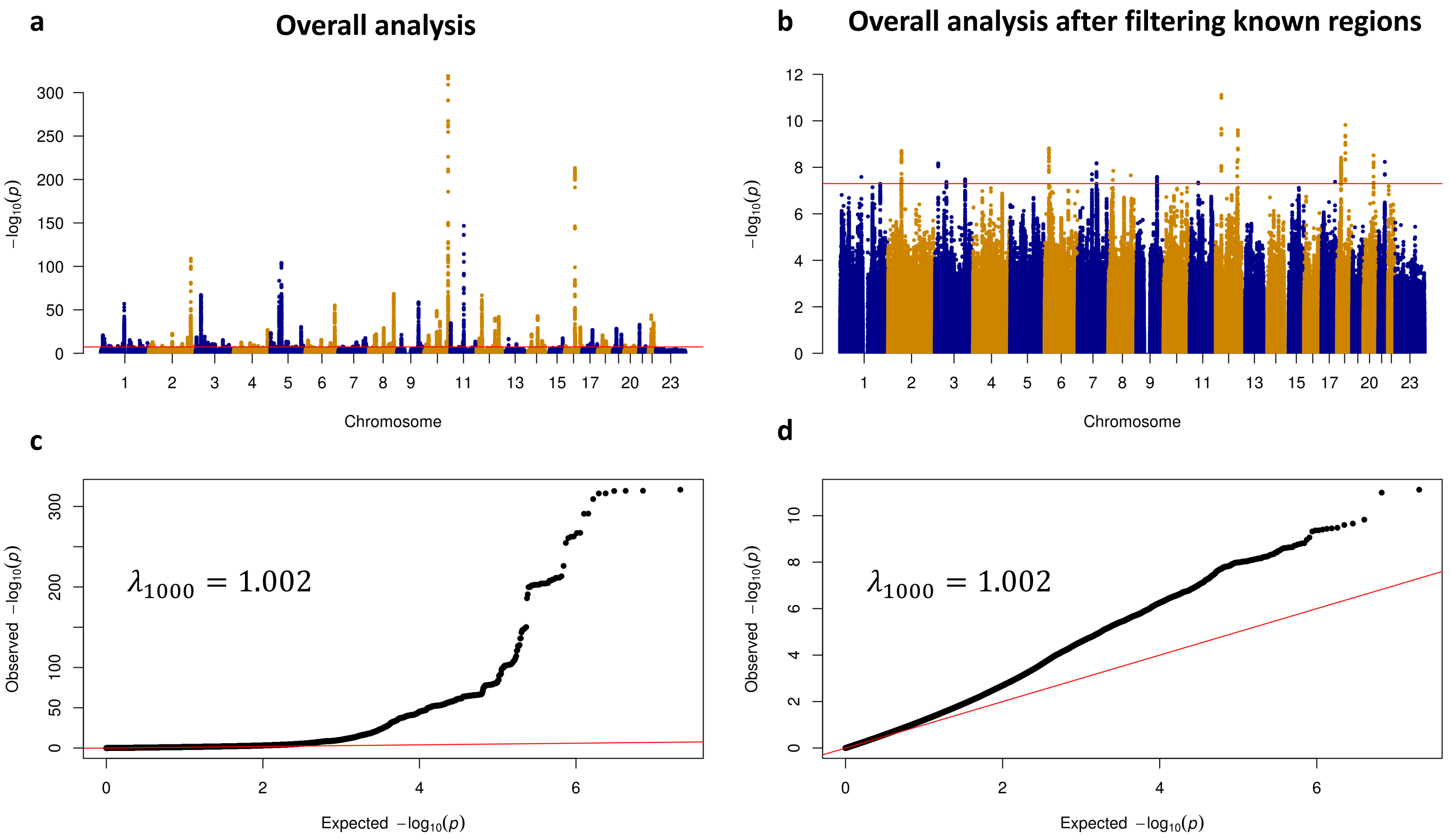


**1)** $\lambda_{1000}$ scale the genomic inflation factor $\lambda$ to a study with sample size of 1000 cases and 1000 controls using the formula $\lambda_{1000} = 1 + 500 * (\lambda - 1)/(\frac{1}{n_{cases}} + \frac{1}{n_{control}})$

**Supplementary figure 3**. SNP associations with breast cancer risk using a mixed-effect two-stage model (**Oline Methods**) accounting for tumor heterogeneity according to the ER, PR, HER2, and grade. **a)** Manhattan plot showing -log$_{10}P$ values for SNP associations with breast cancer risk. **b)** Manhattan plot showing -log$_{10}P$ values for SNP associations with breast cancer risk after excluding previously known regions (Online Methods) and 22 loci identified through standard logistic regression analysis (Supplementary Figure 2). **c)** QQ plot[1] of observed P-values versus expected P-values for all SNPs. **d)** QQ plot of observed P-values versus expected P-values for remaining SNPs after excluding previously known regions and 22 loci identified through standard logistic regression analysis.

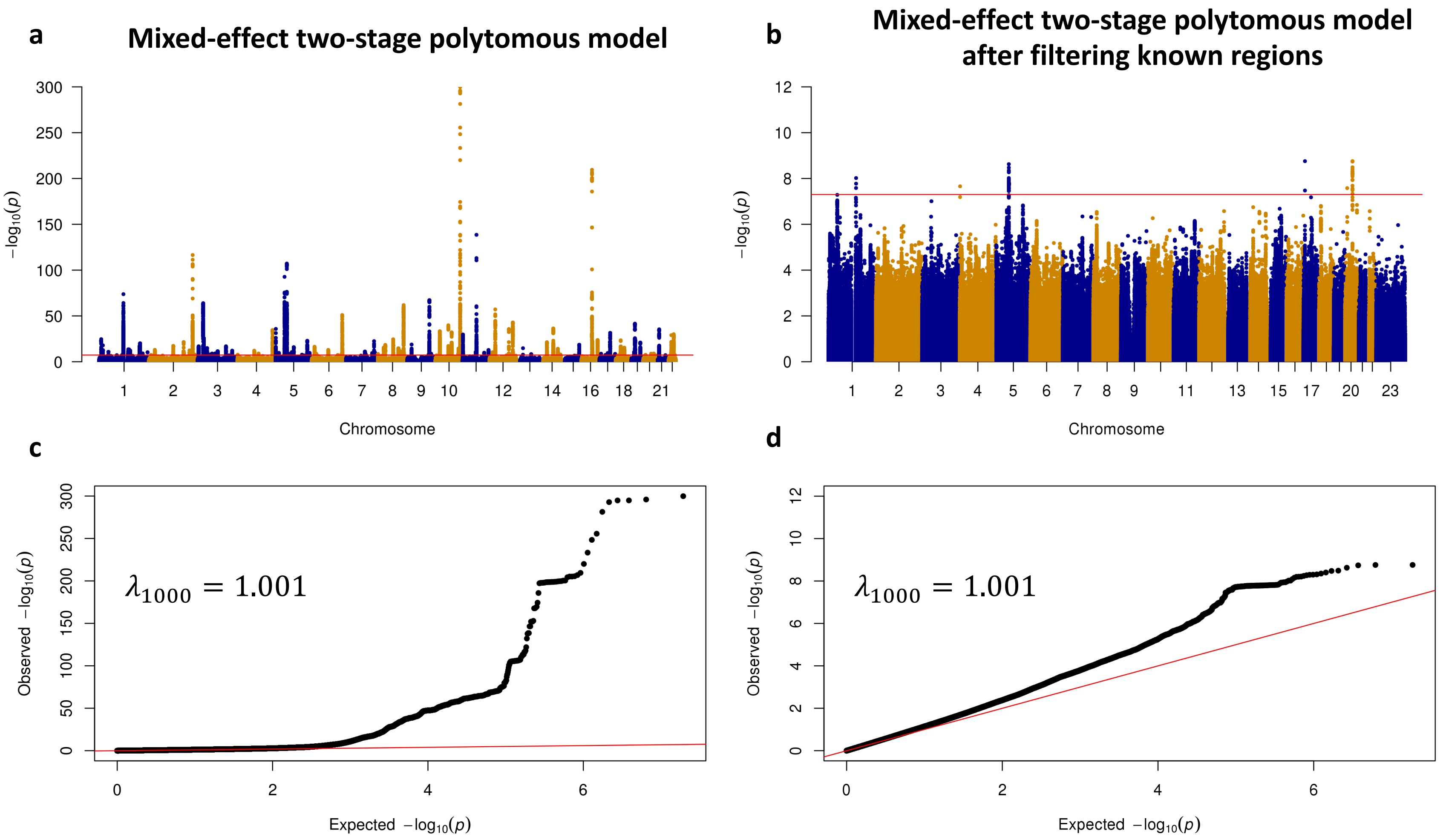


**1)** $\lambda_{1000}$ scale the genomic inflation factor $\lambda$ to a study with sample size of 1000 cases and 1000 controls using the formula $\lambda_{1000} = 1 + 500 * (\lambda - 1)/(\frac{1}{n_{cases}} + \frac{1}{n_{control}})$

**Supplementary figure 4.** SNP associations with breast cancer risk using a fixed-effect two-stage model (**Oline Methods**) accounting for tumor heterogeneity according to the ER, PR, HER2, and grade. **a)** Manhattan plot showing -$\log_{10}$P values for SNP associations with breast cancer risk. **b)** Manhattan plot showing -$\log_{10}$P values for SNP associations with breast cancer risk after excluding previously known regions (Online Methods) and 22 loci identified through standard logistic regression analysis (Supplementary Figure 2). **c)** QQ plot[1] of observed P-values versus expected P-values for all SNPs. **d)** QQ plot of observed P-values versus expected P-values for remaining SNPs after excluding previously known regions and 22 loci identified through standard analysis.

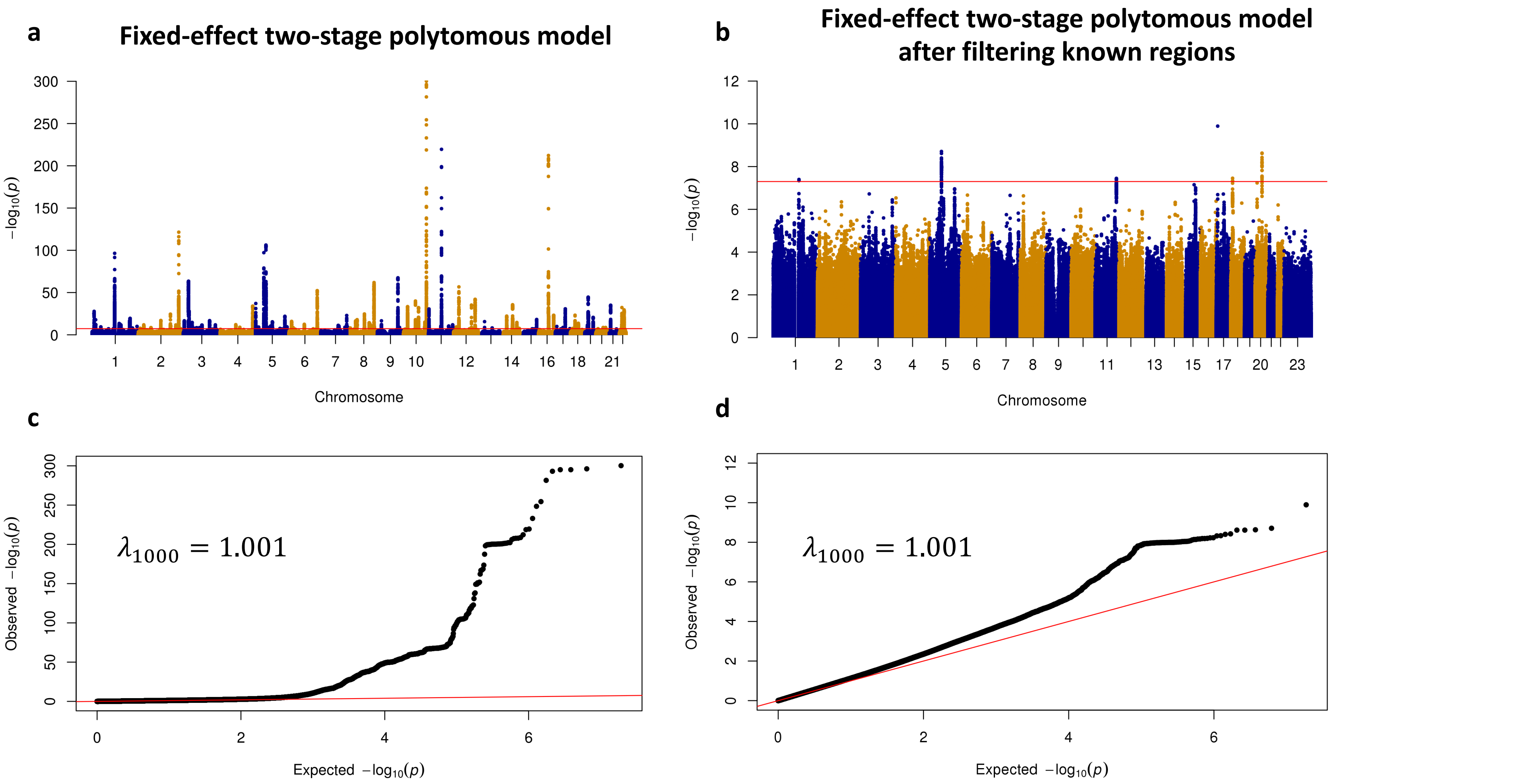


**1)** $\lambda_{1000}$ scale the genomic inflation factor $\lambda$ to a study with sample size of 1000 cases and 1000 controls using the formula $\lambda_{1000} = 1 + 500 * (\lambda - 1)/(\frac{1}{n_{cases}} + \frac{1}{n_{control}})$

**Supplementary figure 5**. SNP association with triple negative (TN) breast cancer risk using a fixed-effect meta-analysis of results between BCAC TN and CIMBA *BRCA1 carriers.* **a)** Manhattan plot showing -$\log_{10}$P values for SNP associations with TN breast cancer risk. **b)** Manhattan plot showing -$\log_{10}$P values for SNP associations with TN breast cancer risk after excluding previously known regions (Online Methods). **c)** QQ plot[1] of observed P-values versus expected P-values for all SNPs **d)** QQ plot of observed P-values versus expected P-values for remaining SNPs after excluding previously known regions.

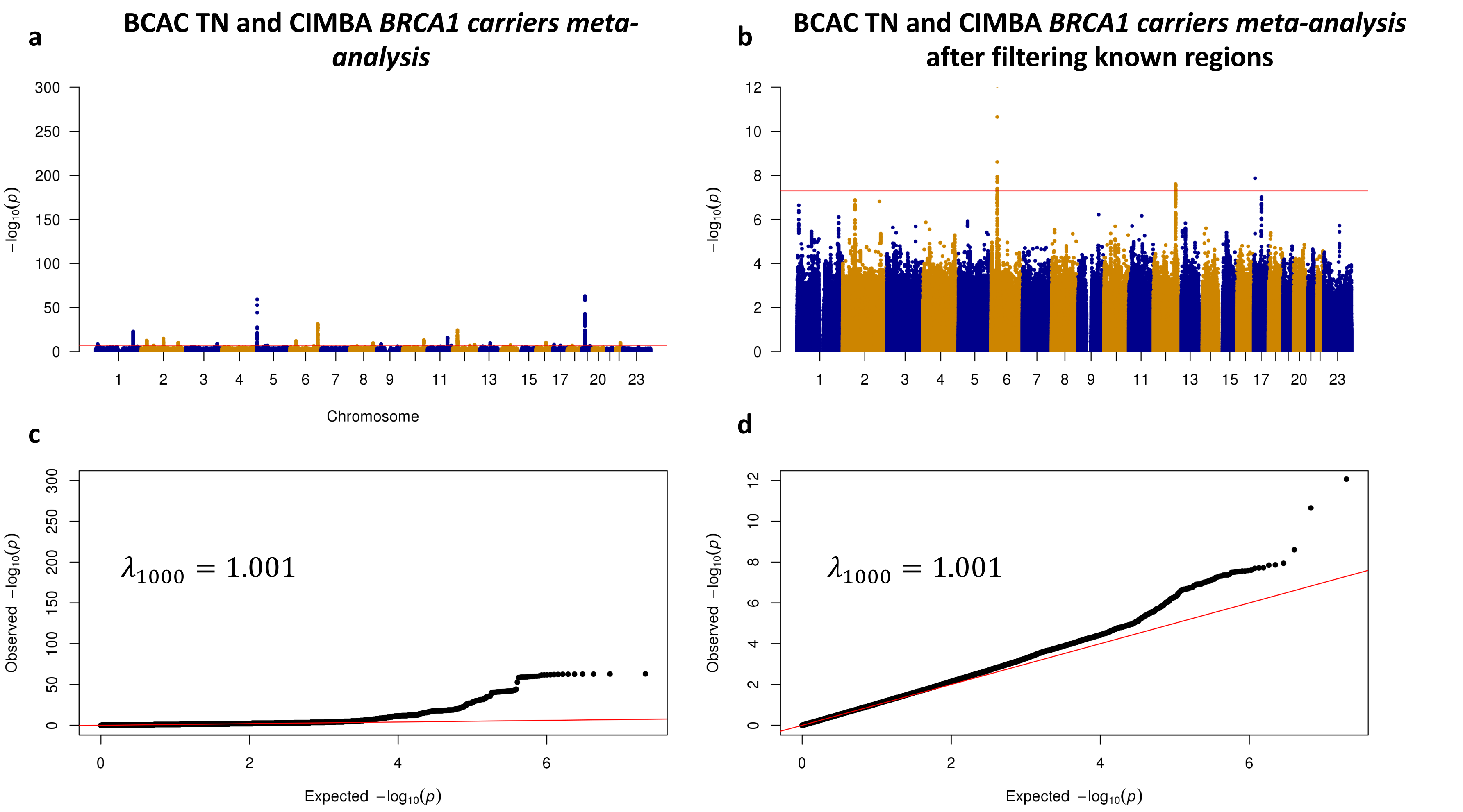


**1)** $\lambda_{1000}$ scale the genomic inflation factor $\lambda$ to a study with sample size of 1000 cases and 1000 controls using the formula $\lambda_{1000} = 1 + 500 * (\lambda - 1)/(\frac{1}{n_{cases}} + \frac{1}{n_{control}})$

**Supplementary figure 6.** Regional plots of the 32 identified breast cancer SNPs. The first 22 SNPs were identified through standard logistic regression, the following eight SNPs were identified through two-stage polytomous regression, the last two SNPs were identified through meta-analysis of BCAC TN and CIMBA *BRCA1* carriers. Plotted area is showing ±500 KB region around the identified susceptibility SNP.

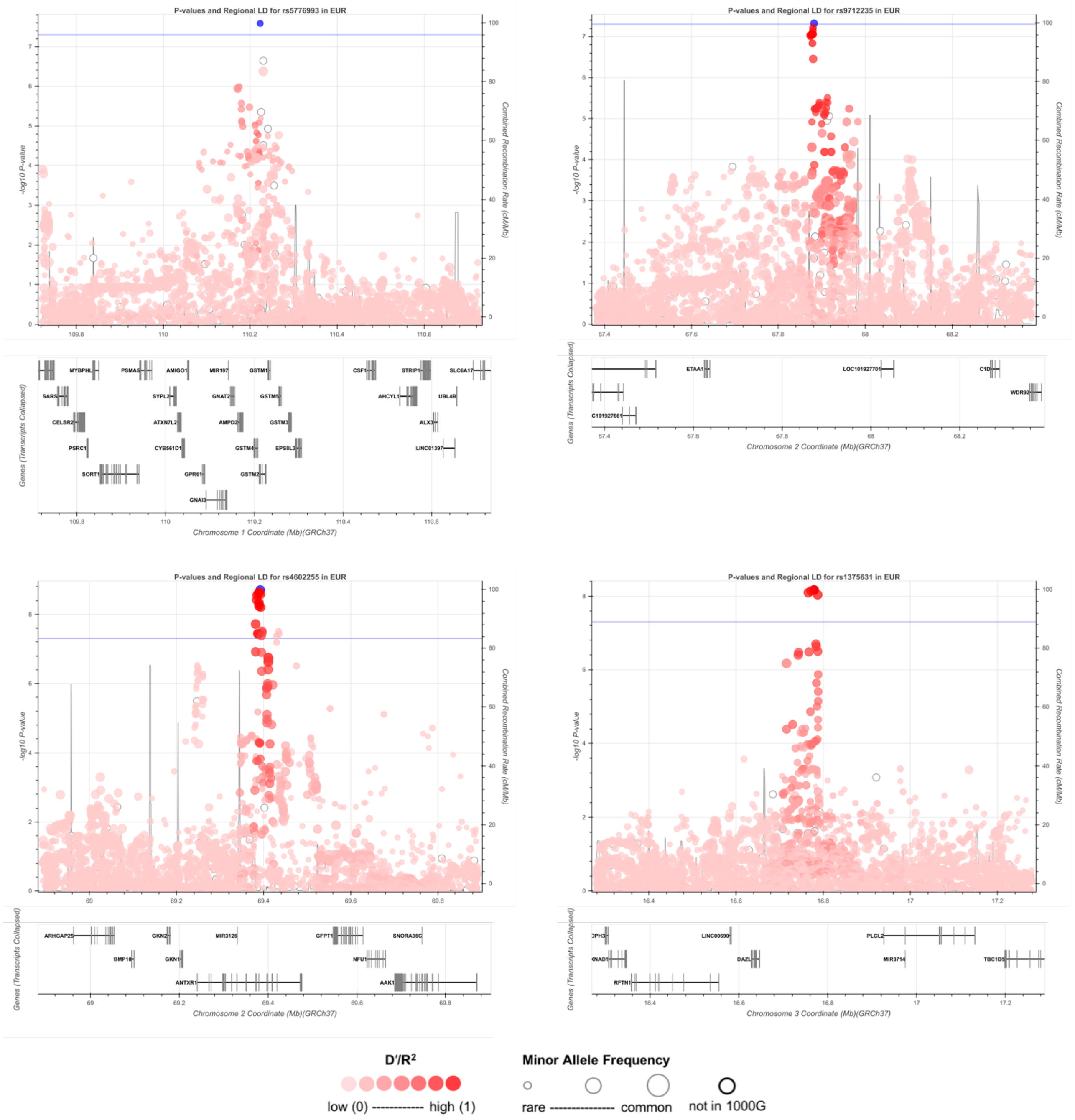

**Supplementary figure 6 continued.** Regional plots of the 32 identified breast cancer SNPs. The first 22 SNPs were identified through standard logistic regression, the following eight SNPs were identified through two-stage polytomous regression, the last two SNPs were identified through meta-analysis of BCAC TN and CIMBA *BRCA1* carriers. Plotted area is showing ±500 KB region around the identified susceptibility SNP.

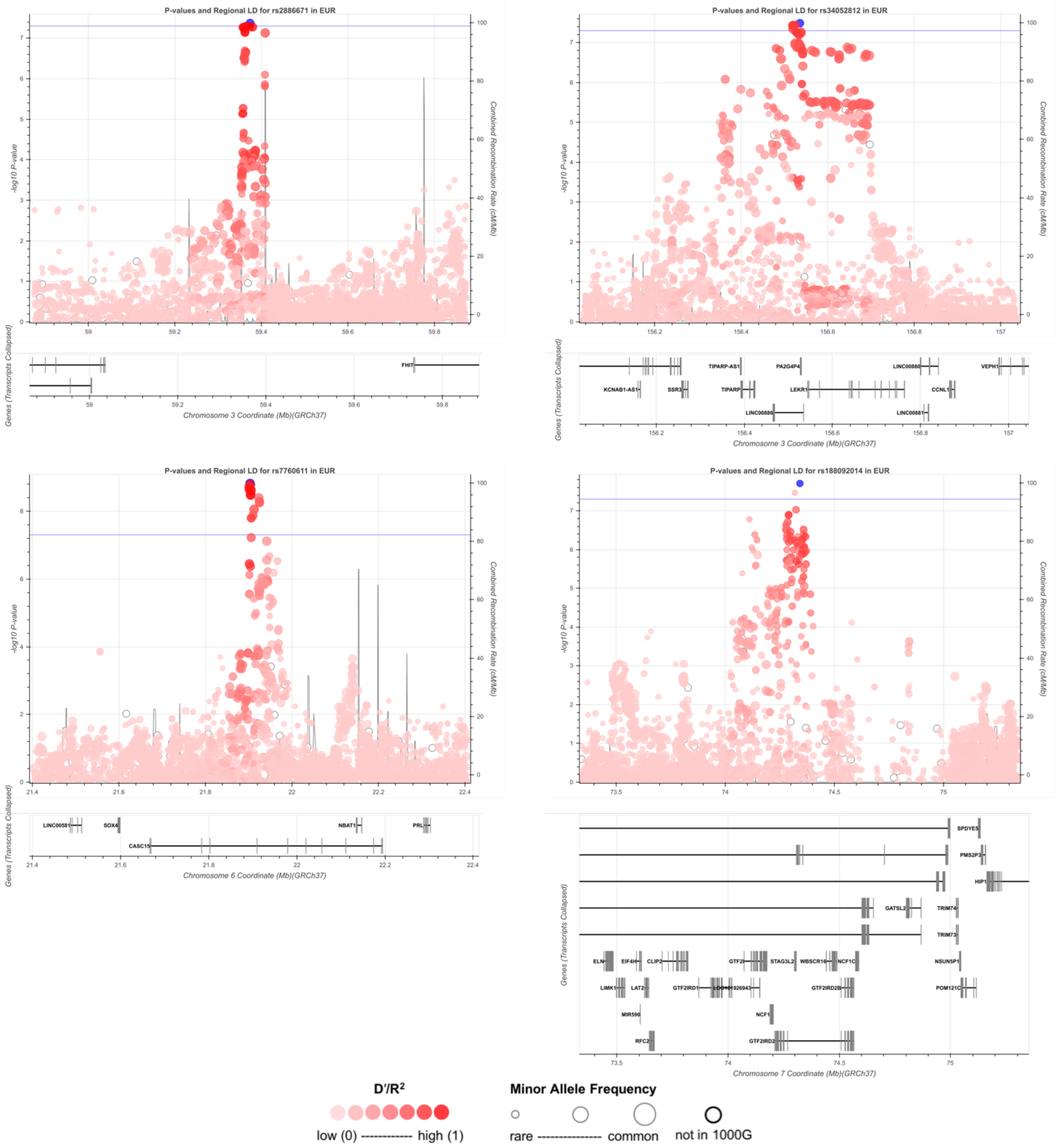

**Supplementary figure 6 continued.** Regional plots of the 32 identified breast cancer SNPs. The first 22 SNPs were identified through standard logistic regression, the following eight SNPs were identified through two-stage polytomous regression, the last two SNPs were identified through meta-analysis of BCAC TN and CIMBA *BRCA1* carriers. Plotted area is showing ±500 KB region around the identified susceptibility SNP.

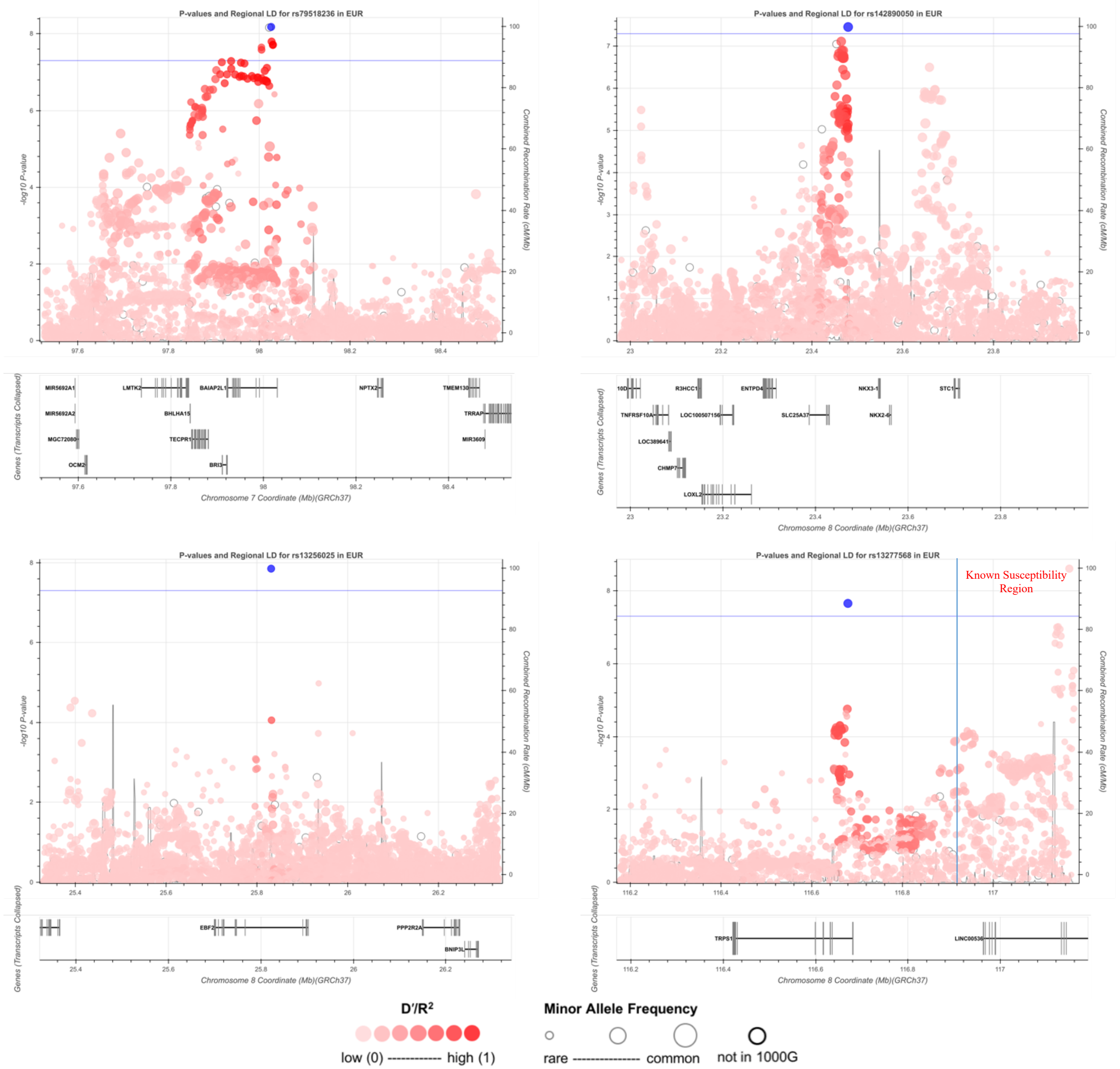

**Supplementary figure 6 continued.** Regional plots of the 32 identified breast cancer SNPs. The first 22 SNPs were identified through standard logistic regression, the following eight SNPs were identified through two-stage polytomous regression, the last two SNPs were identified through meta-analysis of BCAC TN and CIMBA *BRCA1* carriers. Plotted area is showing ±500 KB region around the identified susceptibility SNP.

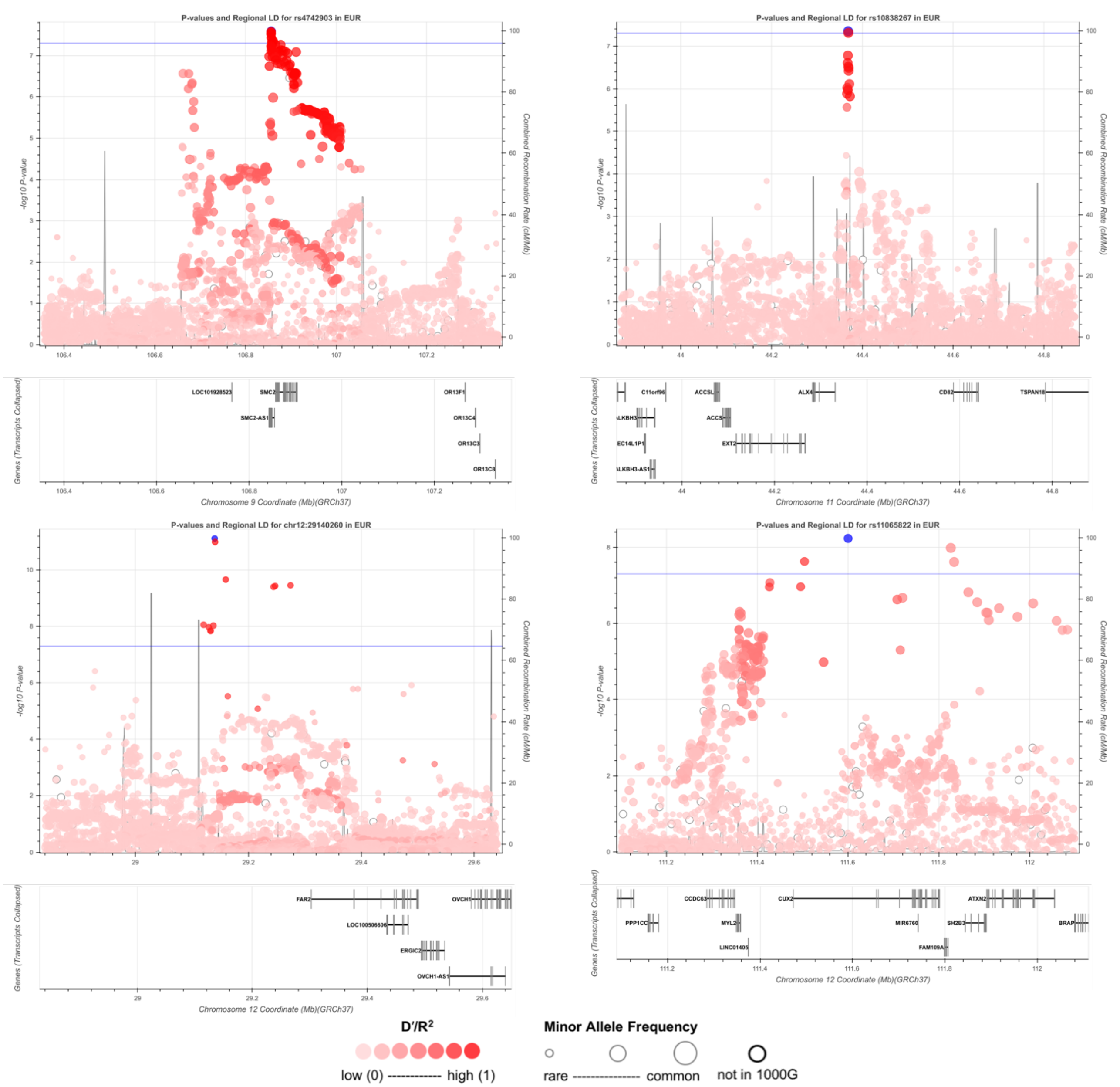

**Supplementary figure 6 continued.** Regional plots of the 32 identified breast cancer SNPs. The first 22 SNPs were identified through standard logistic regression, the following eight SNPs were identified through two-stage polytomous regression, the last two SNPs were identified through meta-analysis of BCAC TN and CIMBA *BRCA1* carriers. Plotted area is showing ±500 KB region around the identified susceptibility SNP.

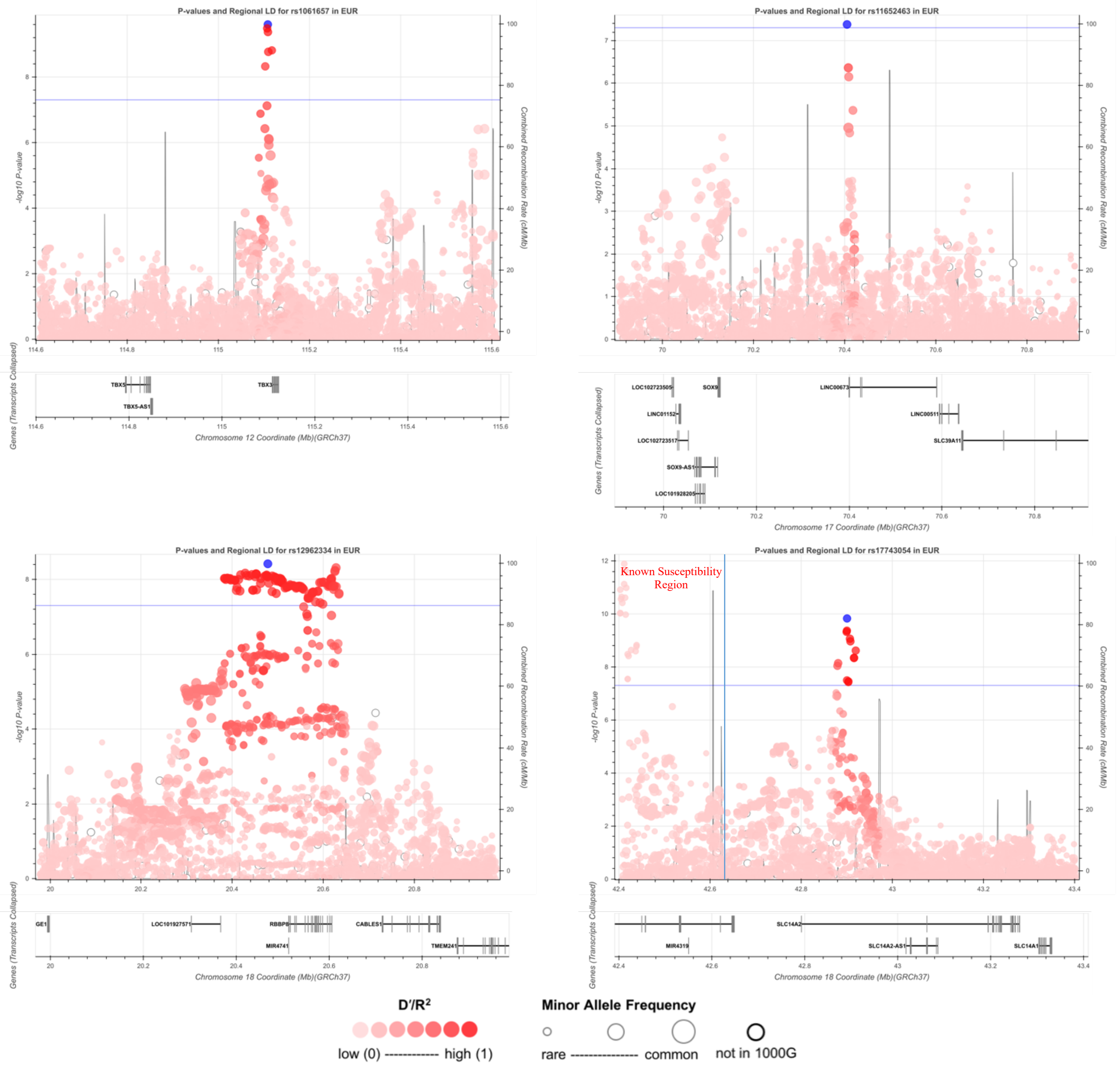

**Supplementary figure 6 continued.** Regional plots of the 32 identified breast cancer SNPs. The first 22 SNPs were identified through standard logistic regression, the following eight SNPs were identified through two-stage polytomous regression, the last two SNPs were identified through meta-analysis of BCAC TN and CIMBA *BRCA1* carriers. Plotted area is showing ±500 KB region around the identified susceptibility SNP.

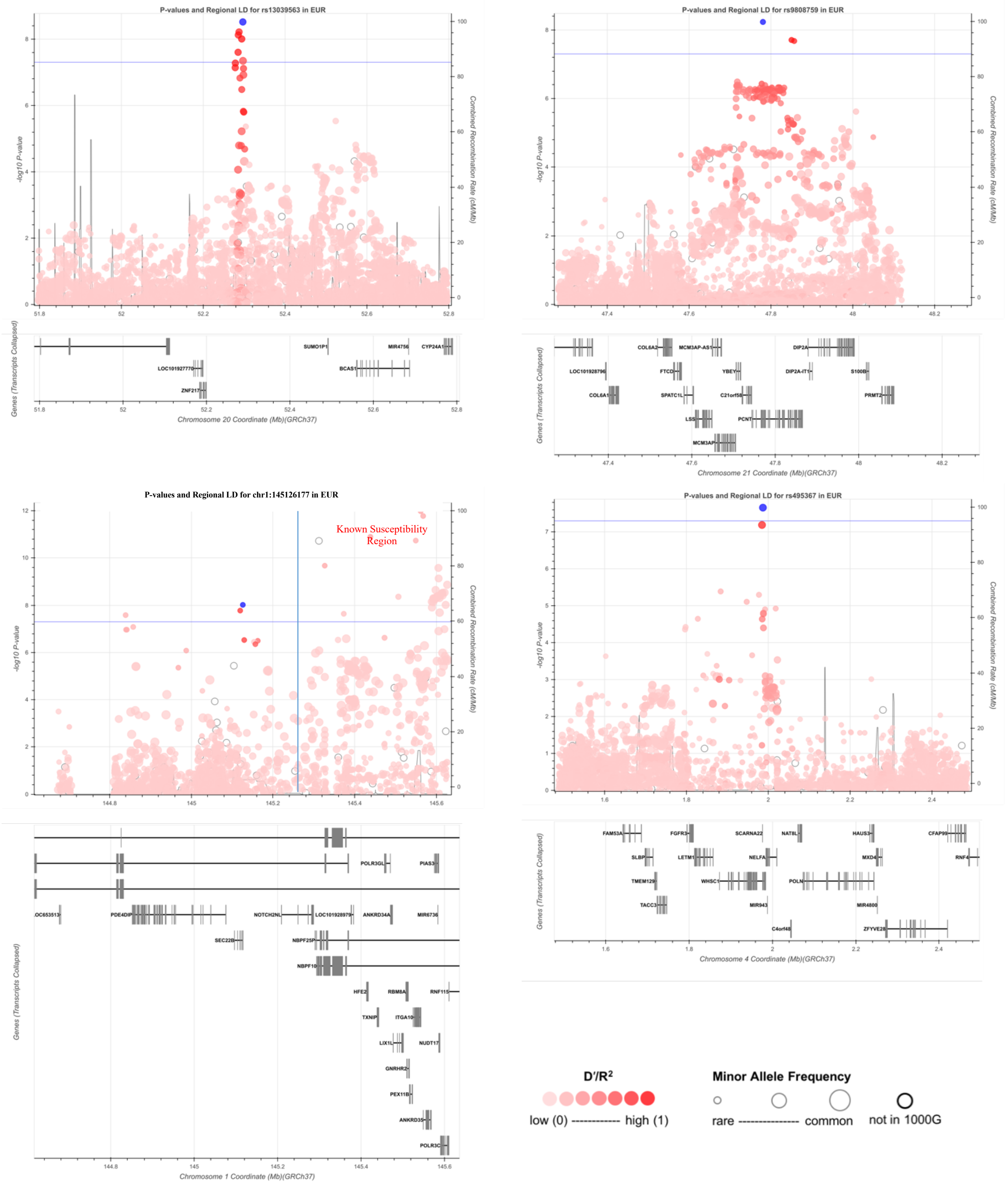

**Supplementary figure 6 continued.** Regional plots of the 32 identified breast cancer SNPs. The first 22 SNPs were identified through standard logistic regression, the following eight SNPs were identified through two-stage polytomous regression, the last two SNPs were identified through meta-analysis of BCAC TN and CIMBA *BRCA1* carriers. Plotted area is showing ±500 KB region around the identified susceptibility SNP.

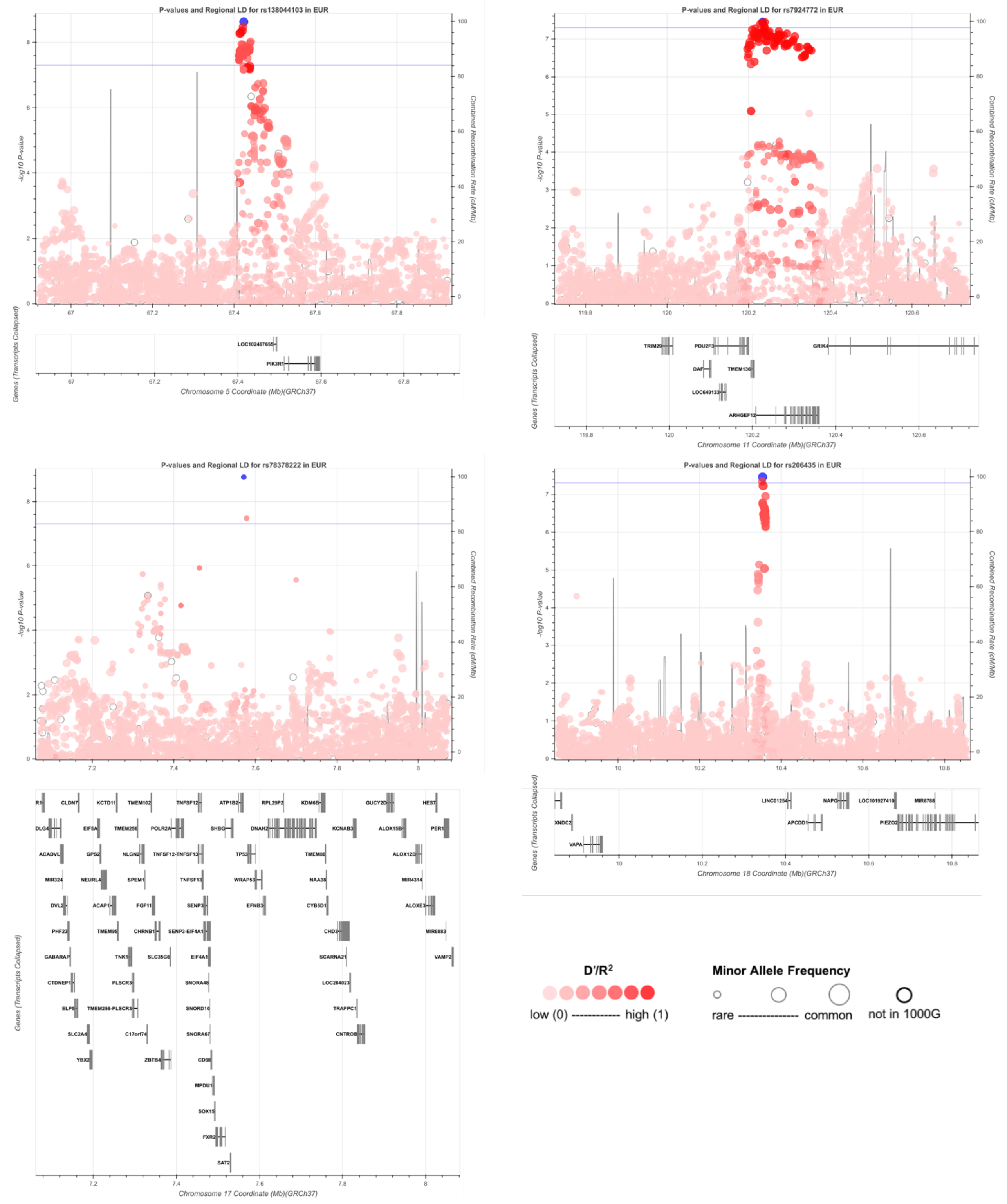

**Supplementary figure 6 continued.** Regional plots of the 32 identified breast cancer SNPs. The first 22 SNPs were identified through standard logistic regression, the following eight SNPs were identified through two-stage polytomous regression, the last two SNPs were identified through meta-analysis of BCAC TN and CIMBA *BRCA1* carriers. Plotted area is showing ±500 KB region around the identified susceptibility SNP.

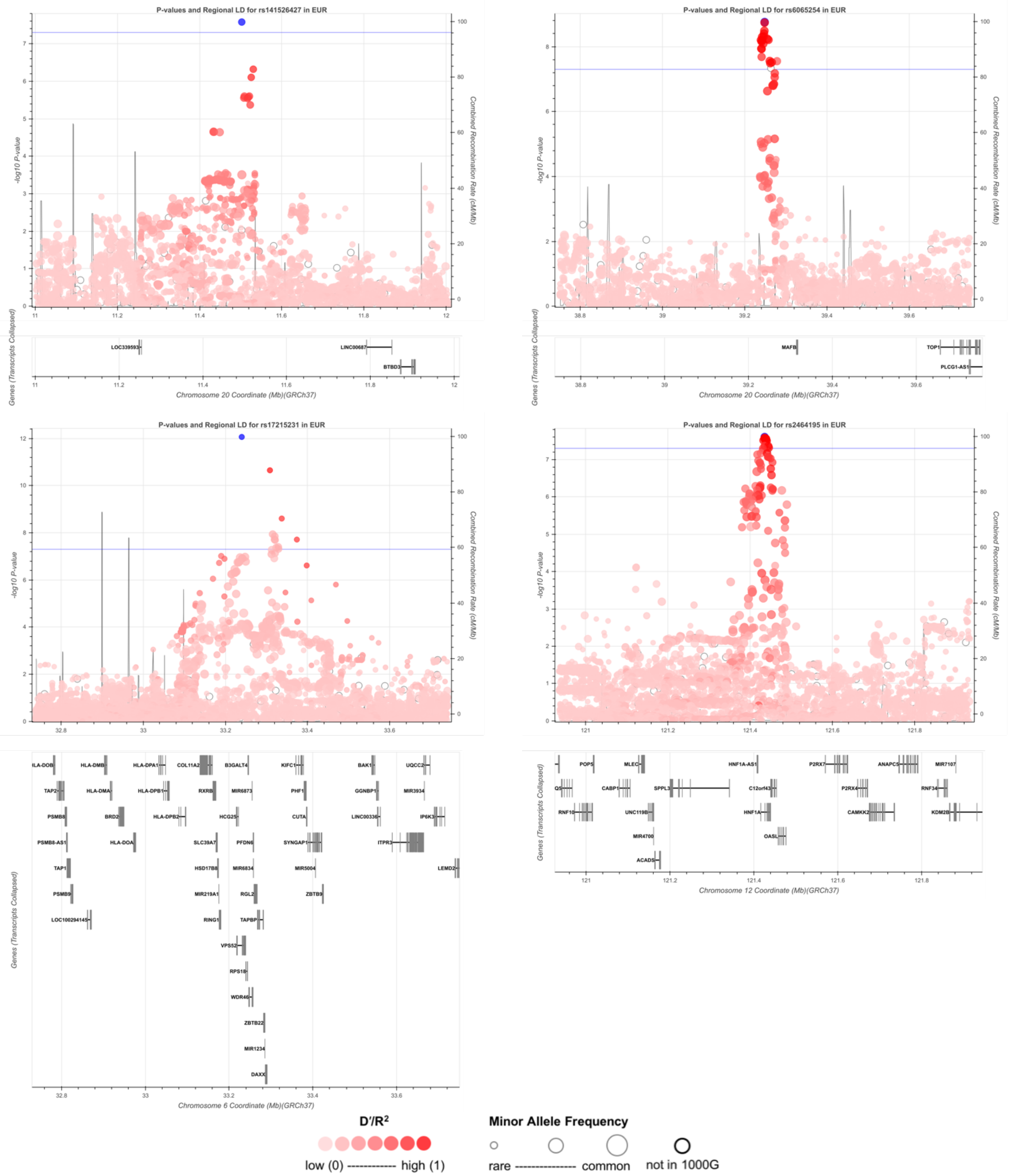

**Supplementary figure 7**. Country Specific sensitivity analysis of 8 genome-wide significant loci identified using the two-stage regression models. We also show chr22:40042814 though this locus was dropped since the signal was observed only in studies from the USA.

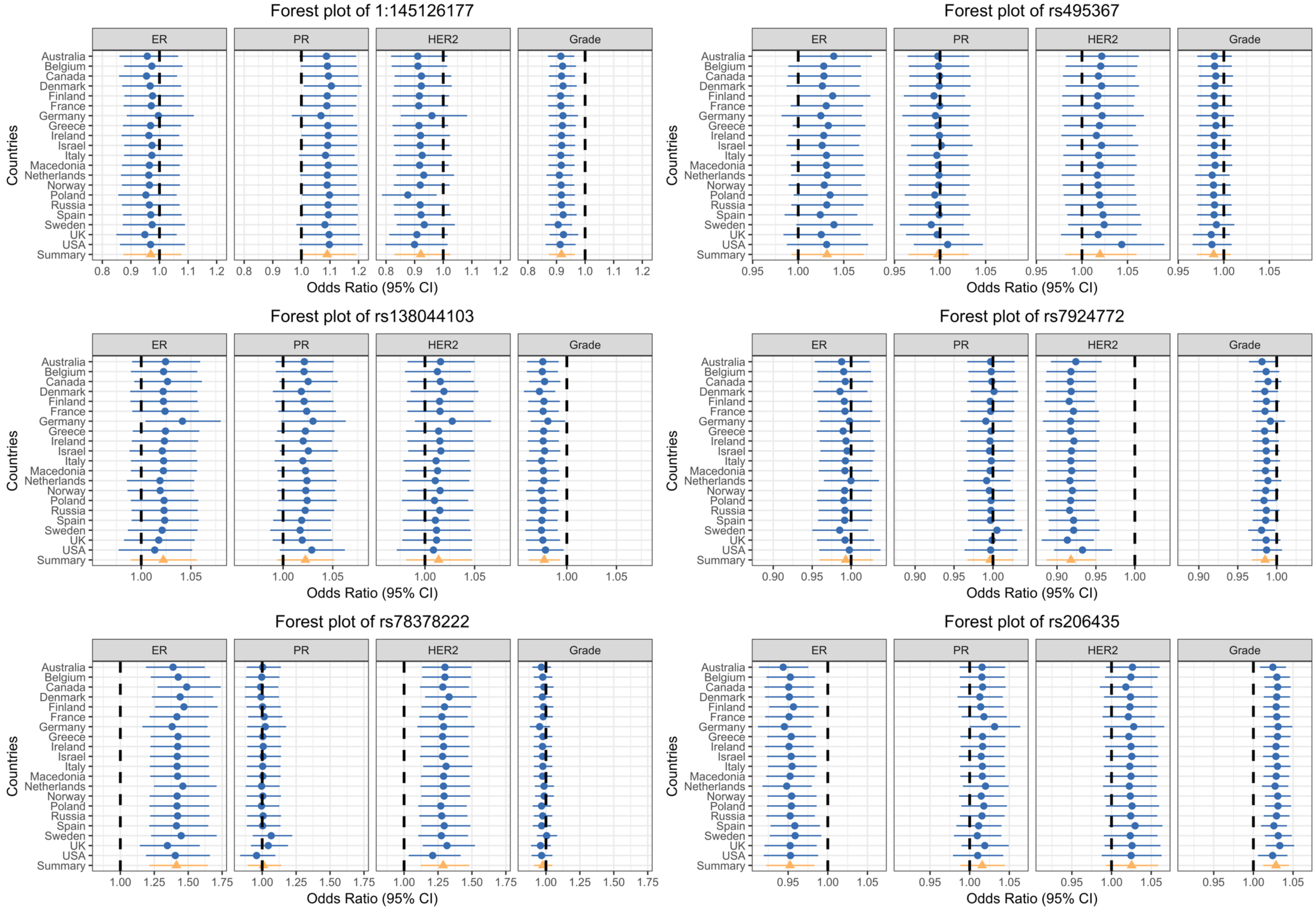

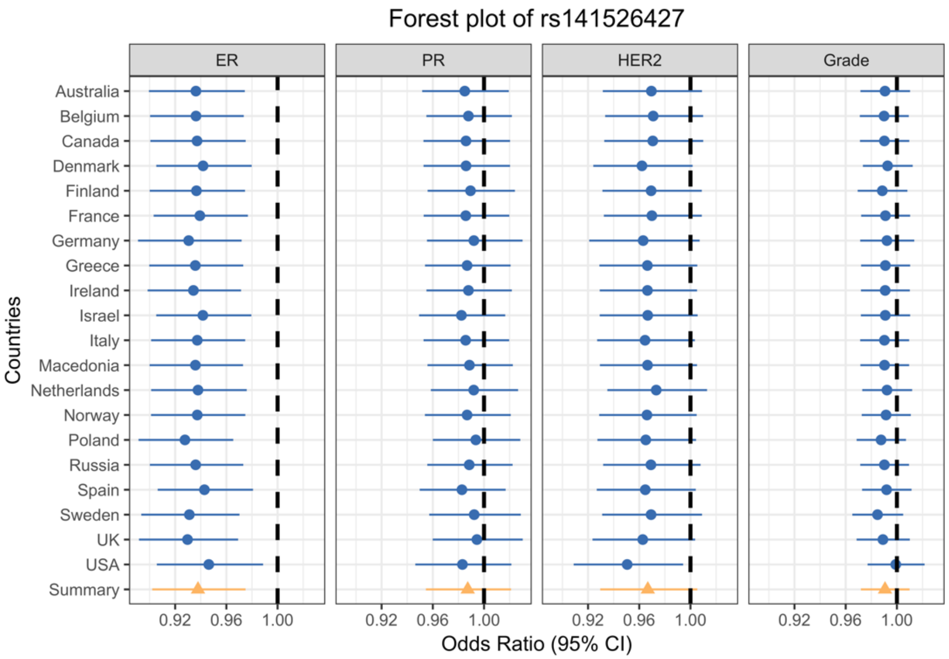
Forest plot of rs141526427
ER
PR
HER2
Grade
Countries
Australia
Belgium
Canada
Denmark
Finland
France
Germany
Greece
Ireland
Israel
Italy
Macedonia
Netherlands
Norway
Poland
Russia
Spain
Sweden
UK
USA
Summary
0.92
0.96
1.00
Odds Ratio (95% CI)

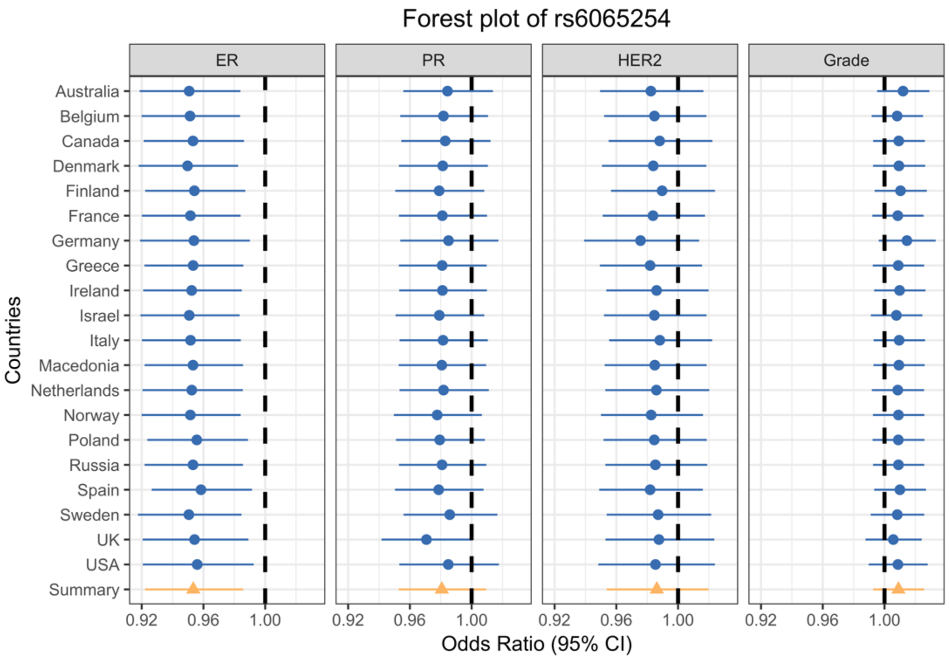
Forest plot of rs6065254
ER
PR
HER2
Grade
Countries
Australia
Belgium
Canada
Denmark
Finland
France
Germany
Greece
Ireland
Israel
Italy
Macedonia
Netherlands
Norway
Poland
Russia
Spain
Sweden
UK
USA
Summary
0.92
0.96
1.00
Odds Ratio (95% CI)

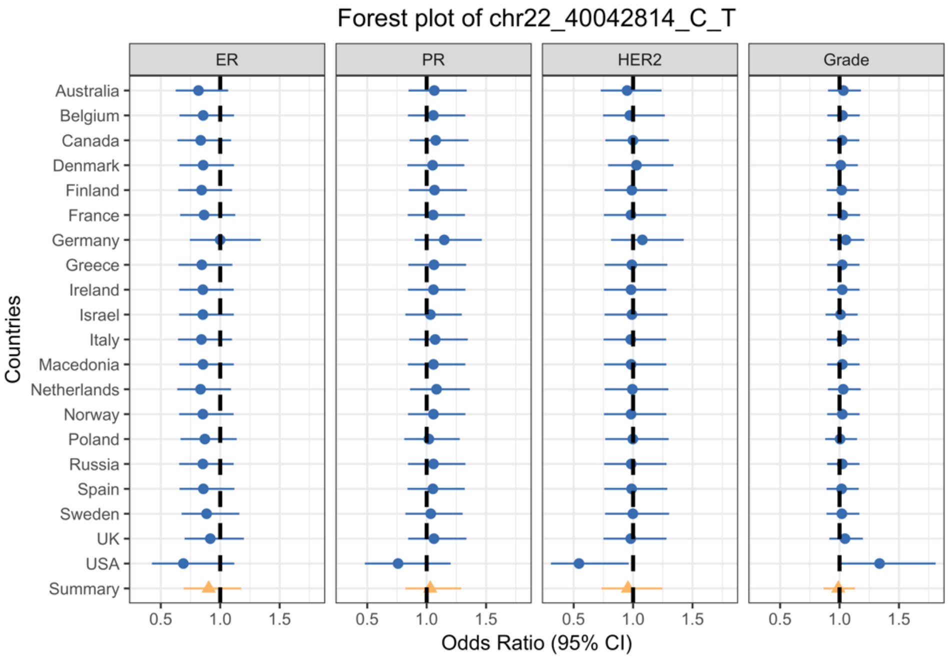
Forest plot of chr22_40042814_C_T
ER
PR
HER2
Grade
Countries
Australia
Belgium
Canada
Denmark
Finland
France
Germany
Greece
Ireland
Israel
Italy
Macedonia
Netherlands
Norway
Poland
Russia
Spain
Sweden
UK
USA
Summary
0.5
1.0
1.5
Odds Ratio (95% CI)

**Supplementary Figure 8**. Risk[1] (right panel) and p-values (left panel) of SNPs for breast cancer subtypes defined by intrinsic-like subtypes[2] among loci identified using standard logistic regression

| SNP | chromosome | Position | MAF | Global etiologic heterogeneity $P^3$ | Tumor characteristic heterogeneity $P^4$ ER[5] | PR[5] | HER2[5] | grade |
|---|---|---|---|---|---|---|---|---|
| rs5776993 | 1 | 110,222,901 | 0.12 | 0.50 | 0.29 | 0.54 | 0.35 | 0.29 |
| rs9712235 | 2 | 67,881,757 | 0.26 | 6.70E-03 | 3.68E-04 | 0.02 | 0.03 | 0.85 |
| rs4602255 | 2 | 69,392,128 | 0.45 | 0.17 | 0.69 | 0.67 | 0.62 | 0.11 |
| rs1375631 | 3 | 16,778,867 | 0.5 | 0.40 | 0.09 | 0.21 | 0.57 | 0.55 |
| rs2886671 | 3 | 59,373,745 | 0.43 | 0.67 | 0.75 | 0.52 | 0.43 | 0.39 |
| rs34052812 | 3 | 156,535,958 | 0.33 | 0.98 | 0.74 | 0.72 | 0.62 | 0.97 |
| rs7760611 | 6 | 21,903,533 | 0.47 | 1.40E-03 | 0.03 | 0.28 | 0.17 | 0.87 |
| rs188092014 | 7 | 74,341,926 | 0.19 | 0.19 | 0.05 | 0.30 | 0.78 | 0.19 |
| rs79518236 | 7 | 98,026,554 | 0.23 | 1.57E-03 | 0.37 | 0.02 | 0.30 | 1.94E-03 |
| rs142890050 | 8 | 23,480,253 | 0.46 | 0.48 | 0.60 | 0.69 | 0.91 | 0.26 |
| rs13256025 | 8 | 25,831,778 | 0.2 | 0.69 | 0.32 | 0.59 | 0.81 | 0.80 |

Breast cancer risk by subtypes

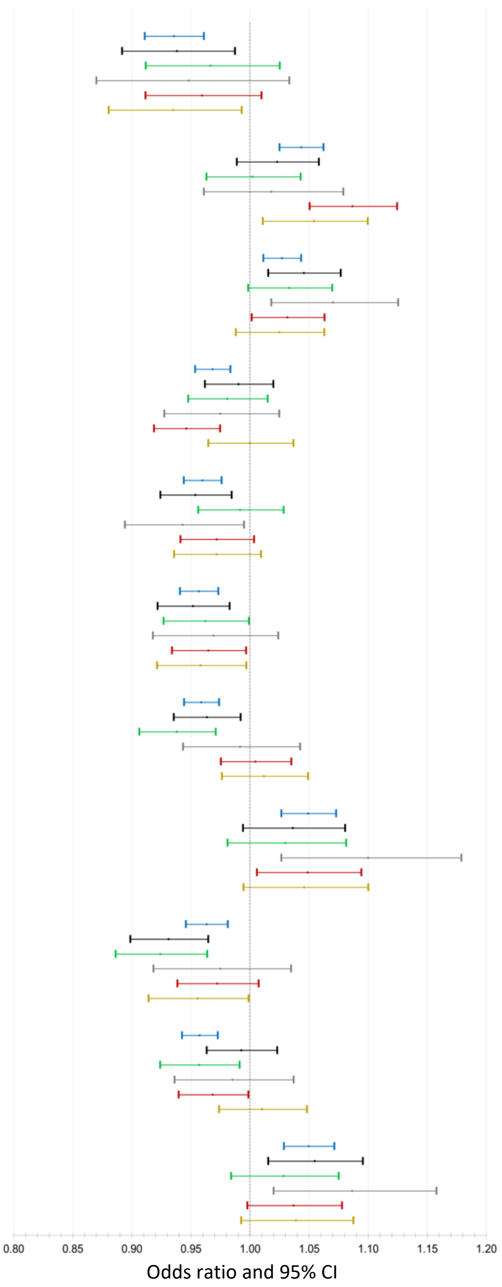


Luminal A-like — Luminal B/HER2-negative-like — Luminal B-like — HER2-enriched-like — Triple-negative — *BRCA1* mutation carriers

1 Per-minor allele odds ratio (95% confidence limits)

2. Luminal A-like (ER+ and/or PR+, HER2-, grade 1 & 2); luminal B/HER2-negative-like (ER+ and/or PR+, HER2-, grade 3); luminal B-like (ER+ and/or PR+, HER2+); HER2-enriched-like (ER- and PR-, HER2+); triple-negative (ER-, PR-, HER2-)

3. Based on a mixed-effect two-stage polytomous model testing for heterogeneity between susceptibility SNPs and ER, PR, HER2, and grade, where ER was entered into the model as a fixed-effect term and PR, HER2, and grade were entered into the model as random-effect terms.

4. Results from second stage case-case parameters from a fixed effect two-stage polytomous model testing for heterogeneity between susceptibility SNPs and ER, PR, HER2, and grade, where ER, PR, HER2, and grade are mutually adjusted for each other

5. Estrogen receptor (ER), progesterone receptor (PR) and human epidermal growth factor receptor 2 (HER2)

**Supplementary Figure 8 continued**. Risk[1] of breast cancer subtypes defined by intrinsic-like subtypes[2] among loci identified using standard logistic regression

| SNP | chromosome | Position | MAF | Global etiologic heterogeneity $P^3$ | Tumor characteristic heterogeneity $P^4$ | | | |
|---|---|---|---|---|---|---|---|---|
| | | | | | ER | PR | HER2 | grade |
| rs13277568 | 8 | 116,679,547 | 0.37 | 0.06 | 0.90 | 0.19 | 0.17 | 0.34 |
| rs4742903 | 9 | 106,856,793 | 0.44 | 2.80E-03 | 0.34 | 0.37 | 0.05 | 0.01 |
| rs10838267 | 11 | 44,368,892 | 0.45 | 0.85 | 0.39 | 0.54 | 0.69 | 0.81 |
| 12:29140260 | 12 | 29,140,260 | 0.09 | 0.39 | 0.31 | 0.63 | 0.67 | 0.37 |
| rs11065822 | 12 | 111,600,134 | 0.37 | 0.87 | 0.52 | 0.70 | 0.69 | 0.78 |
| rs1061657 | 12 | 115,108,136 | 0.26 | 0.04 | 0.07 | 0.45 | 0.35 | 0.38 |
| rs11652463 | 17 | 70,405,095 | 0.31 | 0.21 | 0.63 | 0.17 | 0.90 | 0.92 |
| rs12962334 | 18 | 20,477,934 | 0.32 | 0.06 | 0.02 | 0.28 | 0.70 | 0.15 |
| rs17743054 | 18 | 42,900,892 | 0.28 | 2.40E-06 | 0.04 | 0.25 | 0.01 | 0.01 |
| rs13039563 | 20 | 52,296,849 | 0.24 | 4.88E-03 | 0.14 | 0.26 | 0.81 | 0.44 |
| rs9808759 | 21 | 47,780,223 | 0.07 | 0.71 | 0.50 | 0.29 | 0.44 | 0.29 |

Breast cancer risk by subtypes

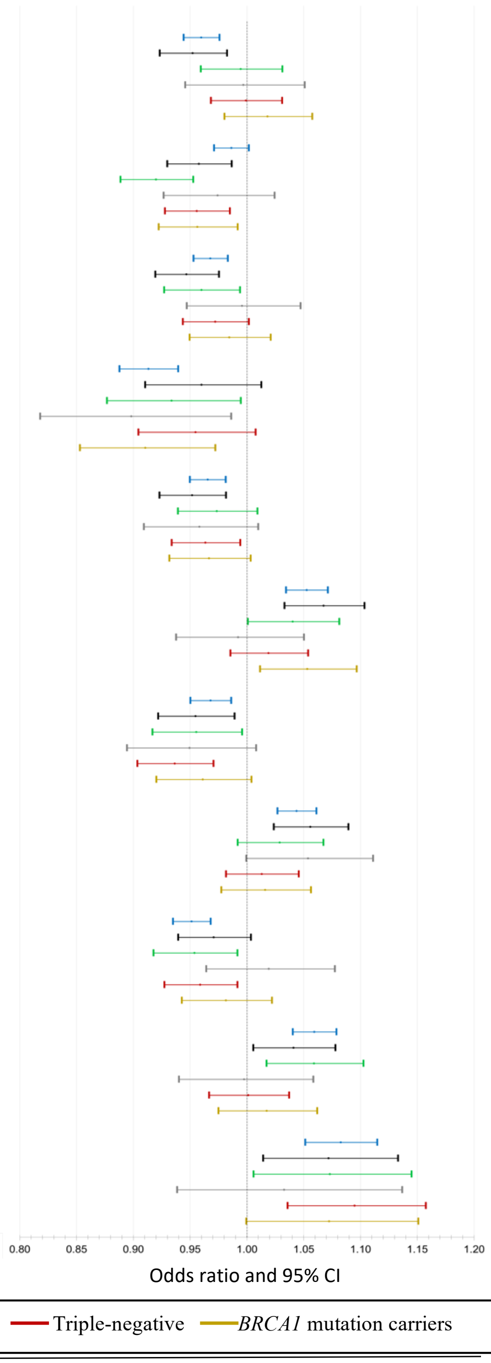


1 Per-minor allele odds ratio (95% confidence limits)

2. Luminal A-like (ER+ and/or PR+, HER2-, grade 1 & 2); luminal B/HER2-negative-like (ER+ and/or PR+, HER2-, grade 3); luminal B-like (ER+ and/or PR+, HER2+); HER2-enriched-like (ER- and PR-, HER2+); triple-negative (ER-, PR-, HER2-)

3. Based on a mixed-effect two-stage polytomous model testing for heterogeneity between susceptibility SNPs and ER, PR, HER2, and grade, where ER was entered into the model as a fixed-effect term and PR, HER2, and grade were entered into the model as random-effect terms.

4. Results from second stage case-case parameters from a fixed effect two-stage polytomous model testing for heterogeneity between susceptibility SNPs and ER, PR, HER2, and grade, where ER, PR, HER2, and grade are mutually adjusted for each other

5. Estrogen receptor (ER), progesterone receptor (PR) and human epidermal growth factor receptor 2 (HER2)

**Supplementary Figure 9** Risk[1] of breast cancer subtypes defined by intrinsic-like subtypes[2] among loci identified using the two-stage polytomous logistic regression model and the CIMBA / BCAC triple-negative meta-analysis

| SNP | chromosome | Position | MAF | Global etiologic heterogeneity $P^3$ | Tumor characteristic heterogeneity $P^4$ | | | |
|---|---|---|---|---|---|---|---|---|
| | | | | | ER[5] | PR[5] | HER2[5] | grade |
| chr1:145126177 | 1 | 145,126,177 | 0.04 | 2.8E-06 | 5.04E-01 | 5.35E-02 | 1.18E-01 | 6.87E-04 |
| rs495367 | 4 | 1,986,972 | 0.35 | 5.8E-02 | 1.35E-01 | 9.22E-01 | 3.09E-01 | 2.42E-01 |
| chr5:67424121 | 5 | 67,424,121 | 0.45 | 5.2E-07 | 1.70E-01 | 1.31E-01 | 4.20E-01 | 2.79E-03 |
| rs7924772 | 11 | 120,233,626 | 0.39 | 1.4E-03 | 6.69E-01 | 8.31E-01 | 1.41E-06 | 9.95E-02 |
| rs78378222 | 17 | 7,571,752 | 0.01 | 9.1E-08 | 7.01E-06 | 8.96E-01 | 2.67E-04 | 5.15E-01 |
| rs206435 | 18 | 10,354,649 | 0.5 | 1.1E-09 | 2.79E-03 | 2.51E-01 | 1.44E-01 | 2.83E-04 |
| rs141526427 | 20 | 11,502,618 | 0.25 | 6.2E-05 | 1.26E-03 | 4.44E-01 | 8.88E-02 | 3.22E-01 |
| rs6065254 | 20 | 39,248,265 | 0.39 | 7.3E-07 | 4.34E-03 | 1.98E-01 | 3.92E-01 | 2.74E-01 |
| rs17215231 | 6 | 33,239,869 | 0.08 | 2.4E-06 | 4.40E-02 | 2.46E-01 | 9.12E-03 | 7.87E-03 |
| rs2464195 | 12 | 121,435,475 | 0.37 | 1.0E-02 | 8.92E-02 | 9.99E-01 | 4.05E-01 | 3.41E-01 |

Breast cancer risk by subtypes

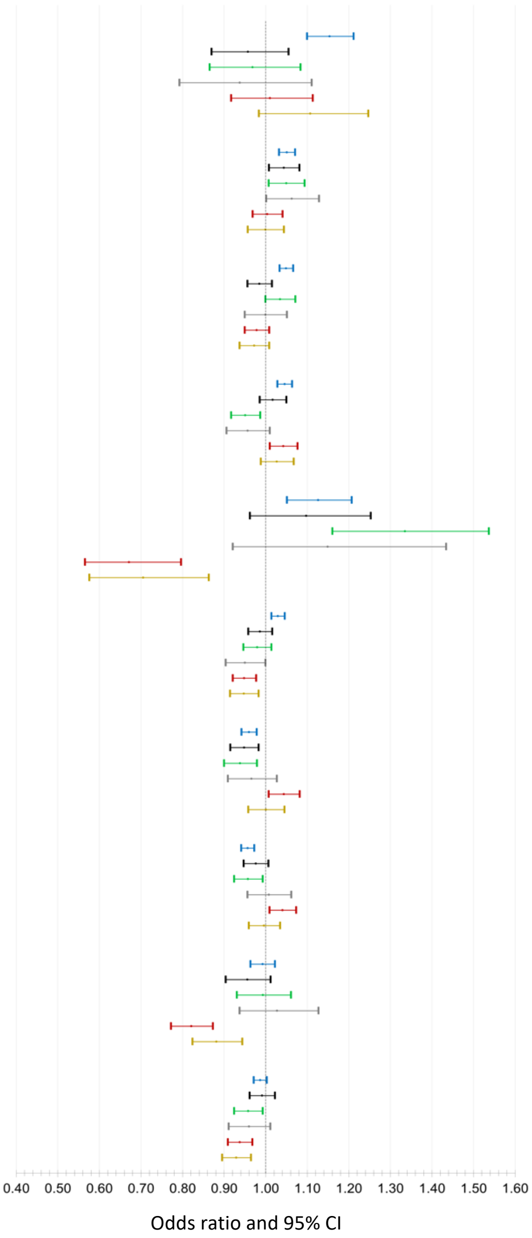


Luminal A-like — Luminal B/HER2-negative-like — Luminal B-like — HER2-enriched-like — Triple-negative — *BRCA1* mutation carriers

1 Per-minor allele odds ratio (95% confidence limits)

2. Luminal A-like (ER+ and/or PR+, HER2-, grade 1 & 2); luminal B/HER2-negative-like (ER+ and/or PR+, HER2-, grade 3); luminal B-like (ER+ and/or PR+, HER2+); HER2-enriched-like (ER- and PR-, HER2+); triple-negative (ER-, PR-, HER2-)

3. Based on a mixed-effect two-stage polytomous model testing for heterogeneity between susceptibility SNPs and ER, PR, HER2, and grade, where ER was entered into the model as a fixed-effect term and PR, HER2, and grade were entered into the model as random-effect terms.

4. Results from second stage case-case parameters from a fixed effect two-stage polytomous model testing for heterogeneity between susceptibility SNPs and ER, PR, HER2, and grade, where ER, PR, HER2, and grade are mutually adjusted for each other

5. Estrogen receptor (ER), progesterone receptor (PR) and human epidermal growth factor receptor 2 (HER2)

**Supplementary figure 10**. **a)** Enrichment analysis[1] results for 24 non-cell-type-specific, publicly available annotations for luminal A-like subtypes and TN subtypes. **b)** Enrichment analysis[1] results for 24 main annotations with ±500 bp extension for luminal A-like subtypes and TN subtypes. No significant differences were found between luminal A-like and TN after adjusting for multiple testing.

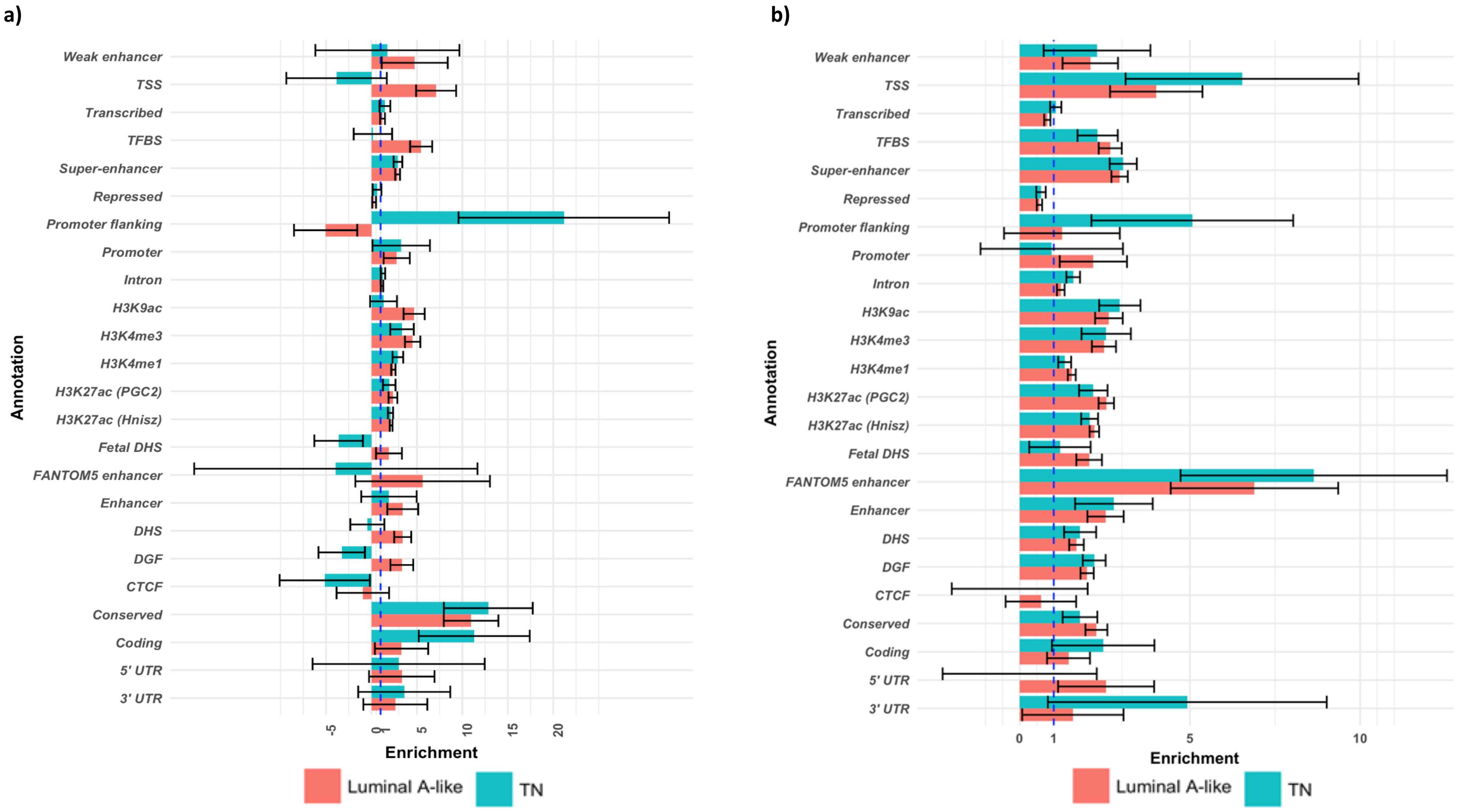


[1] Error bars represent Jackknife standard errors around the estimates of enrichment.

**Supplementary figure 11**. Enrichment analysis results for 220 cell-type-specific annotations of four histone marks - H3K4me1, H3K4me3, H3K9ac and H3K27ac – in the luminal A-like and TN subtypes. Both luminal A-like and TN subtypes were enriched for gastrointestinal cell types and suppression of central nervous system cells.

**a)** Heatmap showing patterns of cell-type specific enrichment for histone marks H3K27ac in luminal A-like tumors and TN tumors

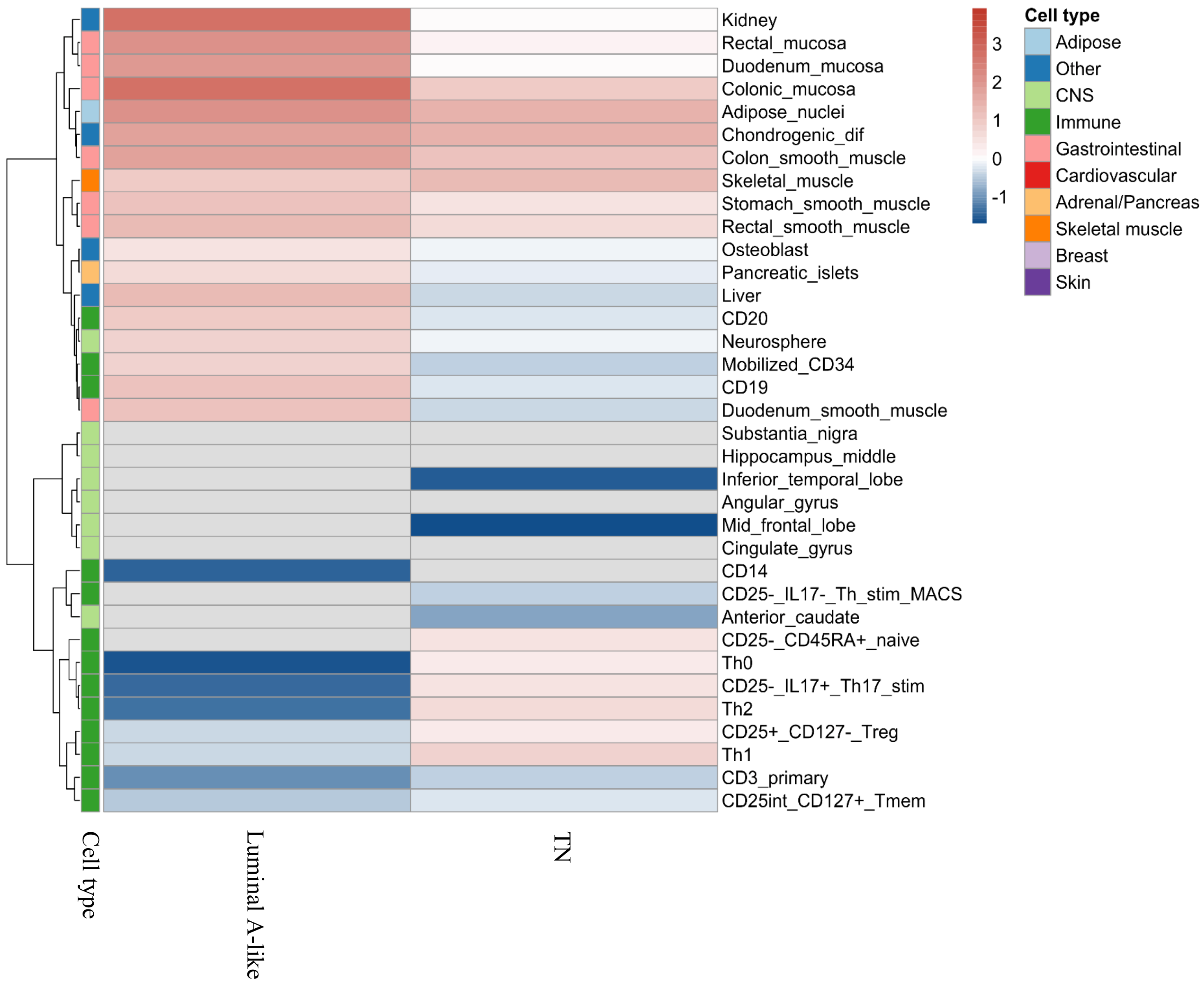

**b)** Heatmap showing patterns of cell-type specific enrichment for histone marks H3K4me1 in luminal A-like tumors and TN tumors

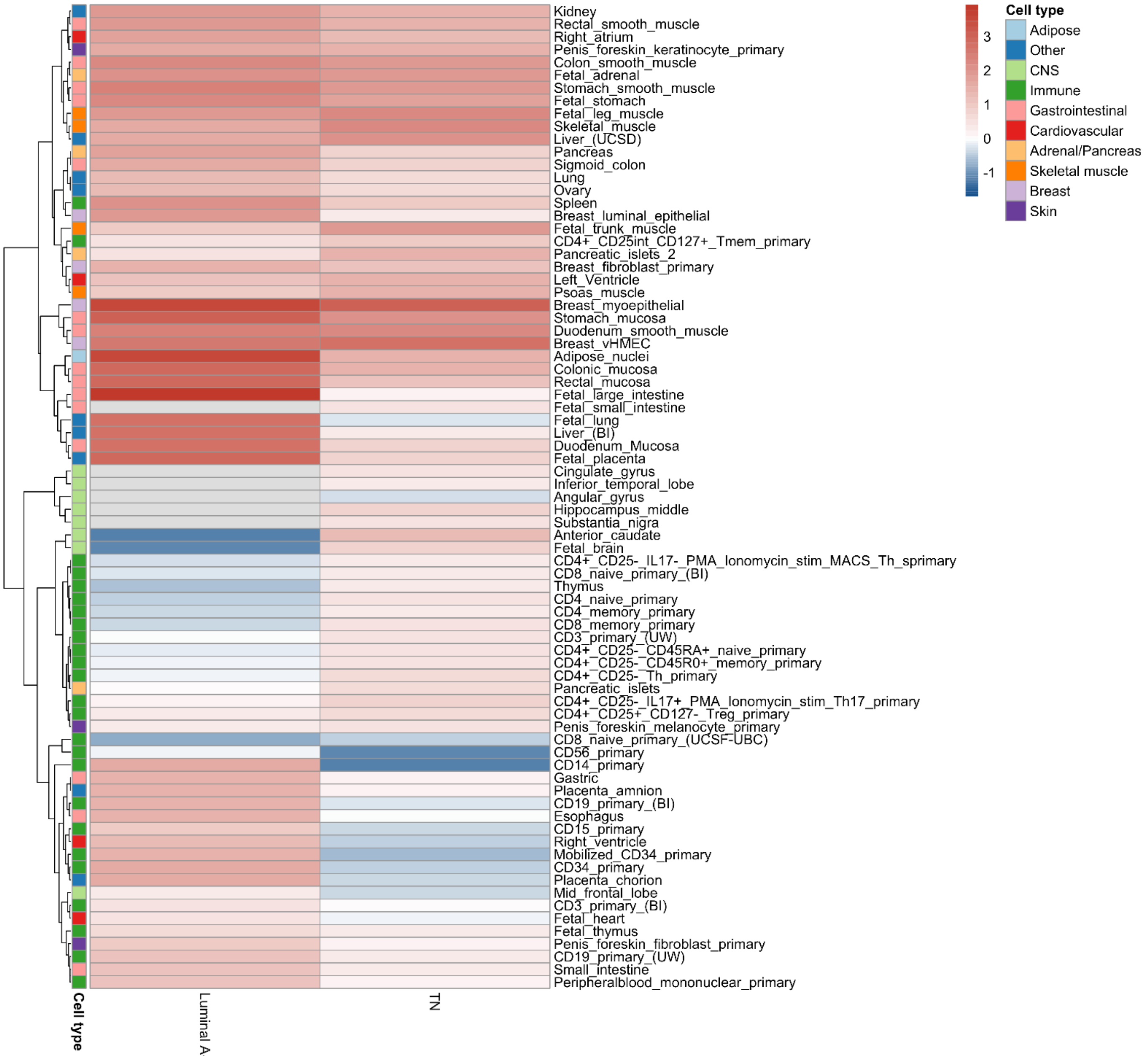

**c)** Heatmap showing patterns of cell-type specific enrichment for histone marks H3K4me3 in luminal A-like tumors and TN tumors

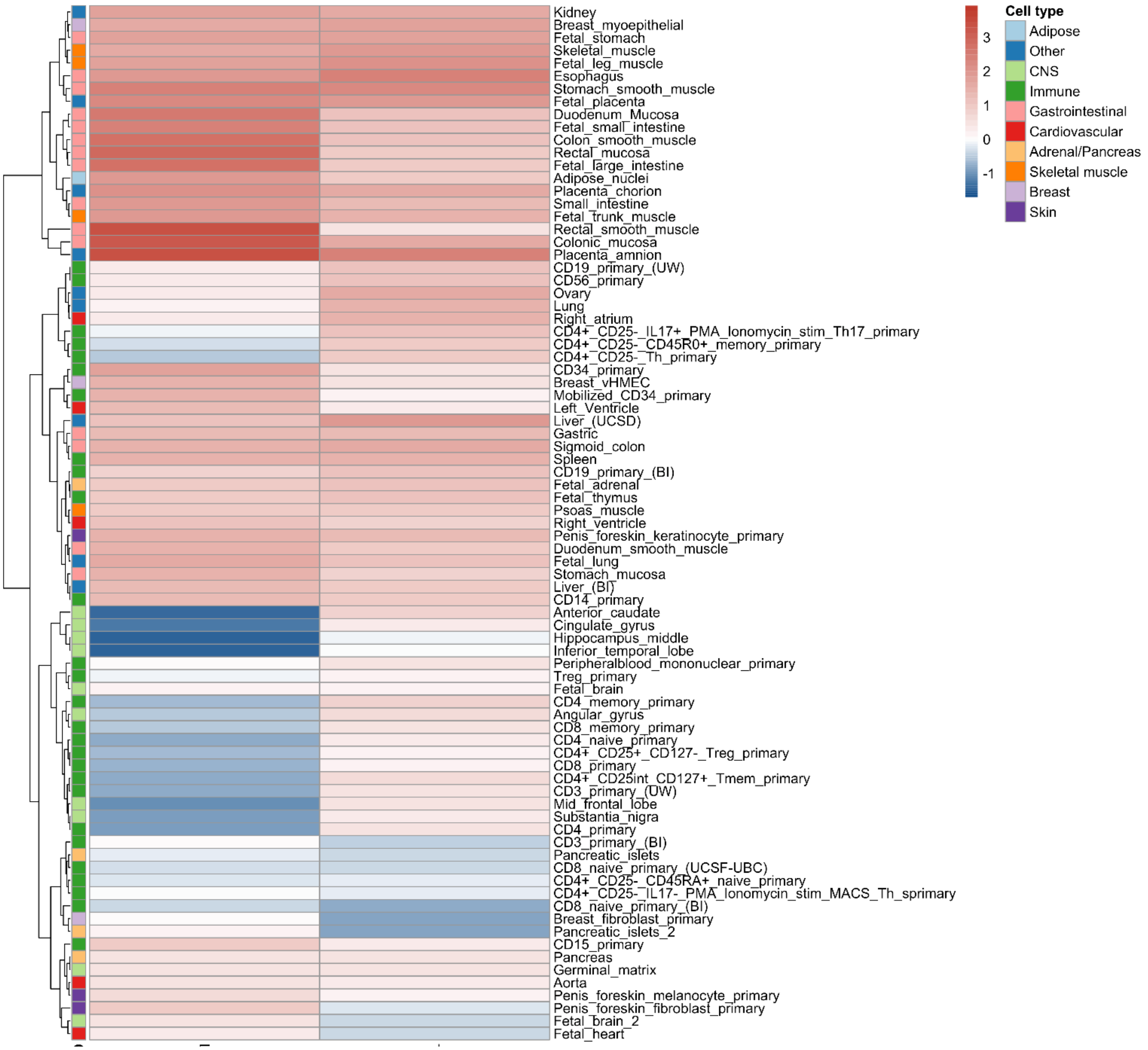

**d)** Heatmap showing patterns of cell-type specific enrichment for histone marks H3K9ac in luminal A-like tumors and TN tumors

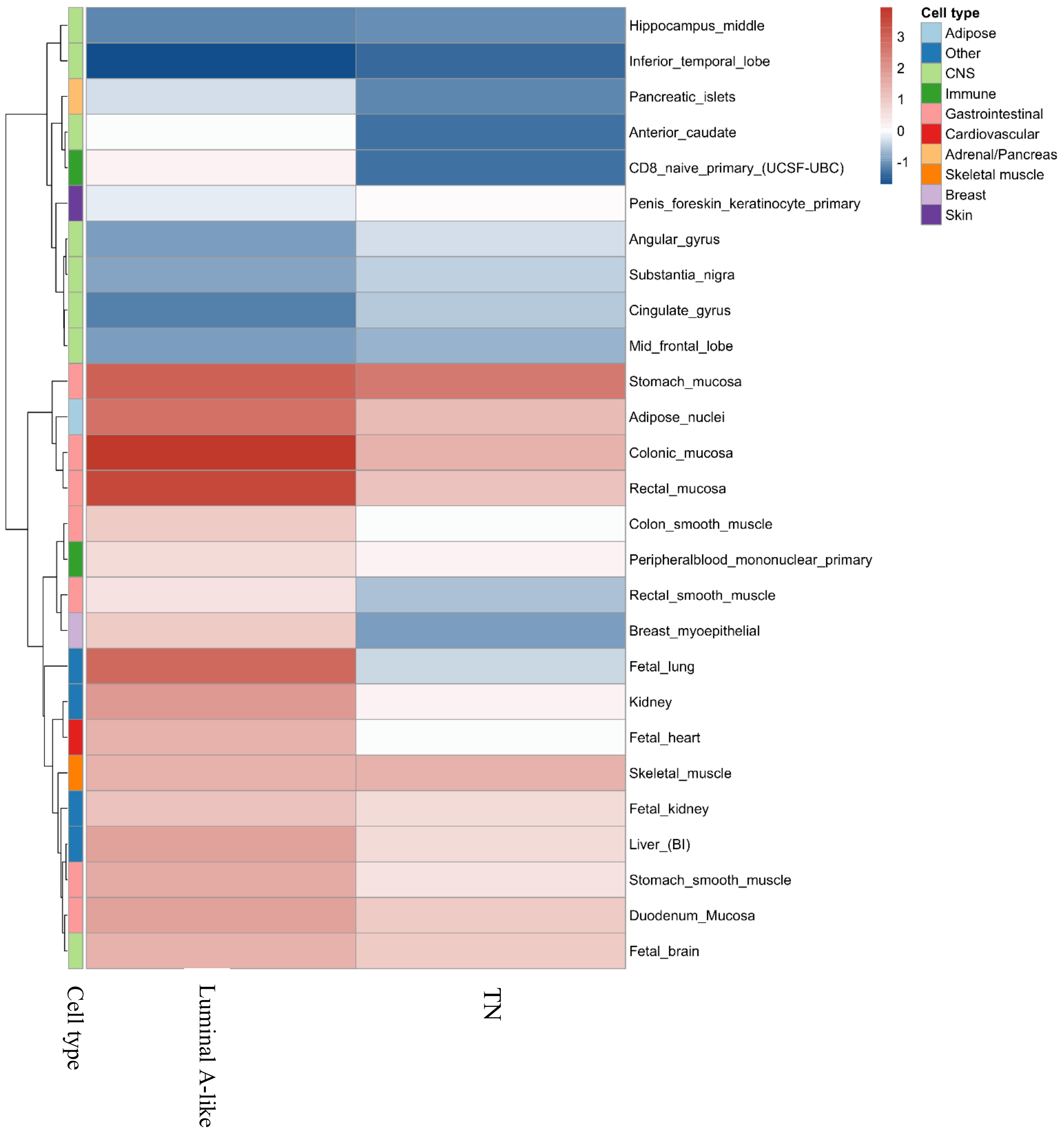

**Supplementary Note**

The two-stage polytomous logistic regression model allows us to efficiently test for genetic associations while accounting for tumor marker correlations and large amounts of missing tumor data [1]. We used this method to detect breast cancer susceptibility SNPs while taking account of four tumor characteristics: estrogen receptor (ER; ER-positive vs ER-negative), progesterone receptor (PR; PR-positive vs PR-negative), human epidermal growth factor receptor 2 (HER2; HER2-positive vs HER2-negative), and grade (defined as grade 1, grade 2, and grade 3). Below we describe in greater detail how we applied this method

### Two-stage polytomous model

In our study, we investigated for underlying heterogenous associations according to ER, PR, HER2, and grade; however, we will first start the discussion of fitting a two-stage polytomous model by first focusing on ER, PR, and HER2, and then discuss including grade in the model. The cross combination of ER, PR, and HER2 results in eight distinct breast cancer subtypes (8 = 2x2x2). Let N denote the total sample size and let $\mathrm{D_i}$ denote the disease status of ith subject which can take values from $\{0,1,2, \dots ,8\}$ and $\mathrm{i} = 1,2, \dots, \mathrm{N}$. $\mathrm{D_i} = 0$ represent a control, and $\mathrm{D_i} = m$ represent the ith subject with the breast cancer subtypes M. Let $\mathrm{G_i}$ denote the genotype of a SNP for ith subject, taking values from $\{0,1,2\}$. Let $\mathbf{X_i}$ denote the other covariates for the ith subject, for example principal components or age. In the first stage of the model, we fit a standard "saturated" polytomous logistic regression model:

$$\Pr(\mathrm{D_i} = \mathrm{m} | \mathrm{G_i}, \mathrm{X_i}) = \frac{\exp\left(\beta_m G_i + \boldsymbol{\eta}_{\boldsymbol{m}}^{\boldsymbol{T}} \boldsymbol{X_i}\right)}{1 + \sum_{m=1}^{8} \exp\left(\beta_m G_i + \boldsymbol{\eta}_{\boldsymbol{m}}^{\boldsymbol{T}} \boldsymbol{X_i}\right)}, \quad (1)$$

where $\beta_{\mathrm{m}}$ is the regression coefficient for SNP (G) associated with the mth subtype and $\eta_m$ is the vector of regression coefficients for the other covariate (X) associated with mth subtype.

Each cancer subtype m is defined through a unique combination of ER, PR, and HER2; therefore, we can alternatively index the parameters $\beta_{\mathrm{m}}$ as $\beta_{s_1 s_2 s_3}$, where $s_1, s_2, s_3 \in \{0, 1\}$ for the three binary tumor characteristics. Originally, $\beta_1$ represented the regression coefficient of the ER-, PR-, HER2- subtype. With this indexing, $\beta_1$ can be alternatively written as $\beta_{000}$ and, thus with this reparameterization we can represent the log odds ratio of the eight subtypes as:

$$\beta_{s_1 s_2 s_3} = \theta^{(0)} + \theta_1^{(1)} s_1 + \theta_2^{(1)} s_2 + \theta_3^{(1)} s_3 + \theta_{12}^{(2)} s_1 s_2 + \theta_{13}^{(2)} s_1 s_3 + \theta_{23}^{(2)} s_1 s_3 + \theta_{123}^{(3)} s_1 s_2 s_3, \quad (2)$$

where $\theta_0^{(0)}$ represents the case-control log odds ratio for a reference subtypes versus the controls. We have chosen ER-, PR-, HER2- as the reference subtype, but any subtype can be chosen as the reference subtype. $\theta_{\mathrm{k}}^{(1)}$ represents the case-case log odds ratio for the kth tumor characteristic after adjusting for the other tumor characteristics. We also refer $\theta_{\mathrm{k}}^{(1)}$ as the main effect of the kth tumor characteristic. $\theta_{\mathrm{k_1 k_2}}^{(2)}$ represents how the case-case log odds ratio associated with $\mathrm{k_1}$th tumor characteristic is modified by levels of the $\mathrm{k_2}$th tumor characteristic and vice versa. We also refer to $\theta_{\mathrm{k_1 k_2}}^{(2)}$ as the pairwise interaction between the $\mathrm{k_1}$th tumor characteristic and the $\mathrm{k_2}$th tumor characteristic. $\theta_{123}^{(3)}$ represents the third order interaction of the three tumor characteristics. This decomposition is equivalent to the first stage polytomous logistic regression since both the first stage and second stage have eight parameters. We can specify different two stage models by assuming different second stage parameters to be equal to 0. For example, the baseline two-stage model is represented by:

$$\beta_{s_1 s_2 s_3} = \theta^{(0)}. \tag{3}$$

This baseline model assumes all of the subtypes have the same log odds ratio and is equivalent to a standard case-control logistic regression testing the association between an exposure and breast cancer, irrespective of tumor subtypes. We can also constrain all of the second stage pairwise interactions and higher order interactions to be 0:

$$\beta_{s_1 s_2 s_3} = \theta^{(0)} + \theta_1^{(1)} s_1 + \theta_2^{(1)} s_2 + \theta_3^{(1)} s_3. \tag{4}$$

This additive two-stage model assumes the case-case log odds ratio of a tumor characteristic are not affected by interactions with the other tumor characteristics.

By adding the second stage pairwise interactions parameters into the model, we can also construct the pairwise interaction two-stage polytomous model:

$$\beta_{s_1 s_2 s_3} = \theta^{(0)} + \theta_1^{(1)} s_1 + \theta_2^{(1)} s_2 + \theta_3^{(1)} s_3 + \theta_{12}^{(2)} s_1 s_2 + \theta_{13}^{(2)} s_1 s_3 + \theta_{23}^{(2)} s_1 s_3. \tag{5}$$

This model evaluates how two tumor characteristics are modified by each other. For example, $\theta_{12}^{(2)}$ measures how the case-case log odds ratio associated of ER is modified by the status of PR and vice versa. If we further add the three-way interaction term between ER, PR, and HER2, then this model becomes saturated (as shown in in Equation 2) and is equivalent to the polytomous logistic regression.

When we add the three-level ordinal variable tumor grade into the model, we can define 24 (2x2x2x3) breast cancer subtypes. We can apply the same decomposition as implemented with three tumor characteristics to provide the following additive two-stage model:

$$\beta_{s_1 s_2 s_3 s_4} = \theta^{(0)} + \theta_1^{(1)} s_1 + \theta_2^{(1)} s_2 + \theta_3^{(1)} s_3 + \theta_4^{(1)} s_4, \tag{6}$$

where $\theta_4^{(1)}$ is the main effect of grade and $s_4$ can take the values from $\{1, 2, 3\}$. In this model, we assume the grade main effect linearly changes, meaning the average log odds ratios difference between grade 3 versus grade2 is the same the as the difference between grade 2 versus grade1. We can always describe the link between the first stage parameters and second stage parameters in Equation (6) in matrix form:

$$\begin{matrix} \text{ER} - \text{PR} - \text{HER2} - \text{grade1} \\ \text{ER} + \text{PR} - \text{HER2} - \text{grade1} \\ \text{ER} - \text{PR} + \text{HER2} - \text{grade1} \\ \text{ER} + \text{PR} + \text{HER2} - \text{grade1} \\ \text{ER} - \text{PR} - \text{HER2} + \text{grade1} \\ \text{ER} + \text{PR} - \text{HER2} + \text{grade1} \\ \text{ER} - \text{PR} + \text{HER2} + \text{grade1} \\ \text{ER} + \text{PR} + \text{HER2} + \text{grade1} \\ \dots \\ \text{ER} + \text{PR} + \text{HER2} + \text{grade3} \end{matrix} \quad \boldsymbol{\beta} = \begin{bmatrix} \beta_1 \\ \beta_2 \\ \beta_3 \\ \beta_4 \\ \beta_5 \\ \beta_6 \\ \beta_7 \\ \beta_8 \\ \dots \\ \beta_{24} \end{bmatrix} = \begin{bmatrix} 1 & 0 & 0 & 0 & 1 \\ 1 & 1 & 0 & 0 & 1 \\ 1 & 0 & 1 & 0 & 1 \\ 1 & 1 & 1 & 0 & 1 \\ 1 & 0 & 0 & 1 & 1 \\ 1 & 1 & 0 & 1 & 1 \\ 1 & 0 & 1 & 1 & 1 \\ 1 & 1 & 1 & 1 & 1 \\ \dots & \dots & \dots & \dots & \dots \\ 1 & 1 & 1 & 1 & 3 \end{bmatrix} \begin{bmatrix} \theta^{(0)} \\ \theta_1^{(1)} \\ \theta_2^{(1)} \\ \theta_3^{(1)} \\ \theta_4^{(1)} \end{bmatrix} = \mathbf{Z} \begin{bmatrix} \theta^{(0)} \\ \boldsymbol{\theta^H} \end{bmatrix} = \mathbf{Z\theta}, \tag{7}$$

where $\boldsymbol{\beta}$ is a vector of regression coefficients of the first stage parameters, $\boldsymbol{\theta}$ is the vector of all the second stage parameters, and $\boldsymbol{\theta^H}$ is a vector of second stage main effects.

### Hypothesis testing of two-stage polytomous logistic regression

Under the two-stage model framework, there are three different tests we can construct. The first is the global association test:

$$H_0: \theta^{(0)} = 0 \; and \; \boldsymbol{\theta^H} = \mathbf{0} \; versus \; H_1: either \; \theta^{(0)} \neq 0 \text{ or } \boldsymbol{\theta^H} \neq \mathbf{0}\,. \tag{8}$$

This test is designed to test whether a SNP is associated with any of the 24 breast cancer subtypes. If the null hypothesis is rejected under this setting, then at least one of the first stage subtype case-control log odds ratios $\beta_m$ is significantly not equal to 0.

The second test is the global heterogeneity test:

$$H_0: \boldsymbol{\theta^H} = \mathbf{0} \; versus \; H_1: \boldsymbol{\theta^H} \neq \mathbf{0}\,. \tag{8}$$

This test is designed to test whether the associations between a SNP and any two breast cancer subtypes are significantly different from each other. If the null hypothesis is rejected under this setting, then we can conclude that at least two of the first stage subtypes case-control log odds ratios are significantly different with each other ($\beta_{m_1} \neq \beta_{m_2}$).

If the global heterogeneity test is significant, then we can construct the third hypothesis tests, the specific tumor marker heterogeneity test:

$$H_0: \boldsymbol{\theta}^{\boldsymbol{H}}_{(k)} = 0 \; versus \; H_1: \boldsymbol{\theta}^{\boldsymbol{H}}_{(k)} \neq 0\,. \tag{9}$$

This test is designed to test which tumor character is the source of the observed heterogeneity in the global heterogeneity test. Under the additive two-stage model in Equation (6), for example, we can test $H_0: \theta_1^{(1)} = 0 \; versus \; H_0: \theta_1^{(1)} \neq 0$ . This is designed to test whether the case-case log odds ratio of ER is significant not equaling to 0 after adjusting for the effects of PR, HER2 and grade.

#### Mixed effect two-stage polytomous model

Although the additive two-stage model decreases the degrees of freedoms compared to the first stage polytomous logistic regression, the degrees of freedom of the two-stage model are still penalized when additional tumor characteristics are included into the model. To address this issue, we developed the mixed effect two-stage polytomous model to enter tumor characteristic variables into the model as either fixed- or random-effect terms. In this model, we keep the second stage main effect of ER ($\theta_1^{(1)}$) as a fixed effect since there is strong *a priori* evidence that ER is a common source of heterogeneity [2]. On the other hand, as there is minimal evidence suggesting that tumor characteristics such as PR, HER2, and grade are sources of heterogeneity, we assume the case-case parameter of PR ($\theta_2^{(1)}$), HER2 ($\theta_3^{(1)}$) and grade ($\theta_4^{(1)}$) as random effects. These random parameters have an assumed arbitrary distribution with mean 0 and variance $\sigma^2$. We always keep the baseline effect $\theta^{(0)}$ as fixed since it captures the overall association between a SNP and breast cancer. Under the mixed effect two stage model, the global test for association is:

$$H_0: \theta^{(0)} = 0, \theta_1^{(1)} = 0, \sigma^2 = 0 \; versus \; H_1: either \; \theta^{(0)}, \theta_1^{(1)}, or \; \sigma^2 \neq 0 \tag{10}$$

The rejection of the null hypothesis implies that the SNP is significantly associated with at least one of the 24 breast cancer subtypes. The global heterogeneity test under the mixed effect two-stage model would be:

$$H_0: \theta_1^{(1)} = 0 \; and \; \sigma^2 = 0 \; versus \; H_1: either \; \theta_1^{(1)} \text{or } \sigma^2 \neq 0 \, . \tag{11}$$

The rejection of the null hypothesis would imply that the SNP's associations between at least two breast cancer subtypes are significantly different. However, the specific tumor marker heterogeneity test for a specific tumor marker is not applied in the mixed effect two-stage model because it requires the estimate of case-case log odds ratio of PR, HER2 and grade which are note estimated when modeled as random effects.

### Two-stage model for intrinsic subtypes of breast cancer

In previous sections, we showed how the first stage case control log odds ratios of breast cancer subtypes are decomposed to the case control log odds ratio of a reference subtype and the into case-case parameters of tumor characteristics. Using the hierarchical second stage decomposition, the two-stage model can also estimate the case control log odds ratio of specific breast cancer subtypes of interest. In our study we defined five intrinsic-like breast cancer subtypes based on tumor status of ER, PR, HER2 and grade: (1) luminal A-like (ER+ and/or PR+, HER2-, grade 1 & 2); (2) luminal B/HER2-negative-like (ER+ and/or PR+, HER2-, grade 3); (3) luminal B-like (ER+ and/or PR+, HER2+); (4) HER2-enriched-like (ER- and PR-, HER2+), and (5) triple negative (TN; ER-, PR-, HER2-). To estimate the case-control log odds ratios of these five intrinsic subtypes we can construct the two-stage model as:

$$\begin{matrix} \text{ER} - \text{PR} - \text{HER2} - \text{grade1} \\ \text{ER} + \text{PR} - \text{HER2} - \text{grade1} \\ \text{ER} - \text{PR} + \text{HER2} - \text{grade1} \\ \text{ER} + \text{PR} + \text{HER2} - \text{grade1} \\ \text{ER} - \text{PR} - \text{HER2} + \text{grade1} \\ \text{ER} + \text{PR} - \text{HER2} + \text{grade1} \\ \text{ER} - \text{PR} + \text{HER2} + \text{grade1} \\ \text{ER} + \text{PR} + \text{HER2} + \text{grade1} \\ \cdots \\ \text{ER} + \text{PR} + \text{HER2} + \text{grade3} \end{matrix} \quad \boldsymbol{\beta} = \begin{bmatrix} \beta_1 \\ \beta_2 \\ \beta_3 \\ \beta_4 \\ \beta_5 \\ \beta_6 \\ \beta_7 \\ \beta_8 \\ \cdots \\ \beta_{24} \end{bmatrix} = \begin{bmatrix} 0 & 0 & 0 & 0 & 1 \\ 1 & 0 & 0 & 0 & 0 \\ 1 & 0 & 0 & 0 & 0 \\ 1 & 0 & 0 & 0 & 0 \\ 0 & 0 & 0 & 1 & 0 \\ 0 & 1 & 0 & 0 & 0 \\ 0 & 1 & 0 & 0 & 0 \\ 0 & 1 & 0 & 0 & 0 \\ \cdots & \cdots & \cdots & \cdots & \cdots \\ 0 & 1 & 0 & 0 & 0 \end{bmatrix} \begin{bmatrix} \theta_1 \\ \theta_2 \\ \theta_3 \\ \theta_4 \\ \theta_5 \end{bmatrix} \quad \begin{matrix} \text{Luminal A} - \text{like, low grade} \\ \text{Luminal B} - \text{like} \\ \text{Luminal B/HER2} - \text{negative} - \text{like} \\ \text{HER2 enriched} - \text{like} \\ \text{Triple negative} \end{matrix} \tag{12}$$

Under this model, the second stage parameters provide estimates of case-control log odds ratios for the five tumor subtypes. This model is similar to directly fitting a polytomous logistic regression. However, we have incorporated into the two-stage model an efficient missing data algorithm that allows to take advantage of subjects with incomplete tumor characteristic data. The missing data algorithm has been described in detail elsewhere [1].

### Modified LD score regression

Since the two-stage polytomous logistic regression implements an EM algorithm to account for missing tumor characteristics data, the effective sample size is not equivalent to the sample size of cases with complete tumor characteristic data. In this case the sample size is not available, but the log odds ratio for each SNP $\hat{\beta}_j$ and the standard error $s_j$ are given.

Under a case-control study, we consider the logistic regression model

$$log\left(\frac{p}{1-p}\right) = \alpha + \left(\boldsymbol{\beta}^{(J)}\right)^{\boldsymbol{T}} \boldsymbol{X},$$

where $\boldsymbol{\beta}^{(J)} = (\beta_1^{(J)}, \beta_2^{(J)}, \dots, \beta_M^{(J)})$ are the joint effect sizes. We define the heritability as $h^2 = var((\boldsymbol{\beta}^{(J)})^T \boldsymbol{X})$, assuming X is standardized with mean 0 variance 1. If X is in the original 0, 1, 2 scale, we multiply the $\hat{\beta}_j$ and $s_j$ by $\sqrt{2p_j(1-p_j)}$ to standardize, where $p_j$ is the minor allele frequency for the jth SNP. Therefore, the expected chi-square statistics ($z_j^2$) of SNP j is

$$\begin{aligned} \mathrm{E}(z_j^2|l_j) &= \frac{E(\hat{\beta}_j^2|l_j)}{s_j^2} = \frac{\left[E\left\{(\hat{\beta}_j - \beta_j)^2 \middle| l_j\right\} + 2E[(\hat{\beta}_j - \beta_j)\beta_j|l_j] + E(\beta_j^2|l_j)\right]}{s_j^2} \\ &= \frac{\left[E\left\{(\hat{\beta}_j - \beta_j)^2 \middle| l_j\right\} + E(\beta_j^2|l_j)\right]}{s_j^2} \\ &= 1 + \frac{E\{(\sum_k r_{jk}\beta_k^{(J)})^2\}}{s_j^2} \\ &= 1 + \frac{h^2}{M}\frac{l_j}{s_j^2}, \end{aligned} \tag{13}$$

where $l_j = \sum_k r_{jk}^2$ is the LD score of the SNP j and $1/s_j^2$ is the effective sample size for SNP j. The modified LD score regression formula is:

$$\mathrm{E}(z_j^2|l_j) = 1 + \frac{h^2}{M}\frac{l_j}{s_j^2}. \tag{14}$$

To estimate the genetic correlation between two traits, the expected value of $z_{1j}z_{2j}$ for a SNP j is

$$\begin{aligned} \mathrm{E}(\mathrm{z}_{1\mathrm{j}}z_{2j}|l_j) &= \frac{E(\hat{\beta}_{1j}\,\hat{\beta}_{2j}|l_j)}{s_{1j}s_{2j}} \\ &= \frac{\left[E\{(\hat{\beta}_{1j} - \beta_{1j})(\hat{\beta}_{2j} - \beta_{2j})|l_j\} + E(\beta_{1j}\beta_{2j}|l_j)\right]}{s_{1j}s_{2j}} \\ &= \frac{s_{12j}}{s_{1j}s_{2j}} + \frac{E(\sum_k r_{jk}\beta_{1k}^{(J)} \sum_k r_{jk}\beta_{2k}^{(J)}\,|l_j)}{s_{1j}s_{2j}} \\ &= \frac{s_{12j}}{s_{1j}s_{2j}} + \frac{\rho_g}{M}\frac{l_j}{s_{1j}s_{2j}}, \end{aligned} \tag{15}$$

where $\rho_g$ is the genetic covariance between the two different traits. Under this case, $1/s_{1j}^2$ and $1/s_{2j}^2$ are the effective sample size for SNP j for the two traits respectively. The modified LD score regression for genetic covariance is

$$\mathrm{E}(\mathrm{z}_{1\mathrm{j}}z_{2j}|l_j) = \frac{s_{12j}}{s_{1j}s_{2j}} + \frac{\rho_g}{M}\frac{l_j}{s_{1j}s_{2j}}. \tag{16}$$

The genetic correlation is given by $\frac{\rho_g}{\sqrt{h_1^2 h_2^2}}$.

**Effective sample size of cases of two-stage polytomous model**

The two-stage polytomous model implements the EM algorithm to impute missing tumor characteristics; therefore, the effective sample size of cases is not equivalent to the actual number of cases with available tumor characteristic data. We estimated the effective sample sizes to help demonstrate the benefit of using the EM algorithm to impute missing tumor characteristics and to aid comparability with previous studies (Supplementary Table 4). To estimate the effective sample size, suppose we have a complete dataset with no missing tumor characteristics, the sample size is $n_k$ for the kth subtype and $n_0$ for the control. If we fit a two-stage polytomous model for the jth SNP, the corresponding log odds ratio for kth subtype is $\hat{\beta}_{jk}$ and the standard error is $s_{jk}$. Then, approximately:

$$var(\hat{\beta}_{jk}|p_j) \approx \frac{n_0 + n_k}{2 * p_j(1 - p_j)(n_0 n_k)},$$

where $p_j$ is the MAF of the jth SNP. Now we consider fitting a two-stage polytomous model with missing tumor characteristics. Given the standard error $s_{jk}$ of the log odds ratio and the control sample size, we have the estimate of effective number of cases as,

$$\hat{n}_k = \left(\frac{1}{n_0} - 2s_{jk}^2 p_j(1 - p_j)\right)^{-1}.$$

We used the median estimates of effective sample size of cases for all SNPs as the final estimate.
